# Optimizing twin prime editing components for scalable genome editing and therapy in spinocerebellar ataxia type 3

**DOI:** 10.1101/2025.07.22.666117

**Authors:** Lee Wha Gwon, Jung Bae Seong, Hyeon-Gu Yeo, Yeounsun Oh, Junghyung Park, Jinyoung Won, Sang Je Park, Young-Hyun Kim, Jae-won Huh, Aryun Kim, Youngjeon Lee, Seung Hwan Lee

**Affiliations:** Department of Life Science, Chung-Ang University, Seoul 06974, Republic of Korea; National Primate Research Center (NPRC), Korea Research Institute of Bioscience and Biotechnology (KRIBB), Cheongju 28116, Republic of Korea; KRIBB School of Bioscience, University of Science and Technology (UST), Daejeon 34113, Republic of Korea; Department of Functional Genomics, KRIBB School of Bioscience, Korea University of Science & Technology (UST), Daejeon 34113, Republic of Korea; Department of Neurology, College of Medicine, Chungbuk National University, Cheongju, South Korea; Department of Neurology, Chungbuk National University Hospital, Cheongju, South Korea

## Abstract

Recent advances in prime editing technologies using CRISPR modules fused with reverse transcriptase (RT) have enabled efficient and precise reprogramming of target genomic sequences. Twin prime editing using two coordinated prime editor complexes is a promising strategy for inducing extensive genomic modifications via reverse transcribed complementary templates. However, current twin prime editing systems still require improvements in editing efficiency, accuracy, and intended edit predictability. Here, efficiency and precision of twin prime editing were enhanced via engineering and optimizing conventional SpCas9(H840A)-RT-based prime editor components. A La domain–fused prime editor (La-SpCas9(H840A)-RT) and optimized pegRNAs were developed, achieving a 1.75 ± 0.21-fold increase in gene editing efficiency at multiple genomic loci in human-derived cell lines without increasing off-target activity. La-SpCas9(H840A)-RT facilitated efficient ∼2.8 kb GFP transgene knock-in at target loci and eliminated the expanded polyQ tract in *ATXN3* in patient-derived mutant cell lines modeling spinocerebellar ataxia type 3. The advanced twin prime editing platform expands genome engineering capabilities beyond existing CRISPR-based systems and holds great promise for diverse biotechnological and therapeutic applications.

## Introduction

CRISPR technology enables targeted DNA editing based on reprogrammed guide RNAs^1,2^. It has rapidly evolved into advanced forms enabling high target specific gene editing, with prime editing-based universal single-base gene editing^3–6^ to large-scale DNA editing^7–9^. Prime editing technology fuses a reverse transcriptase (RT) to a CRISPR-Cas9 module targeting specific DNA to induce versatile gene editing using a target DNA-matching pegRNA ^4,10^. Recently, engineered versions of the prime editing module, PE2 or PE3, incorporate function-specific domains, to improve gene editing accuracy and efficiency^11–14^. A major limitation of this prime editing technology lies in difficulty integrating long edited DNA flaps into target sites after RT copies pegRNA, making it challenging to edit regions longer than a few base-pairs^5^. This twin prime editing approach uses two prime editor modules to create extended complementary DNA flaps for inducing deletion, replacement, or insertion, enabling efficient large-scale DNA editing that a single prime editor cannot handle^15–19^. Twin prime editing currently shows minimal off-target editing and is an effective gene editing strategy, particularly when combined with other recombinases^7–9^. This aimed to improve twin prime editing efficiency and precision by optimizing pegRNAs, components of the prime editor, and attaching the functional La protein domain to enable high-efficiency large-scale editing. In this study, we demonstrate that the twin prime editing strategy, utilizing fully-matched pegRNAs developed herein and the La-SpCas9(H840A)-RT module, achieves superior genome editing efficiency and precision compared to conventional systems employing SpCas9(H840A)-RT. Across multiple genomic loci in human-derived cell lines, this optimized twin prime editing system exhibited significantly enhanced editing efficiency while minimizing unintended insertions and deletions (indels), thereby improving editing fidelity. Notably, sequential application of twin prime editing and the Bxb1 recombinase enabled precise insertion of large gene cassettes. These findings collectively establish a robust and accurate genome engineering framework capable of inducing large-scale genetic modifications in human cells, thereby advancing the translational potential of this platform for therapeutic applications in vivo. The proof-of-concept study in the SCA3 model suggests this platform could make a substantial contribution to gene therapy, particularly in the correction of complex and extensive genetic mutations—such as those found in hemophilia^20–22^ and Huntington’s disease^23,24^—that are challenging to address using conventional genome editing technologies.

## Results

### Improvement of twin prime editing efficiency in human-derived cell lines via optimization of prime editor components and targeting strategies

In this study, various strategic approaches were explored to improve twin prime editing efficiency and precision **(Figure 1)**. First, protein-level engineering was introduced to enhance the efficiency of each prime editor **(Figure 1a)**. Conventional prime editors^4,25^ were constructed by fusing a reverse transcriptase (RT) to SpCas9 in its nickase (H840A) or wild-type form, resulting in SpCas9(H840A)-RT and SpCas9(WT)-RT. Here, engineered La-SpCas9(H840A)-RT and La-SpCas9(WT)-RT were developed via La domain fusion^13^, and these four protein module combinations were used to induce twin prime editing **(Supplementary Figure S1)**. Second, pegRNA PBS and RTT region length and complementarity between the target DNA and pegRNA pair were adjusted to induce optimized twin prime editing outcomes **(Figure 1b)**. Twin prime editing at the three genomic loci (*HEK3, FANCF*, and *AAVS1*) in human-derived cell lines (HEK293FT) **(Supplementary Figure S2)** yielded highly consistent editing results. Notably, twin prime editing with SpCas9(H840A)-RT showed higher accuracy in inserting *attB* or *attP* sequences into target DNA than with SpCas9(WT)-RT **(Figure 1b, Supplementary Figure S3, S4)**. Gene editing patterns included complete insertion of *attB* or *attP* sequences (precise Twin-PE) and partial insertion of *attB*/*attP* sequences or complete indels **(Supplementary Figure S3)**. Partial insertion and complete indel rates were higher when twin prime editing was induced with SpCas9(WT)-RT than with SpCas9(H840A)-RT **(Figure 1b, Supplementary Figure S3, S4)**. When designing pegRNAs for each twin prime editor, three distinct types were employed: fully complementary (Whole overlap), homology arm-forming (Homology arm), and partially complementary (Partial overlap) **(Figure 1b, Supplementary Figure S4, left)**. Across all target loci, pegRNA designs based on either the ‘Whole overlap’ or ‘Partial overlap’ strategy consistently yielded higher-fidelity twin prime editing than the ‘Homology arm’ design, regardless of whether SpCas9(H840A)-RT or SpCas9(WT)-RT was used **(Figure 1b, Supplementary Figure S4, right)**. Notably, when twin prime editing was induced using the SpCas9(WT)-RT module in combination with pegRNAs, whose complementary binding regions formed homology arms with the target DNA, a substantial portion of the editing outcomes consisted of precisely generated indels **(Figure 1b, Supplementary Figure S4, middle row)**. Among all twin prime editor combinations, precise editing efficiency significantly increased (1.40 ± 0.26-fold increase) when La domain–fused La-SpCas9(H840A)-RT or La-SpCas9(WT)-RT modules were designed with fully or partially complementary and pegRNAs **(Figure 1b, top and bottom row)**. However, no increase was observed in any of the three genomic sites (*HEK3*, *FANCF*, and *AAVS1*) in designs forming homology arms at the same target loci **(Figure 1b, middle row)**. When different sequences (*attB* or *attP*) were inserted into the target gene (*AAVS1*), similar editing trends were observed across all four twin prime editing combinations **(Supplementary Figure S4)**. La-SpCas9(H840A)-RT was selected as the optimized twin prime editing module and all subsequent experiments were conducted using pegRNAs designed in the ‘Whole overlap’ configuration, with comparative analyses performed against SpCas9(H840A)-RT.

**Fig. 1.**
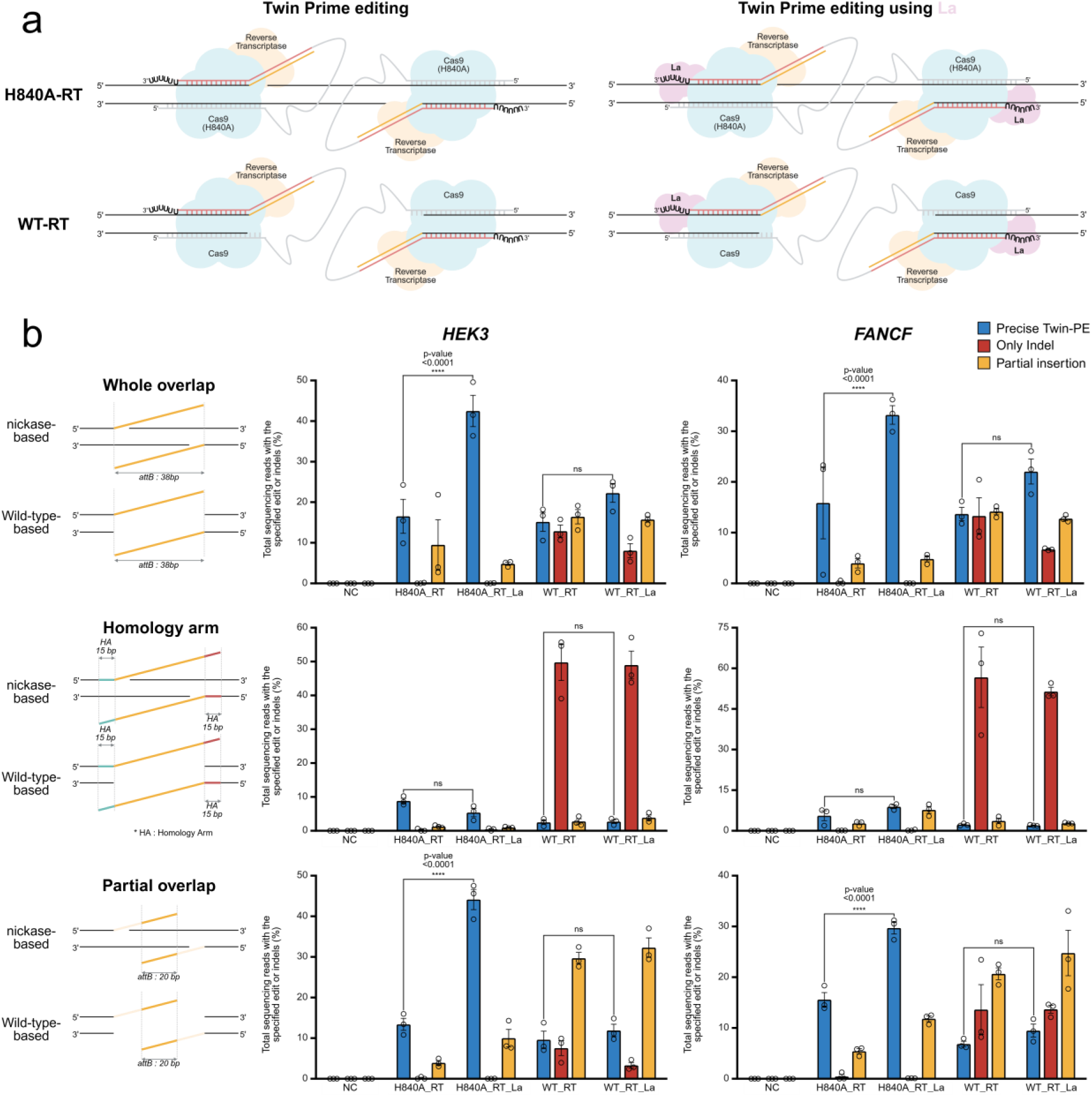
| Optimization of prime editor components and gene targeting strategies for effective twin prime editing in human-derived cells. (a) Schematic of twin prime editing using conventional prime editors based on nickase (H840A) or wild-type (WT) SpCas9 modules, or prime editors engineered with the La domain. RT: reverse transcriptase, La: La domain (1–194 aa), SpCas9(H840A): SpCas9 nickase, SpCas9(WT): SpCas9 wild-type. (b) Comparative experiment assessing the efficiency of 38 bp *attB* sequence insertion using twin prime editing at target gene loci (*HEK3* and *FANCF*) in human-derived cells (HEK293FT). The prime editor was combined with three types of pegRNA overlap strategies in based on whether SpCas9 was nickase (H840A) or wild-type (WT), and whether the La domain (1–194 aa) was fused to SpCas9. SpCas9(H840A/WT)-RT: prime editor based on the nickase (H840A) or wild-type (WT) SpCas9 module; La-SpCas9(H840A/WT)-RT: prime editor based on the nickase (H840A) or wild-type (WT) SpCas9 module fused with La domain; Whole overlap pegRNA pair: the entire sequence to be inserted into target DNA is fully complementary to both pegRNAs; Homology arm pegRNA pair: both pegRNAs include the sequence to be inserted alongside homology arm regions corresponding to target DNA; Partial overlap pair: only 20 bp of the sequence to be inserted into target DNA is complementarily shared between the two pegRNAs. Twin prime editing efficiency and pattern analysis were classified into three forms: precise Twin-PE (%), Only Indel (%), and Partial Insertion (%), and the frequency (%) of each was measured (Supplementary Figure S3). Each histogram represents the mean ± standard error of the mean (SEM) from three independent measurements. P-values were calculated using two-way ANOVA and Dunnett’s test (ns: not significant, *P = 0.0332, **P = 0.0021, ***P = 0.0002, ****P <0.0001). Precise Twin-PE (%): The proportion of precisely edited alleles reflecting the intended twin prime editing outcome. Only Indel (%): The frequency of unintended editing events resulting solely in insertions or deletions without intended sequence insertion. Partial Insertion (%): The rate of imprecise editing events characterized by incomplete incorporation of the target insertion sequence. NC: negative control; H840A_RT: SpCas9(H840A)-RT; H840A_RT_La: La-SpCas9(H840A)-RT; WT_RT: SpCas9(WT)-RT; WT_RT_La: La-SpCas9(WT)-RT.

### Efficient induction of twin prime editing across multiple genomic loci in human-derived cells

Based on previous experimental results, additional validation experiments were conducted using human-derived cells to induce precise sequence insertion into target DNA via twin prime editing **(Figure 2)**. Twin prime editing using the nickase (H840A) form shows more precise gene editing than that using the wild-type (WT) form **(Figure 1, Supplementary Figure S4)**. Therefore, we compared twin prime editing efficiency and precision at 13 genomic loci using the conventional SpCas9(H840A)-RT and the engineered, optimized La-SpCas9(H840A)-RT modules **(Figure 2a-k)**. This study showed that using the La-SpCas9(H840A)-RT module causes a significant increase in precise gene editing efficiency **(Figure 2l)**. As expected, unlike twin prime editing based on the wild-type module, the SpCas9(H840A)-RT and La-SpCas9(H840A)-RT modules showed low levels of unintended indel formation. However, a minor rate of partial insertions, presumably owing to the malfunction of a single prime editor, was observed as a byproduct in the twin prime editing outcomes. Overall, the La-SpCas9(H840A)-RT module led to a higher ratio of precise gene insertion to total aberrant outcomes (Precise twin prime editing/Partial insertion + Only indel) across diverse genomic loci in human-derived cells than SpCas9(H840A)-RT **(Figure 2l)**. Specifically, La-SpCas9(H840A)-RT showed no statistically significant increase in unintended indel formation compared to SpCas9(H840A)-RT, whereas precise gene editing efficiency showed significantly increased (average 1.75 ± 0.21-fold) **(Figure 2l, inset)**.

**Fig. 2.**
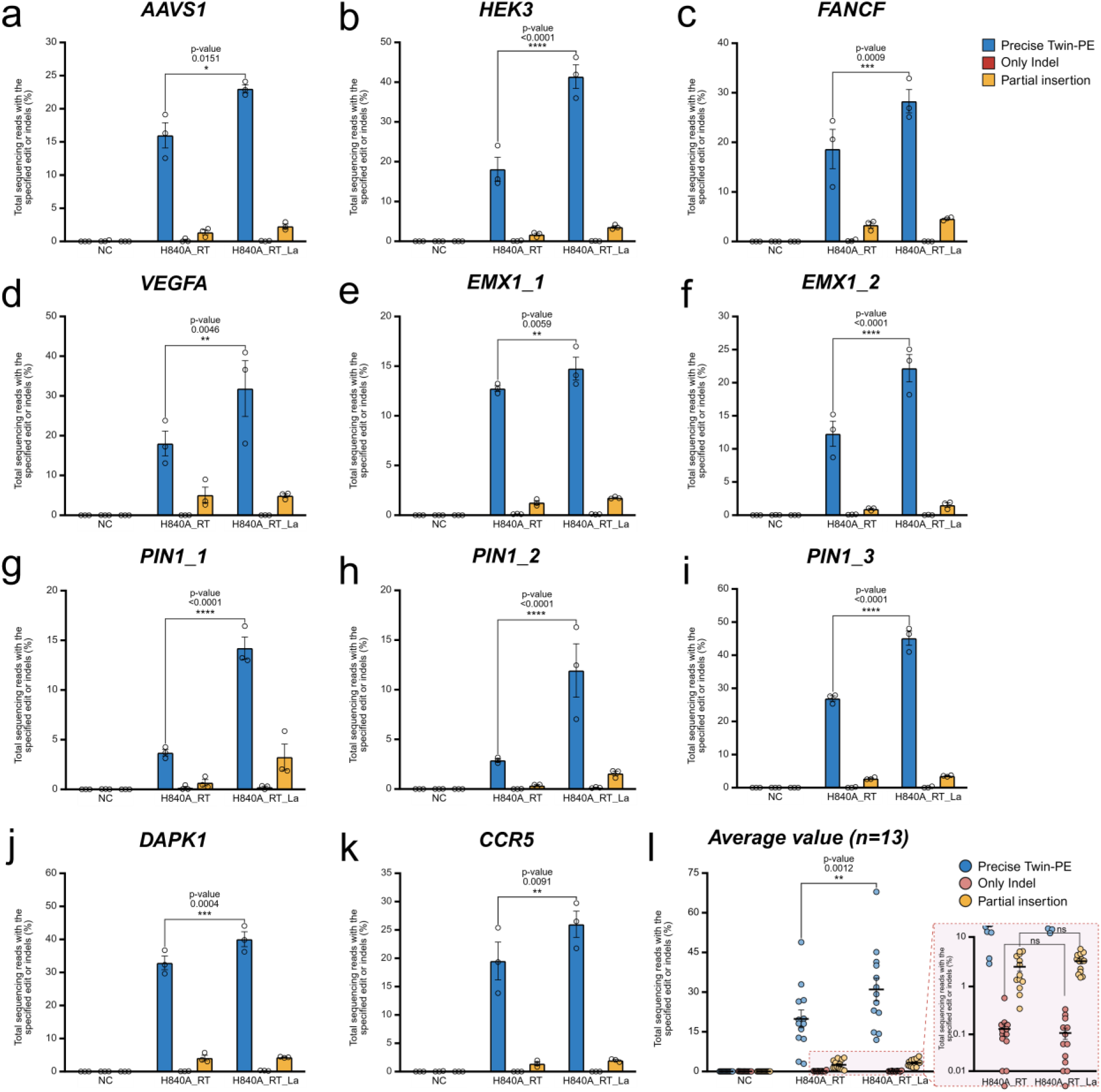
| Verification of optimized twin prime editing efficiency targeting various endogenous genes in human-derived cells. (a–k) Comparison of twin prime editing efficiency (%) based on SpCas9(H840A)-RT or La-SpCas9(H840A)-RT targeting 11 endogenous loci in human-derived cells (HEK293FT). (l) Histogram showing the average twin prime editing efficiency at 13 endogenous loci (n = 13, including the final two loci presented in **Fig. 3**) in human-derived cells. The small inset shows enlarged graphs of “only indel” or “partial insertion” patterns resulting from twin prime editing. Each histogram represents the mean ± standard error of the mean (SEM) from three independent experiments. P-values were calculated using two-way ANOVA and Dunnett’s test (ns: not significant, *P = 0.0332, **P = 0.0021, ***P = 0.0002, ****P <0.0001). NC: negative control; H840A_RT: SpCas9(H840A)-RT; H840A_RT_La: La-SpCas9(H840A)-RT.

### Comparative validation of off-target effects using the optimized twin prime editing system

Next, to compare and analyze the accuracy of La-SpCas9(H840A)-RT-induced twin prime editing with those of the conventional SpCas9(H840A)-RT module, *attB* sequence insertion was examined at various genomic loci (*HEK3, VEGFA, PIN1*) **(Figure 3)**. To simultaneously confirm off-target editing at genomic loci sequence similarity to the targets, targeted amplicon sequencing was conducted on candidate sites obtained via *in silico* prediction^26^. For each pegRNA pair used in twin prime editing, off-target candidates on both the sense **(Figure 3, Left)** and antisense **(Figure 3, Right)** strands were individually detected with high resolution (indel frequency >0.1), allowing up to three mismatches. Subsequently, targeted amplicons were generated for each sample edited with either the optimized La-SpCas9(H840A)-RT module or the conventional SpCas9(H840A)-RT module, followed by NGS-based sequencing and comparative analysis. Consequently, except for a minimal signal (indel frequency ∼0.1) detected at an off-target candidate (off-target 1) corresponding to the *HEK3* antisense pegRNA **(Figure 3, Top, Right)**, no significant off-target editing was observed for any of the pegRNA pairs at the three tested genomic loci. These results indicated that using the optimized La-SpCas9(H840A)-RT module maximizes editing efficiency without increasing off-target effects. Specifically, the twin prime editing mechanism based on complementary pegRNA pairing using the La-SpCas9(H840A)-RT module demonstrated high efficiency and safety, suggesting its potential applicability to human gene therapy.

**Fig. 3.**
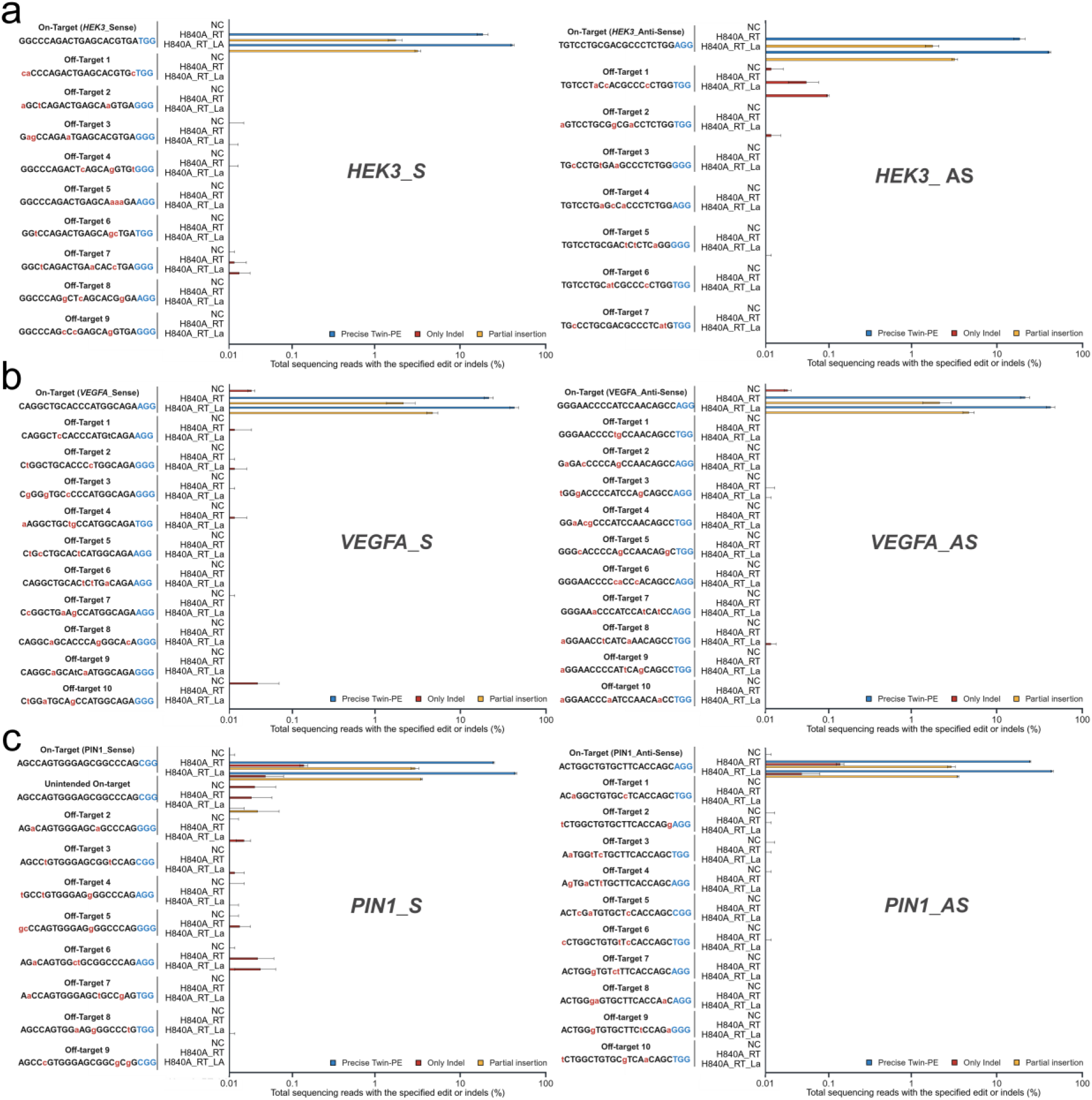
| Off-target effect analysis of optimized twin prime editing in human-derived cells. Twin prime editing used SpCas9(H840A)-RT or La-SpCas9(H840A)-RT at three endogenous gene loci (*HEK3* (a), *VEGFA* (b), *PIN1* (c)) in human-derived cells (HEK293FT). Off-target candidate sequences predicted *in silico* for each pegRNA pair (S-/AS-strand targeting) were analyzed using targeted amplicon sequencing. Sequences listed on the left side of each histogram represent on-target sequences used for twin prime editing and off-target sequences predicted *in silico* based on those on-targets. Blue letters indicate the PAM (NGG) sequence recognized by SpCas9. Red lowercase letters indicate mismatched bases relative to the on-target sequence. S: Sense; AS: Anti-sense; NC: negative control; H840A_RT: SpCas9(H840A)-RT; H840A_RT_La: La-SpCas9(H840A)-RT.

### Induction of large-scale gene insertion using the optimized twin prime editing system

Large-scale genomic regions within target DNA can be edited using twin prime editing^15,19^. Thus, we investigated whether efficient large-scale genome editing could be induced in human-derived cells using the La-SpCas9(H840A)-RT module, which has been validated for both efficiency and safety in this study **(Figure 4)**. In recently developed technologies such as PASSIGE^8^ or PASTE^9^, donor sequence insertion is facilitated via Bxb1 recombinase using *attB* sequences inserted into target genes using single or dual prime editor modules. Consequently, we hypothesized that if high-efficiency *attB* insertion into target DNA is achieved via twin prime editing using the La-SpCas9(H840A)-RT module, the resulting difference in twin prime editing efficiency improves donor sequence insertion via Bxb1 within an effective range. To test this hypothesis, *GFP* insertion efficiency into target DNA using both SpCas9(H840A)-RT and La-SpCas9(H840A)-RT modules was compared **(Figure 4a)**. The donor vector was designed approximately 2.8 kb and included a recombinase recognition sequence (*attP*) along with the GFP coding sequence. Sequential *attB* and donor vector insertion into the target DNA (*GAPDH* locus) via twin prime editing was confirmed feasible **(Supplementary Figure S5a)**. Therefore, the La-SpCas9(H840A)-RT module significantly increased precise *attB* insertion efficiency via twin prime editing compared to SpCas9(H840A)-RT, without substantially changing unintended indel formation **(Figure 4b)**. When two pairs of pegRNAs (set 1, 2) were used to target the *GAPDH* locus via twin prime editing, a similar trend of increased editing efficiency with La-SpCas9(H840A)-RT was observed. Therefore, twin prime editing using two pegRNA pairs was conducted, followed by Bxb1-mediated donor vector insertion **(Figure 4c, d, Supplementary Figure S5b)**. When the second pegRNA pair (Set 2) was used in combination with the La-SpCas9(H840A)-RT module, significant large-scale editing was successfully induced (average 2.98±1.51-fold) **(Figure 4d)**. These results supported the original hypothesis that La-SpCas9(H840A)-RT–based twin prime editing is more effective than SpCas9(H840A)-RT–based editing and following *attB* sequence insertion via twin prime editing enabled efficient recombination with an *attP*-GFP donor vector.

**Fig. 4.**
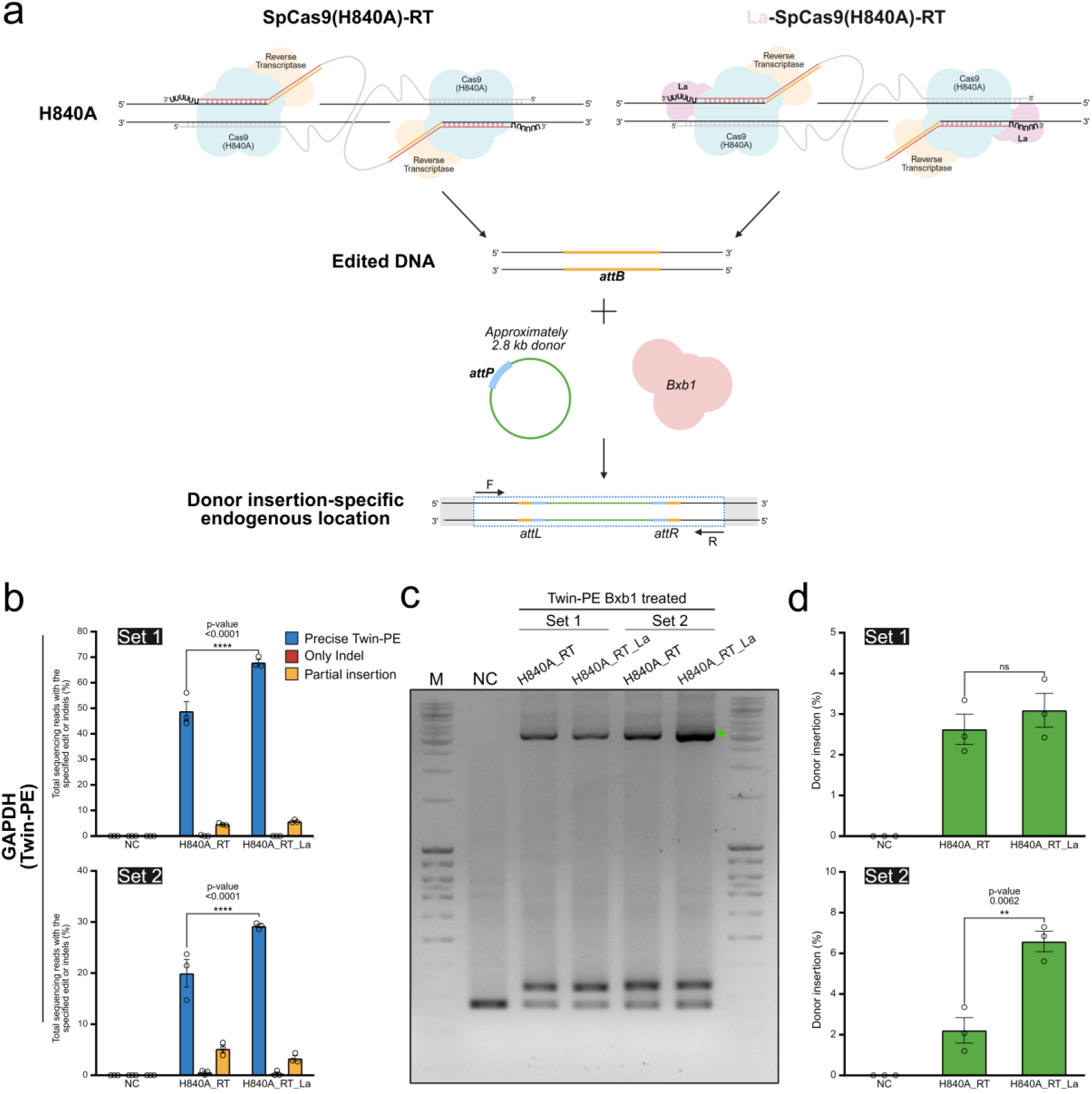
| Verification of large gene insertion efficiency via optimized twin prime editing in human-derived cells. (a) Schematic of large DNA insertion using sequential reactions of twin prime editing and Bxb1 recombinase in human-derived cells (HEK293FT). After inducing *attB* sequence insertion at a genomic locus using SpCas9(H840A)-RT or La-SpCas9(H840A)-RT, donor DNA containing an *attP* sequence was recombined via Bxb1 to insert *GFP*. (b) Comparison of twin prime editing efficiency at the *GAPDH* locus in human-derived cells using SpCas9(H840A)-RT or La-SpCas9(H840A)-RT. Set1 and Set2 represent the respective combinations of pegRNAs used for twin prime editing at the *GAPDH* locus. (c) Comparison of donor DNA insertion efficiency via sequential twin prime editing and Bxb1 recombinase action. Agarose gel image (1.5%) showing PCR results from the *GAPDH* locus in cells treated with each prime editor and the Bxb1 enzyme. Top band (Asterisk): Fragment containing inserted donor DNA. Middle band: Fragment with *attB* insertion only via twin prime editing. Bottom band: Wild-type DNA fragment without editing. (d) Histogram showing donor DNA insertion efficiency quantified via band intensity from (c) (donor DNA insertion (%) = size-normalized top band intensity/size-normalized total intensity). All histograms represent the mean ± SEM from three independent experiments. P-values were calculated by two-tailed unpaired Welch’s t-test (ns: not significant, *P = 0.0332, **P = 0.0021, ***P = 0.0002, ****P <0.0001). Donor insertion: Percentage of donor DNA correctly inserted into the *GAPDH* locus.

### Therapeutic application of the optimized twin prime editing system in SCA3

Finally, to establish a conceptual basis for gene therapy in human disease using the La-SpCas9(H840A)-RT module, a proof-of-concept study targeting Spinocerebellar Ataxia 3 was conducted (SCA3) **(Figure 5)**. Spinocerebellar ataxia is a genetic disorder caused by *ATXN3* mutations and manifests in an autosomal dominant manner^27^. Among various spinocerebellar ataxia types, SCA3 has the highest incidence in humans and is a well-characterized genetic disorder with a clear phenotypic correlation owing to polyQ expansion within a single gene (*ATXN3*)^27,28^. To date, no clinically approved and effective treatments exist for SCA3, though gene editing-based strategies for polyQ deletion and normal-length replacement have gained increasing attention^27^. Accordingly, we aimed to conceptually demonstrate an effective therapeutic strategy for SCA3 by applying twin prime editing using the La-SpCas9(H840A)-RT module developed in this study. The twin prime editing system was designed to remove the polyQ repeat by inducing splicing signal substitution triggered exon skipping and activating an in-frame stop codon in the downstream region of the mutation, enabling the expression of a C-terminally truncated form of the ATXN3 protein (known to exhibit reduced cytotoxicity in human cells)^29^ **(Figure 5a)**. To test this hypothesis, a human-derived mutant HEK293FT-*ATXN3*-Q84 cell line was generated **(Supplementary Figure S6)**. A donor DNA vector containing a polyQ X84 repeat and homology arms was designed to target the polyQ X13 region of exon 10 in *ATXN3* and CRISPR-Cas9 was used to induce polyQ X84 insertion **(Supplementary Figure S6a)**. The cells were subsequently cultured and selected at the single-cell level to isolate clones with precisely inserted polyQ X84 sequences **(Supplementary Figure S6b)**. The isolated single-cell clone, HEK293FT-*ATXN3*-Q84, was validated by detecting expression of a larger ATXN3 protein than the wild-type via western blotting **(Supplementary Figure S6c)** and confirming accurate polyQ X84 insertion in *ATXN3* through TP-PCR **(Supplementary Figure S6d)** and Sanger sequencing **(Supplementary Figure S6e)**. To enable efficient and precise twin prime editing in the HEK293FT-*ATXN3*-Q84 cell line, various combinations of sense– and antisense-pegRNAs targeting the upstream and downstream regions flanking the polyQ X84 sequence in *ATXN3* were tested (S1-AS1 to 5, S2-AS1 to 5), and high-efficiency candidate pairs were selected **(Supplementary Figure S7a)**. Consequently, the S1-AS1 or S1-AS3 combinations showed the highest efficiency in inducing the intended gene editing **(Supplementary Figure S7b)**. Therefore, precise twin prime editing was performed **(Figure 5a-d, Supplementary Figure S8a-c)**. When polyQ-repeat targeted twin prime editing (dual stop codon TAA-*attB*-TAA or TAG-*attB*-TAG insertion for early stop generation) was induced in the HEK293FT-*ATXN3*-Q84 cell line, both constructs achieved effective gene editing. Compared to the SpCas9(H840A)-RT module, La-SpCas9(H840A)-RT–based twin prime editing resulted in a 1.38±0.1-fold average increased in precise gene editing efficiency **(Figure 5d, Supplementary Figure S8b)**. Following efficient twin prime editing at the DNA level, elimination of polyQ repeats was confirmed at both RNA and protein levels in accordance with the newly introduced encoded genetic information. In the La-SpCas9(H840A)-RT treated group of mutant HEK293FT-*ATXN3*-Q84 cells, where editing of the polyQ region was induced, total RNA was extracted, reverse-transcribed into cDNA, and analyzed via sequencing. As expected, exon skipping was precisely induced via removal of the splicing signal at the intron 10–11 junction, leading to direct joining of exon 9 to exon 11 and formation of a premature stop codon **(Supplementary** Figure 5d-g). Furthermore, compared with both untreated mutant and wild-type HEK293FT cells, La-SpCas9(H840A)-RT-treated mutant HEK293FT-*ATXN3*-Q84 cells showed a marked reduction in ATXN3 protein aggregates, confirming effective polyQ repeat elimination at the protein level **(Figure 5e-f)**. These results indicated that La-SpCas9(H840A)-RT-based twin prime editing developed in this study represents an effective and fundamental therapeutic strategy for genetic disorders caused by pathogenic polyQ expansion mutations.

**Fig. 5.**
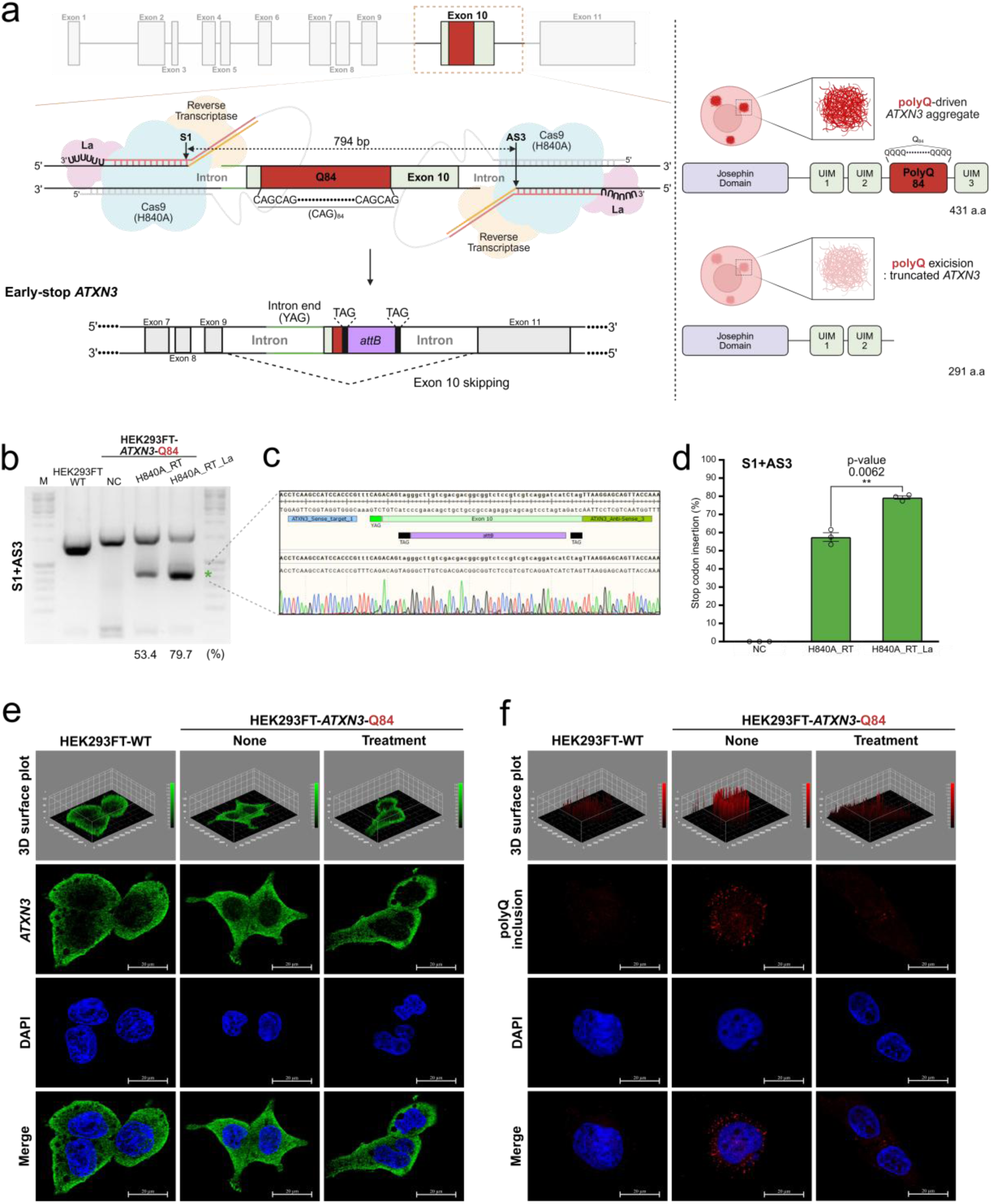
| Validation of a therapeutic concept using optimized twin prime editing targeting the polyQ repeat in SCA3 model cell line. (a) Schematic illustrating the strategy for insertion of an early stop codon (TAA or TAG) and removal of polyQ repeats in *ATXN3* of a human-derived SCA3 disease model cell line using twin prime editing. Left figure shows that each pegRNA pair targets the upstream region (intron between exon 9 and exon 10) and downstream region (intron between exon 10 and exon 11) of polyQ in exon 10 of *ATXN3*, leading to a stop codon insertion upstream of the polyQ sequence and expression of a truncated ATXN3 protein. The figure on the right compares aberrant ATXN3 protein aggregation in the mutant HEK293FT-ATXN3-Q84 cell line with the expression of a truncated ATXN3 protein lacking the polyQ tract, generated via exon 10 skipping induced by twin prime editing. (b) Comparison of twin prime editing efficiency in the *ATXN3* polyQ region of SCA3 model cells using SpCas9(H840A)-RT or La-SpCas9(H840A)-RT. Downward shift of the lower band in the gel indicates successful stop codon insertion. (c) Sanger sequencing confirming correct a stop codon insertion from the lower-shifted bands in (b). (d) Histogram quantifying twin prime editing efficiency, calculated as stop codon insertion (%) = size-normalized bottom band intensity/size-normalized total band intensity. Each histogram represents the mean ± SEM of three independent experiments. P-values were calculated by two-tailed unpaired Welch’s t-test (ns: not significant, *P = 0.0332, **P = 0.0021, ***P = 0.0002, ****P < 0.0001). NC: negative control; H840A_RT: SpCas9(H840A)-RT; H840A_RT_La: La-SpCas9(H840A)-RT. (e-f) Confocal microscopy images obtained using an ATXN3 protein-specific antibody (e) or a polyQ-specific antibody (f) in wild-type HEK293FT cells, mutant HEK293FT-ATXN3-Q84 cells, and mutant HEK293FT-ATXN3-Q84 cells treated with La-SpCas9(H840A)-RT. The first and second rows depict the quantification and intracellular distribution of the ATXN3 protein (e) and polyQ inclusions (f). The third row shows DAPI-stained images of each cell, and the fourth row presents merged images of the second and third rows. None: nothing treated, Treatment: La-SpCas9(H840A)-RT and pegRNA pair was treated for twin prime editing.

## Discussion

In this study, to address the limitations of the conventional SpCas9(H840A)-RT-based twin prime editing system^15^ and improve large-scale genome editing efficiency and accuracy, pegRNA design was optimized within the twin prime editing system and an engineered La-SpCas9(H840A)-RT module was developed, which was comparatively validated across various genomic loci in human-derived cell lines. The conventional twin prime editing system based on dual SpCas9(H840A)-RT modules has been limited by inefficiency and imprecision, primarily due to the inherent complementary ambiguity of the 3’ flap structures generated via RT-mediated pegRNA extension at each gene-specific target site^15,18^. In contrast, the advanced twin prime editing strategy developed in this study overcomes these limitations by (1) employing optimally designed complementary flap structures and (2) utilizing a dual La domain-fused prime editor module, thereby enabling efficient and precise large-scale genome editing across a broad range of genomic loci. Notably, editing outcomes were strongly influenced the degree of complementarity between pegRNAs, regardless of the target sequence type at the editing site. In this study, “whole overlap” and “partial overlap” pegRNA designs demonstrated significantly higher editing efficiencies than those containing extended homology arms. When combined with SpCas9(WT)-RT or La-SpCas9(WT)-RT modules that induce DNA double-strand breaks, these pegRNAs substantially reduced random insertion and deletion (indel) frequency. In twin prime editing experiments comparing the three pegRNA types (whole overlap, homology arm, and partial overlap), the whole overlap design enabled high-efficiency and high-precision editing with both SpCas9(H840A)-RT and SpCas9(WT)-RT modules, demonstrating the importance of pegRNA design tailored to each DNA target. The improved twin prime editing technology using the La domain–fused SpCas9(H840A)-RT module showed enhanced large-scale genome editing efficiency based on pegRNA complementarity compare to the conventional SpCas9(H840A)-RT system. Notably, application of the La-SpCas9(H840A)-RT module to 13 genomic target sites in human-derived cell lines such as HEK293FT resulted in a large fold increase in precise editing efficiency, with unintended indel formation frequency comparable to that of the conventional SpCas9(H840A)-RT module, ensuring predictable gene editing outcomes. The results obtained in this study suggest that the La domain may contribute to the function or stability of the individual prime editor components^13^, improving overall twin prime editing efficiency and highlighting the importance of structural improvements via prime editor engineering for efficient large-scale genome editing. In addition to enhanced editing property, La-SpCas9(H840A)-RT module induced twin prime editing achieved high-precision gene editing without meaningful off-target effects. The improved gene editing precision relative to stochastic DNA misediting events observed in other engineered CRISPR-Cas modules^30–32^ possibly stems from the inherent property of twin prime editing, requiring complementary binding between two pegRNAs to initiate editing^15,16^. Based on the enhanced La-SpCas9(H840A)-RT module, we induced successful insertion of approximately 2.8 kb of GFP into target DNA using La-SpCas9(H840A)-RT–based twin prime editing for efficient *attB* insertion, followed by Bxb1 recombinase–mediated integration. Compared with existing CRISPR-based technologies, this approach yielded greater precision and large-scale gene insertion efficiency, enhancing one-stop genome editing via twin prime editing-recombinase integration. In particular, during large donor vector insertion via Bxb1 following *attB* insertion based on twin prime editing, the La-SpCas9(H840A)-RT module maintained stable insertion efficiency without increasing unintended indel formation compared to the conventional SpCas9(H840A)-RT module, demonstrating superior genome integrity. Furthermore, this study demonstrated the potential of gene therapy using twin prime editing via a proof-of-concept experiment in SCA3 (HEK293FT-*ATXN3*-Q84) cell lines. In conventional SCA3 models, pathogenesis is driven by polyQ repeat expansion in *ATXN3*^27,28^. This study demonstrated precise elimination of the excessively expanded polyQ tract by exon 10 skipping and activating an in-frame stop codon (downstream of the polyQ region in exon 11) to generate truncated form of ATXN3 protein without donor DNA. The La-SpCas9(H840A)-RT module yielded a dramatic fold increase in editing efficiency compared with SpCas9(H840A)-RT and exhibited a high proportion of precise editing. These findings indicated that the optimized La-SpCas9(H840A)-RT-based twin prime editing system can effectively eliminate pathogenic repeat mutation sequences at the DNA level, mitigating mutation-induced cytotoxicity and offering a fundamental therapeutic strategy for genetic disorders caused by single-gene mutations such as SCA3.

A limitation of this study is that all experiments were conducted at the *in vitro* level using human-derived cell lines, leaving potential concerns such as off-target effects, immune responses, or delivery efficiency unresolved for *in vivo* or clinical applications. Additionally, the stability and potential toxicity of the La domain during long-term expression in cells require further investigation. Future research should include *in vivo* validation using animal models, optimizing twin prime editing activity, improving delivery systems, and comprehensive long-term safety assessments of large-scale gene insertion. Collectively, this study presents a new paradigm in large-scale genome editing by structurally improving the efficiency and accuracy of twin prime editing. In particular, pegRNA design optimization and recombinase-based strategies applied to La-SpCas9(H840A)-RT-based twin prime editing demonstrates strong potential as core technology application in genome engineering, disease model development, and gene therapy. These technological advancements are expected to support therapeutic strategies for diseases requiring large-scale gene modifications (e.g., hemophilia, SCA) and facilitate research involving the insertion of large reporter genes.

## Materials and methods

### Cloning of prime editor and pegRNA expression vectors

All protein expression vectors, including SpCas9(H840A)-RT, La-SpCas9(H840A)-RT, SpCas9(WT)-RT, and La-SpCas9(WT)-RT, were constructed under the control of the cytomegalovirus (CMV) promoter and subcloned using Gibson Assembly Master Mix (New England Biolabs, E2611L). Paired pegRNA expression plasmids encoding the target sequence, primer binding site (PBS), and reverse transcription template (RTT) regions were constructed under the control of a U6 promoter **(Supplementary Table S1)**. Target sequences were inserted into the pegRNA backbone via Golden Gate cloning using T4 ligase (NEB, M0202L) and dsDNA fragments with overhangs complementary to Bsa1-digested sites in the backbone vector. Digestion was performed using Bsa1, followed by ligation of a target-containing ssDNA fragment synthesized from Macrogen and annealed via gradual cooling. PBS and RTT sequences were inserted by digesting the previously modified pegRNA backbone vector with BsmB1 and ligating dsDNA fragments containing the PBS and RTT regions using the same method.

### Mammalian cell culture and transfection

HEK293FT cells were obtained from Invitrogen (R70007) and cultured in Dulbecco’s Modified Eagle’s Medium (Gibco, 11995065) supplemented with 10% fetal bovine serum (Gibco, 16000044) and 1× penicillin-streptomycin (Welgene, LS202-02) at 37 °C in a humidified 5% CO₂ incubator. Mutant HEK293FT-*ATXN3*-Q84 cells were maintained under identical conditions. Cells were seeded in 48-well plates (Corning, 3548) under the same conditions as previously described. Transfection was performed at ∼60% confluency after 16–20 h of incubation. For twin prime editing, 750 ng of SpCas9(H840A)-RT, La-SpCas9(H840A)-RT, SpCas9(WT)-RT, or La-SpCas9(WT)-RT and 125 ng of each pegRNA expression vector were used. For donor-mediated GFP insertion, 500 ng of SpCas9(H840A)-RT or La-SpCas9(H840A)-RT, 50 ng of each pegRNA vector, 200 ng of CMV-Bxb1 (Addgene #182142), and 200 ng of donor plasmid were co-transfected. All plasmids used in this study were purified using Nucleobond Xtra Midi Plus (MN740412.5) Transfections were performed using a mixture of 0.5 μL P3000 and 0.75 μL Lipofectamine 3000 (Thermo Fisher Scientific, L3000008) in 25 μL Opti-MEM (Gibco, 31985062) according to the manufacturer’s protocol.

### Genomic DNA extraction and amplicon preparation

Genomic DNA was extracted72 h post-plasmid transfection using the AccuPrep® Genomic DNA Extraction Kit (Bioneer, K-3032) following the manufacturer’s instructions. Target genomic loci were amplified via primary PCR using KOD One Master Mix (Toyobo, KMM-101) and primers listed in **Supplementary Table S2**. PCR was conducted under the following cycling conditions: 98 °C for 2 min; 30 cycles at 98 °C for 10 s, 60 °C for 10 s, 68 °C for 10 s; and a final extension at 68 °C for 5 min. Amplicons were further processed using nested PCR with Phusion High-Fidelity DNA (NEB, M0530L) to append Illumina sequencing adapters and barcodes **(Supplementary Table S2)** under the following conditions: 98 °C for 5 min; 30 cycles at 95°C for 2 min, 62 °C for 10 s, 72 °C for 10 s; and 72 °C for 5 min. PCR products were verified via 2% agarose gel electrophoresis and purified using a QIAquick PCR Purification Kit (Qiagen, 28106).

### Targeted amplicon sequencing and data analysis

Cas-OFFinder^26^ (http://www.rgenome.net/cas-offinder/) predicted genome-wide candidate off-target sites derived from the target sequence **(Supplementary Table S3)**. To evaluate potential off-target mutations via twin-prime editing, PCR was performed on the on-target site and the top 10 prioritized candidate (allowing up to three mismatches) off-target genome sites. Amplification products containing mutations (on-or off-target) were analyzed via next-generation sequencing (NGS) with barcoded nested PCR (denaturation at 98 °C for 30 s, primer annealing at 62 °C for 30 s, and elongation at 72 °C for 30 s, 35 cycles) using Phusion High-Fidelity DNA Polymerase (NEB, M0530L). Thereafter, the barcoded amplicon mixture was purified using the QIAquick PCR Purification Kit (Qiagen, 28106). Purified amplicons were loaded onto the mini-SEQ analyzer (Illumina MiniSeq system, SY-420-1001), and targeted deep sequencing was performed according to the manufacturer’s protocol. FastQ data were analyzed using Cas-analyzer (http://www.rgenome.net/cas-analyzer/). Frequencies of precise Twin-PE, only indels, and partial insertions (%) were calculated as a proportion of total allele frequency (frequency of Twin-PE, Only indel and Partial insertion (%)/total allele frequency (%)) and plotted using GraphPad Prism (10.4.1).

### Generation and characterization of the mutant HEK293FT-*ATXN3-*Q84 clonal cell line

HEK293FT cells (Invitrogen, R70007) were seeded in 24-well plates (Corning) and transfected at ∼60% confluency after 16–20 h. Transfection used 1 μg of a homemade Cas9-P2A-Blasticidin vector, 500 ng of each sgRNA1 (5′-TCCCAAAGTGCTGGGATTACAGG-3′) and sgRNA2 (GTATGTCAGATAAAGTGTGAAGG) vectors, and 1 μg donor DNA containing a Q84-expanded *ATXN3* exon 10 coding region flanked by 823 bp left and 807 bp right homology arms. Transfection was conducted via electroporation using an Amaxa electroporation kit (V4XC-2032; program: CM-130). After 3 d, the medium was replaced with fresh medium containing 10 μg/mL Blasticidin S, refreshed every 2 d. After 5 d of selection, surviving clones were limiting-diluted and plated in 96-well plates for single-cell culture. Genomic DNA was extracted using the AccuPrep® Genomic DNA Extraction Kit (Bioneer, K-3032), and successful insertion was confirmed via PCR and agarose gel electrophoresis.

### Triplet Repeat PCR (TP-PCR)

To detect CAG expansions in *ATXN3* exon 10, TP-PCR was performed following the method of Mulias et al. The 50 μL reaction included Phusion High-Fidelity DNA polymerase (NEB, M0530L), 10 ng genomic DNA or 1 pg donor plasmid, 1 μM forward and tailing primers, and 0.5 μM reverse primer **(Supplementary Table S2)**. Cycling conditions were as follows: 95 °C for 15 min; 40 cycles at 98 °C for 45 s, 60 °C for 1 min, and 72 °C for 2 min; followed by 72 °C for 5 min. PCR products were analyzed via fragment analysis at Macrogen and quantified using GeneMarker v2.6.3 (SoftGenetics).

### Western blot analysis

To assess polyQ repeat expression from *ATXN3* exon 10, HEK293FT and HEK293FT-*ATXN3*-Q84 cells were harvested via scrapping. Cell pellets were lysed in PRO-PREP buffer (Intron Biotechnology, 17081) following the manufacturer’s protocol. Equal amounts of total protein were resolved using 10% SDS-PAGE at 100 V, 2 h, transferred to nitrocellulose membranes (Bio-Rad, 1620115) at 0.3 A for 3 h, and blocked with 5% skim milk in blocking buffer (Biosesang, TR2005-100-74). Membranes were incubated overnight at 4 °C with anti-SCA3 (Sigma-Aldrich, MAB5360) and anti-β-actin (Santa Cruz, sc-47778) primary antibodies. After washing with TBST (0.1% Tween-20 in TBS), the HRP-conjugated goat anti-mouse secondary antibody was applied and incubated overnight at 4 °C. Detection was performed using enhanced chemiluminescence reagent (CYANGEN, Cod. XLS075,0100) and visualized with the Chemi DocXRS+ system (Bio-Rad, Model. No. Universal Hood II). Densitometry was analyzed using Image Lab software v3.0 (Bio-Rad, 12012931).

### RNA extraction and cDNA synthesis

Total RNA was extracted from HEK293FT-WT, HEK293FT-ATXN3-Q84, and HEK293FT-Q84 cells treated with La-SpCas9(H840A)-RT at 72 h post-transfection using the RNeasy Mini Kit (Qiagen, 74104) according to the manufacturer’s instructions. RNA quantity and purity were assessed using a NanoDrop One spectrophotometer (Thermo Scientific, Waltham, MA, USA, ND-ONE-W^4^). For cDNA synthesis, 1 µg RNA from each sample was used with the PrimeScript™ RT reagent Kit with gDNA Eraser (Takara, RR047A).

### Immunocytochemistry

To investigate the intracellular distribution of polyQ inclusions and ATXN3 protein in HEK293FT-WT, HEK293FT-ATXN3-Q84, and HEK293FT-Q84 cells treated with La-SpCas9(H840A)-RT, cells were seeded on poly-D-lysine-coated coverslips and incubated for 48 h. Thereafter, cells were washed with PBS and fixed with 4 % paraformaldehyde for 2 h at room temperature. Fixed cells were subsequently washed with PBS (Biosesang, PR2004-100-72) and permeabilized using 0.25 % PBST (PBS with 0.25 % Triton X-100). Subsequently, each cell sample was blocked with PBST containing 1% BSA (MP Biomedicals, 160069) for 30 min to reduce non-specific binding of primary antibodies. To detect polyQ inclusions and ATXN3 protein in each cell group, the samples were incubated overnight at 4 °C with either the anti-Polyglutamine-Expansion Disease Marker (Sigma-Aldrich, clone 5TF1-1C2, MAB1574) or the anti-Spinocerebellar Ataxia Type 3 antibodies (Sigma-Aldrich, clone 1H9, MAB5360), diluted in 1% BSA. After removing the primary antibody, coverslips were washed with PBS and incubated for 1 h at room temperature with the secondary antibody, Goat Anti-mouse lgG H&L (Abcam, Alexa Fluor 488, ab150113). Following secondary antibody incubation, cells were washed with PBS and mounted using antifade mounting medium containing DAPI (VECTASHIELD, H-1200-10) for nuclear counterstaining. Images were acquired using a confocal microscope (Carl Zeiss, Meditec, Dublin, CA, USA, LSM-700), and 3D surface plots of polyQ inclusions were generated using ImageJ software (NIH, Bethesda, MD, USA).

## Data availability

All relevant data that support the study findings are available from the corresponding author upon request. All targeted amplicon sequencing data were deposited in the NCBI Sequence Reads Archive database with accession numbers PRJNA1281332, SUB15409413.

## Author contributions

Conceptualization, L.W.G., J.B.S., Y.O. and S.H.L.; Methodology, L.W.G., J.B.S., Y.O., and S.H.L.; Software, L.W.G., Y.O., and S.H.L.; Validation, L.W.G., J.B.S., Y.O., and S.H.L.; Formal Analysis, L.W.G. and S.H.L.; Investigation, L.W.G. and S.H.L.; Resources, L.W.G., J.B.S., Y.O., Y.H.K., J.W.H., Y.L., and S.H.L.; Data Curation, L.W.G. and S.H.L.; Writing-Original Draft, L.W.G. and S.H.L.; Writing-Review & Editing, L.W.G., Y.L., and S.H.L.; Visualization, L.W.G. and S.H.L.; Supervision and Project Administration, Y.L. and S.H.L.; Funding Acquisition, Y.L. and S.H.L.

## Acknowledgements

This research was supported by grants from the National Research Foundation (NRF) funded by the Korean Ministry of Education, Science and Technology (RS-2025-00554011, RS-2025-02218918). This study was also supported by a grant of the Korea Health Technology R&D Project through the Korea Health Industry Development Institute (KHIDI), funded by the Ministry of Health & Welfare, Republic of Korea (grant number: RS-2024-00439579) and grants from the Korea Research Institute of Bioscience and Biotechnology (KRIBB; Research Initiative Program KGM4562532, KGM1072511).

## Life Sciences Reporting Summary

Further information on experimental design is available in the Life Sciences Reporting Summary.

