## supplementary information for "Optimizing twin prime editing components for scalable genome editing and therapy in spinocerebellar ataxia type 3"

#### Supplementary Tables

**Supplementary Table S1. Nucleotide sequences of pegRNAs used for twin prime editing at each DNA target in this study.** For each targeted gene, the sequences of the paired pegRNAs (sense and anti-sense) are presented in the 5′ to 3′ direction, along with the corresponding target DNA sequences. The PAM sequences (NGG) within the target DNA are highlighted in blue. The 3′ extension regions of the pegRNAs, comprising the PBS and RTT sequences, are also provided in the 5′ to 3′ orientation. The *attB* and *attP* sequences inserted into the target DNA are as follows:
*attB*: GGCTTGTCGACGACGGCGGTCTCCGTCGTCAGGATCAT,
*attP*: AGGTTTGTCTGGTCAACCACCGCGGTCTCAGTGGTGTACGGTACAAACCT
To enhance clarity, the scaffold regions of each pegRNA are omitted from the table and instead represented by a shared sequence as shown below:
Scaffold sequence in pegRNA for SpCas9:
GUUUUAGAGCUAGAAAUAGCAAGUUAAAAUAAGGCUAGUCCGUUAUCAACUU GAAAAAGUGGCACCGAGUCGGUGC

PAM: protospacer adjacent motif, PBS: primer binding site, RTT: reverse transcription template

| Gene | Target sequence (5’ to 3’) | 3’ extension (5’ to 3’) |
|---|---|---|
| *AAVS1*_Sense_***attB***_whole_overlap | GGGAAGGGGCAGGAGAGCCAGG | AGGGGCAGGAGAGGGCTTGTCGACGACGGCGGTCTCCGTCGTCAGGATCAT |
| *AAVS1*_Anti-Sense_***attB***_whole_overlap | GTCCTTGGCAAGCCCAGGAGAGG | TTGGCAAGCCCAGATGATCCTGACGACGGAGACCGCCGTCGTCGACAAGCC |
| *AAVS1*_Sense_***attB***_Homology arm | GGGAAGGGGCAGGAGAGCCAGG | AGGGGCAGGAGAGGGCTTGTCGACGACGGCGGTCTCCGTCGTCAGGATCATCTGGGCTTGCCAAGG |
| *AAVS1*_Anti-Sense_***attB***_Homology arm | GTCCTTGGCAAGCCCAGGAGAGG | TTGGCAAGCCCAGATGATCCTGACGACGGAGACCGCCGTCGTCGACAAGCCCTCTCCTGCCCCTTC |
| *AAVS1*_Sense_***attB***_20bp_overlap | GGGAAGGGGCAGGAGAGCCAGG | AGGGGCAGGAGAGGGCTTGTCGACGACGGCGGTCTCCGTCGT |

| | | |
|---|---|---|
| *AAVS1*_Anti-Sense_**attB**_20bp_overlap | GTCCTTGGCAAGCCCAGGAGAGG | TTGGCAAGCCCAGATGATCCTGACGACGGAGACCGCCGTCGT |
| *AAVS1*_Sense_**attP**_whole_overlap | GGGAAGGGGCAGGAGAGCCAGG | AGGGGCAGGAGAGAGGTTTGTCTGGTCAACCACCGCGGTCTCAGTGGTGTACGGTACAAACCT |
| *AAVS1*_Anti-Sense_**attP**_whole_overlap | GTCCTTGGCAAGCCCAGGAGAGG | TTGGCAAGCCCAGAGGTTTGTACCGTACACCACTGAGACCGCGGTGGTTGACCAGACAAACCT |
| *AAVS1*_Sense_**attP**_Homology arm | GGGAAGGGGCAGGAGAGCCAGG | AGGGGCAGGAGAGAGGTTTGTCTGGTCAACCACCGCGGTCTCAGTGGTGTACGGTACAAACCTCTGGGCTTGCCAAGG |
| *AAVS1*_Anti-Sense_**attP**_Homology arm | GTCCTTGGCAAGCCCAGGAGAGG | TTGGCAAGCCCAGAGGTTTGTACCGTACACCACTGAGACCGCGGTGGTTGACCAGACAAACCTCTCTCCTGCCCCTTC |
| *AAVS1*_Sense_**attP**_28bp_overlap | GGGAAGGGGCAGGAGAGCCAGG | AGGGGCAGGAGAGAGGTTTGTCTGGTCAACCACCGCGGTCTCAGTGGTGTAC |
| *AAVS1*_Anti-Sense_**attP**_28bp_overlap | GTCCTTGGCAAGCCCAGGAGAGG | TTGGCAAGCCCAGAGGTTTGTACCGTACACCACTGAGACCGCGGTGGTTGAC |
| *HEK3*_Sense_**attB**_whole_overlap | GGCCCAGACTGAGCACGTGATGG | CAGACTGAGCACGGGCTTGTCGACGACGGCGGTCTCCGTCGTCAGGATCAT |
| *HEK3*_Anti-Sense_**attB**_whole_overlap | TGTCCTGCGACGCCCTCTGGAGG | ACCAGTATCCCGGATGATCCTGACGACGGAGACCGCCGTCGTCGACAAGCC |
| *HEK3*_Sense_**attB**_Homology arm | GGCCCAGACTGAGCACGTGATGG | CAGACTGAGCACGGGCTTGTCGACGACGGCGGTCTCCGTCGTCAGGATCATCCGGGATACTGGTTG |
| *HEK3*_Anti-Sense_**attB**_Homology arm | TGTCCTGCGACGCCCTCTGGAGG | ACCAGTATCCCGGATGATCCTGACGACGGAGACCGCCGTCGTCGACAAGCCCGTGCTCAGTCTGGG |
| *HEK3*_Sense_**attB**_20bp_overlap | GGCCCAGACTGAGCACGTGATGG | CAGACTGAGCACGGGCTTGTCGACGACGGCGGTCTCCGTCGT |
| *HEK3*_Anti-Sense_**attB**_20bp_overlap | TGTCCTGCGACGCCCTCTGGAGG | ACCAGTATCCCGGATGATCCTGACGACGGAGACCGCCGTCGT |
| *FANCF*_Sense_**attB**_whole_overlap | GGAATCCCTTCTGCAGCACCTGG | TCCCAGGTGCTGAATGATCCTGACGACGGAGACCGCCGTCGTCGACAAGCC |
| *FANCF*_Anti-Sense_**attB**_whole_overlap | GGGGTCCCAGGTGCTGACGTAGG | TCCCTTCTGCAGCGGCTTGTCGACGACGGCGGTCTCCGTCGTCAGGATCAT |
| *FANCF*_Sense_**attB**_Homology arm | GGAATCCCTTCTGCAGCACCTGG | TCCCAGGTGCTGAATGATCCTGACGACGGAGACCGCCGTCGTCGACAAGCCGCTGCAGAAGGGATT |
| *FANCF*_Anti-Sense_**attB**_Homology arm | GGGGTCCCAGGTGCTGACGTAGG | TCCCTTCTGCAGCGGCTTGTCGACGACGGCGGTCTCCGTCGTCAGGATCATTCAGCACCTGGGACC |
| *FANCF*_Sense_**attB**_20bp_overlap | GGAATCCCTTCTGCAGCACCTGG | TCCCAGGTGCTGAATGATCCTGACGACGGAGACCGCCGTCGT |
| *FANCF*_Anti-Sense_**attB**_20bp_overlap | GGGGTCCCAGGTGCTGACGTAGG | TCCCTTCTGCAGCGGCTTGTCGACGACGGCGGTCTCCGTCGT |
| *CCR5*_Sense_**attB**_whole_overlap | GCTGTGTTTGCGTCTCTCCCAGG | TGTTTGCGTCTCTGGCTTGTCGACGACGGCGGTCTCCGTCGTCAGGATCAT |
| *CCR5*_Anti-Sense_**attB**_whole_overlap | GTATGGAAAATGAGAGCTGCAGG | GGAAAATGAGAGCATGATCCTGACGACGGAGACCGCCGTCGTCGACAAGCC |
| *Emx1*_Sense_**attB**_whole_overlap_1 | TGGTTGCCCACCCTAGTCATTGG | TGCCCACCCTAGTGGCTTGTCGACGACGGCGGTCTCCGTCGTCAGGATCAT |
| *Emx1*_Sense_**attB**_whole_overlap_2 | ACTCTGCCCTCGTGGGTTTGTGG | TGCCCTCGTGGGTGGCTTGTCGACGACGGCGGTCTCCGTCGTCAGGATCAT |
| *Emx1*_Anti_Sense_**attB**_whole_overlap | ATTGCCACGAAGCAGGCCAATGG | CCACGAAGCAGGCATGATCCTGACGACGGAGACCGCCGTCGTCGACAAGCC |

| | | |
|---|---|---|
| *VEGFA*_Sense_***attB***_whole_overlap | CAGGCTGCACCCATGGCAGAAGG | CTGCACCCATGGCGGCTTGTCGACGACGGCGGTCTCCGTCGTCAGGATCAT |
| *VEGFA*_Anti-Sense_***attB***_whole_overlap | GGGAACCCCATCCAACAGCCAGG | ACCCCATCCAACAATGATCCTGACGACGGAGACCGCCGTCGTCGACAAGCC |
| *DAPK1*_Sense_***attB***_whole_overlap | CGTGTTCAGGCAGGAAAACGTGG | TTCAGGCAGGAAAGGCTTGTCGACGACGGCGGTCTCCGTCGTCAGGATCAT |
| *DAPK1*_Anti-Sense_***attB***_whole_overlap | TTACCTGCCAAGTTCCTCGCCGG | CTGCCAAGTTCCTATGATCCTGACGACGGAGACCGCCGTCGTCGACAAGCC |
| *PIN1*_Sense_***attB***_whole_overlap_1 | AGCCAGTGGGAGCGGCCCAGCGG | AGTGGGAGCGGCCGGCTTGTCGACGACGGCGGTCTCCGTCGTCAGGATCAT |
| *PIN1*_Sense_***attB***_whole_overlap_2 | AGCGGCAACAGCAGCAGTGGTGG | GCAACAGCAGCAGGGCTTGTCGACGACGGCGGTCTCCGTCGTCAGGATCAT |
| *PIN1*_Anti-Sense_***attB***_whole_overlap_1 | TGCGAGCAGCGGACCCTGGCAGG | AGCAGCGGACCCTATGATCCTGACGACGGAGACCGCCGTCGTCGACAAGCC |
| *PIN1*_Anti-Sense_***attB***_whole_overlap_2 | ACTGGCTGTGCTTCACCAGCAGG | GCTGTGCTTCACCATGATCCTGACGACGGAGACCGCCGTCGTCGACAAGCC |
| *GAPDH*_Sense_***attB***_whole_overlap_1 | TCTAGGTATGACAACGAATTTGG | GGTATGACAACGAGGCTTGTCGACGACGGCGGTCTCCGTCGTCAGGATCAT |
| *GAPDH*_Sense_***attB***_whole_overlap_2 | ATTTGGCTACAGCAACAGGGTGG | GGCTACAGCAACAGGCTTGTCGACGACGGCGGTCTCCGTCGTCAGGATCAT |
| *GAPDH*_Anti_Sense_***attB***_whole_overlap_1 | TGGAGGCCATGTGGGCCATGAGG | GGCCATGTGGGCCATGATCCTGACGACGGAGACCGCCGTCGTCGACAAGCC |
| *ATXN3*_Sense_1_***attB***_insertion | CTCAAGCCATCCACCCGCCTCGG | AGCCATCCACCCGggcttgtcgacgacggcggtctccgtcgtcaggatcat |
| *ATXN3*_Sense_2_***attB***_insertion | CCACCACTCCTGGCCATGATAGG | CACTCCTGGCCATggcttgtcgacgacggcggtctccgtcgtcaggatcat |
| *ATXN3*_Anti-Sense_1_***attB***_insertion | GTATGTCAGATAAAGTGTGAAGG | GTCAGATAAAGTGatgatcctgacgacggagaccgccgtcgtcgacaagcc |
| *ATXN3*_Anti-Sense_2_***attB***_insertion | GATTCTTAATATGATTAAAGAGG | CTTAATATGATTAatgatcctgacgacggagaccgccgtcgtcgacaagcc |
| *ATXN3*_Anti-Sense_3_***attB***_insertion | TGGTAACTGCTCCTTAATCCAGG | AACTGCTCCTTAAatgatcctgacgacggagaccgccgtcgtcgacaagcc |
| *ATXN3*_Anti-Sense_4_***attB***_insertion | CTAGCAATCCTCTCCTGCCTTGG | CAATCCTCTCCTGatgatcctgacgacggagaccgccgtcgtcgacaagcc |
| *ATXN3*_AntiSense_5_***attB***_insertion | AAAAATATCCATGATGTCTAAGG | ATATCCATGATGTatgatcctgacgacggagaccgccgtcgtcgacaagcc |
| *ATXN3*_Sense_1_TAA_***attB***_TAA_insertion | CTCAAGCCATCCACCCGCCTCGG | AGCCATCCACCCGtttcagacagtaaggcttgtcgacgacggcggtctccgtcgtcaggatcatctaa |
| *ATXN3*_Anti-Sense_1_TAA_***attB***_TAA _insertion | GTATGTCAGATAAAGTGTGAAGG | GTCAGATAAAGTGttagatgatcctgacgacggagaccgccgtcgtcgacaagccttactgtctgaaa |
| *ATXN3*_Anti-Sense_3_TAA_***attB***_TAA _insertion | TGGTAACTGCTCCTTAATCCAGG | AACTGCTCCTTAAttagatgatcctgacgacggagaccgccgtcgtcgacaagccttactgtctgaaa |
| *ATXN3*_Sense_1_TAG_***attB***_TAG_insertion | CTCAAGCCATCCACCCGCCTCGG | AGCCATCCACCCGtttcagacagtagggcttgtcgacgacggcggtctccgtcgtcaggatcatctag |
| *ATXN3*_Anti-Sense_1_TAG_***attB***_TAG _insertion | GTATGTCAGATAAAGTGTGAAGG | GTCAGATAAAGTGctagatgatcctgacgacggagaccgccgtcgtcgacaagccctactgtctgaaa |

| | | |
|---|---|---|
| *ATXN3*_Anti-Sense_3_ TAG_***attB***_TAG _insertion | TGGTAACTGCTCCTT AATCCAGG | AACTGCTCCTTAActagatgatcctgacgacggagaccgccgtcgtcgacaa gccctactgtctgaaa |

**Supplementary Table S2. Nucleotide sequences of DNA primers used in this study.**

The sequences of DNA primers used to amplify genomic regions encompassing the target editing sites within each gene are listed. For next-generation sequencing (NGS), the forward and reverse adaptor primers used for the production of target-specific amplicons are highlighted in blue.

| Target gene (primer direction) | DNA sequence (5' to 3') |
|---|---|
| *AAVS1*_F1 | TCACCCAGAGACAGTGACCA |
| *AAVS1*_R1 | CTCCGTGCGTCAGTTTTACC |
| *AAVS1*_DF | ACACTCTTTCCCTACACGACGCTCTTCCGATCTGGGCTCAGTCTGAAGAGCAG |
| *AAVS1*_DR | GTGACTGGAGTTCAGACGTGTGCTCTTCCGATCTCCCAGTGCTATCTGGGACAT |
| *HEK3*_F1 | TATATGGCTACCTAAGGTTGA |
| *HEK3*_R1 | GTTTTCTCTTGGGCAATATGG |
| *HEK3*_DF | ACACTCTTTCCCTACACGACGCTCTTCCGATCTTTTCTGCTGCAAGTAAGCAT |
| *HEK3*_DR | GTGACTGGAGTTCAGACGTGTGCTCTTCCGATCTCTCCCTAGGTGCTGGCTTC |
| *FANCF*_F1 | CTTCATCAGAGAGTCCTCCTGG |
| *FANCF*_R1 | GGATATTTCCAAAGCGAAAGGAAGC |
| *FANCF*_DF | ACACTCTTTCCCTACACGACGCTCTTCCGATCTGATGGATGTGGCGCAGGTAG |
| *FANCF*_DR | GTGACTGGAGTTCAGACGTGTGCTCTTCCGATCTCAATCAGTACGCAGAGAGTCG |
| *CCR5*_F1 | CAAAAGGCTGAAGAGCATGA |
| *CCR5*_R1 | CATTTGCAGAAGCGTTTGG |
| *CCR5*_DF | ACACTCTTTCCCTACACGACGCTCTTCCGATCTTTAAAAGCCAGGACGGTCAC |
| *CCR5*_DR | GTGACTGGAGTTCAGACGTGTGCTCTTCCGATCTGACCAGCCCCAAGATGACTA |
| *Emx1*_F1 | TGCCCGTGTCATTAAGAGAGA |
| *Emx1*_R1 | TTCTCTCTGGCCCACTGTGT |
| *Emx1*_DF | ACACTCTTTCCCTACACGACGCTCTTCCGATCTAGTGGCCAGAGTCCAGCTT |
| *Emx1*_DR | GTGACTGGAGTTCAGACGTGTGCTCTTCCGATCTCAGAAGCTGGAGGAGGAAGG |
| *VEGFA*_F1 | GTTTTTCTCGCCCCTAGTCC |
| *VEGFA*_R1 | GCACGACTCCTTCTCCAAAT |
| *VEGFA*_DF | ACACTCTTTCCCTACACGACGCTCTTCCGATCTATGCCCATGCCTTGCTCT |
| *VEGFA*_DR | GTGACTGGAGTTCAGACGTGTGCTCTTCCGATCTTTATATTCCTGTGCCCCTTCC |
| *DAPK1*_F1 | GGCATGTGTGCAGAGAAAGG |
| *DAPK1*_R1 | TGTTGTCACAGAAGGGCAAG |
| *DAPK1*_DF | ACACTCTTTCCCTACACGACGCTCTTCCGATCTGAAGCGGAGCTGAAGTGC |
| *DAPK1*_DR | GTGACTGGAGTTCAGACGTGTGCTCTTCCGATCTACGCCTGATTCAAACTGCAT |
| *PIN1*_F1 | CCAGCTTCCTCTGTTCCATC |
| *PIN1*_R1 | TCACAGAGGCACAGCTACTACC |
| *PIN1*_DF | ACACTCTTTCCCTACACGACGCTCTTCCGATCTGCCGAGTGTACTACTTCAACCA |
| *PIN1*_DR | GTGACTGGAGTTCAGACGTGTGCTCTTCCGATCTCTCCAGGGCCTCCTCCTT |
| *GAPDH*_F1 | ATCCTGGGCTACACTGAGCA |
| *GAPDH*_R1 | GCTGCCCACAGAATAGCTTC |

| | |
|---|---|
| *GAPDH*_DF | ACACTCTTTCCCTACACGACGCTCTTCCGATCTGGTGGCTGGCTCAGAAAA |
| *GAPDH*_DR | GTGACTGGAGTTCAGACGTGTGCTCTTCCGATCTGACTGAGTGTGGCAGGGACT |
| *HEK3*_S_OT_1_F | AGTAGCCAGACCCCCAATTT |
| *HEK3*_S_OT_1_R | GGGAACTGTCGCAGTCTGA |
| *HEK3*_S_OT_1_DF | ACACTCTTTCCCTACACGACGCTCTTCCGATCTAGCATGAACCAGTCAAAAAGTT |
| *HEK3*_S_OT_1_DR | GTGACTGGAGTTCAGACGTGTGCTCTTCCGATCTTGTCGTCGCTGACAATTTCT |
| *HEK3*_S_OT_2_F | AGAGCTCCTGGAGAGGGAGT |
| *HEK3*_S_OT_2_R | CTCTTTGGGCTCTGGGTATG |
| *HEK3*_S_OT_2_DF | ACACTCTTTCCCTACACGACGCTCTTCCGATCTCCAGATTCCTGGTCCAAAGG |
| *HEK3*_S_OT_2_DR | GTGACTGGAGTTCAGACGTGTGCTCTTCCGATCTCAGGGACTGAGAGGGAACAG |
| *HEK3*_S_OT_3_F | TGGCCTTGATTTTAAGCATGA |
| *HEK3*_S_OT_3_R | TCTCCTTGCTTCTGCAGTCA |
| *HEK3*_S_OT_3_DF | ACACTCTTTCCCTACACGACGCTCTTCCGATCTAAGGAGCAGCTCTTCCTGGT |
| *HEK3*_S_OT_3_DR | GTGACTGGAGTTCAGACGTGTGCTCTTCCGATCTACTGACTCCCTCCCCTGGT |
| *HEK3*_S_OT_4_F | GGGCAGCTCAGAGTGGATAG |
| *HEK3*_S_OT_4_R | CCTGTTGGGAAGTGCCTCTA |
| *HEK3*_S_OT_4_DF | ACACTCTTTCCCTACACGACGCTCTTCCGATCTCACTCCCGAGGGAGGTCT |
| *HEK3*_S_OT_4_DR | GTGACTGGAGTTCAGACGTGTGCTCTTCCGATCTCCATGTGTGTTGGAGACACC |
| *HEK3*_S_OT_5_F | TCTGCTTTTGAGGGTGTGAA |
| *HEK3*_S_OT_5_R | AGGACATGGAGCACCCAAAG |
| *HEK3*_S_OT_5_DF | ACACTCTTTCCCTACACGACGCTCTTCCGATCTCAAACCTACCCTCTCCCTCA |
| *HEK3*_S_DT_5_DR | GTGACTGGAGTTCAGACGTGTGCTCTTCCGATCTCCACAAACTTTCGGAACCAG |
| *HEK3*_S_OT_6_F | ATGTGCCACTGACAGTCACC |
| *HEK3*_S_OT_6_R | CACACCTCGCTCCATCTACC |
| *HEK3*_S_OT_6_DF | ACACTCTTTCCCTACACGACGCTCTTCCGATCTGCCAAGGGAGTCACTGTCAT |
| *HEK3*_S_OT_6_DR | GTGACTGGAGTTCAGACGTGTGCTCTTCCGATCTACATTCTGGGTCCTGAGGTG |
| *HEK3*_S_OT_7_F | ACGGGGAGCCTGGATTTA |
| *HEK3*_S_OT_7_R | TCCACTGCCCCTGAGTCC |
| *HEK3*_S_OT_7_DF | ACACTCTTTCCCTACACGACGCTCTTCCGATCTCACTGGTTTCGTGGAAGGTAA |
| *HEK3*_S_OT_7_DR | GTGACTGGAGTTCAGACGTGTGCTCTTCCGATCTCCAGCACTCCAGGACTCTAAA |
| *HEK3*_S_OT_8_F | GGGCAGAAGACGGTGATAAG |
| *HEK3*_S_OT_8_R | TCCTCTGCCCAGTAGCTGTT |
| *HEK3*_S_OT_8_DF | ACACTCTTTCCCTACACGACGCTCTTCCGATCTGCTTTTCAGCTGGGAAATTG |
| *HEK3*_S_OT_8_DR | GTGACTGGAGTTCAGACGTGTGCTCTTCCGATCTCTTACAGGCTGGATGTGGTG |
| *HEK3*_S_OT_9_F | CAGGAGGCCCAGATATACGA |
| *HEK3*_S_OT_9_R | ACACCATACACGCACACACA |
| *HEK3*_S_OT_9_DF | ACACTCTTTCCCTACACGACGCTCTTCCGATCTGCTGGTGTGCTCTCATCACT |
| *HEK3*_S_OT_9_DR | GTGACTGGAGTTCAGACGTGTGCTCTTCCGATCTTGTGCATGCTCACACAGAGA |
| *HEK3*_AS_OT_1_F | GAGAAAGAAAACTGGGTGGCTA |
| *HEK3*_AS_OT_1_R | GGGTGTGAGTAAAGTCAAACTCTG |
| *HEK3*_AS_OT_1_DF | ACACTCTTTCCCTACACGACGCTCTTCCGATCTTTACAGGTGTGAGCCACTGC |
| *HEK3*_AS_OT_1_DR | GTGACTGGAGTTCAGACGTGTGCTCTTCCGATCTGCTTGCCCTTTTTCCTTACC |
| *HEK3*_AS_OT_2_F | GCCAAATCAGGTGTTCTGCT |
| *HEK3*_AS_OT_2_R | ACATTGTGACCCAATCCACA |
| *HEK3*_AS_OT_2_DF | ACACTCTTTCCCTACACGACGCTCTTCCGATCTGGACTTCACCAAGGCTGCTA |
| *HEK3*_AS_OT_2_DR | GTGACTGGAGTTCAGACGTGTGCTCTTCCGATCTGAAAACAGAGGGGCTGTGTC |

| | |
|---|---|
| *HEK3*_AS_OT_3_F | GCCCCATCCTCTAAACCTTG |
| *HEK3*_AS_OT_3_R | TGTTGAATGGTCTCAGCAAA |
| *HEK3*_AS_OT_3_DF | ACACTCTTTCCCTACACGACGCTCTTCCGATCTCTGATGCATGAAGGATGAGTG |
| *HEK3*_AS_OT_3_DR | GTGACTGGAGTTCAGACGTGTGCTCTTCCGATCTCAGGTTCAGCCTCCAAGAAA |
| *HEK3*_AS_OT_4_F | GGACCACCTGTAGGGACAGA |
| *HEK3*_AS_OT_4_R | GTGACAGGCATCACCTCCTT |
| *HEK3*_AS_OT_4_DF | ACACTCTTTCCCTACACGACGCTCTTCCGATCTGTTGGATGGAGGCCGTTAG |
| *HEK3*_AS_OT_4_DR | GTGACTGGAGTTCAGACGTGTGCTCTTCCGATCTGCTTGGCTCCCACAAGTCTA |
| *HEK3*_AS_OT_5_F | CCAGTGCTTAGAACCGAGCTT |
| *HEK3*_AS_OT_5_R | ATTCCACGGAAATTCCTCCT |
| *HEK3*_AS_OT_5_DF | ACACTCTTTCCCTACACGACGCTCTTCCGATCTCCTCTTTGGGGAGGGATTTA |
| *HEK3*_AS_DT_5_DR | GTGACTGGAGTTCAGACGTGTGCTCTTCCGATCTCAACCAGGGCAACAGAAACT |
| *HEK3*_AS_OT_6_F | GGAAAAATAAATGCAAGAGCTGA |
| *HEK3*_AS_OT_6_R | GCACCTAGATCCACACTCTGC |
| *HEK3*_AS_OT_6_DF | ACACTCTTTCCCTACACGACGCTCTTCCGATCTGGCCTCCAATTTTCAGGAGT |
| *HEK3*_AS_OT_6_DR | GTGACTGGAGTTCAGACGTGTGCTCTTCCGATCTGTAGGGTCCAGCCACTTCAG |
| *HEK3*_AS_OT_7_F | TTCCTCCTCTGCAATAAAGCTC |
| *HEK3*_AS_OT_7_R | AGTTCAGTTGGCGGTGTCTC |
| *HEK3*_AS_OT_7_DF | ACACTCTTTCCCTACACGACGCTCTTCCGATCTAATACGCAGCCCAGTGAAAG |
| *HEK3*_AS_OT_7_DR | GTGACTGGAGTTCAGACGTGTGCTCTTCCGATCTGCCCATTGGCGATTAGTAAC |
| *VEGFA*_S_OT_1_F | TGTCTCAATCTTCTTTAGCATCTGA |
| *VEGFA*_S_OT_1_R | CATAACTGTGATGCCCTAAGCA |
| *VEGFA*_S_OT_1_DF | ACACTCTTTCCCTACACGACGCTCTTCCGATCTGCAAATATTCTTTCCTTTGTGC |
| *VEGFA*_S_OT_1_DR | GTGACTGGAGTTCAGACGTGTGCTCTTCCGATCTGTCTTCCCTAGGGCCCTTTC |
| *VEGFA*_S_OT_2_F | CATGCGCCTAACCTCCTTAT |
| *VEGFA*_S_OT_2_R | AGGGCCAGAGCTCAGACAG |
| *VEGFA*_S_OT_2_DF | ACACTCTTTCCCTACACGACGCTCTTCCGATCTACAGCTTTGGCAGAGTGGAC |
| *VEGFA*_S_OT_2_DR | GTGACTGGAGTTCAGACGTGTGCTCTTCCGATCTGACCGTCAAGGCCCTTTT |
| *VEGFA*_S_OT_3_F | CTCCTGGGCAGACCATAAAA |
| *VEGFA*_S_OT_3_R | CATCCCCTCTTCTGCCTTTC |
| *VEGFA*_S_OT_3_DF | ACACTCTTTCCCTACACGACGCTCTTCCGATCTGTGAGCAGGCTGGTCTTAGG |
| *VEGFA*_S_OT_3_DR | GTGACTGGAGTTCAGACGTGTGCTCTTCCGATCTTCTCTCTTCCACACCGTTCA |
| *VEGFA*_S_OT_4_F | CAGATCTGCCGTCCTTGG |
| *VEGFA*_S_OT_4_R | GTTCTGGGCTGTTGATGGAT |
| *VEGFA*_S_OT_4_DF | ACACTCTTTCCCTACACGACGCTCTTCCGATCTCCTGCAGAGGGAGAGATGAG |
| *VEGFA*_S_OT_4_DR | GTGACTGGAGTTCAGACGTGTGCTCTTCCGATCTGATGTGCTCTGTTTCCCTAAAAA |
| *VEGFA*_S_OT_5_F | AATTCCCAGTGTGGTGGTGT |
| *VEGFA*_S_OT_5_R | CAAACAGACTAGCTCAATTACAGACC |
| *VEGFA*_S_OT_5_DF | ACACTCTTTCCCTACACGACGCTCTTCCGATCTAAAAGAGCTGGGTGTTAAAAAGC |
| *VEGFA*_S_DT_5_DR | GTGACTGGAGTTCAGACGTGTGCTCTTCCGATCTAAAGATTTGGCTCATGATTGTG |
| *VEGFA*_S_OT_6_F | CCAGGGCCTTGAAAATACAT |
| *VEGFA*_S_OT_6_R | TTTCTGAAAAGAAATACTTTCAAACTT |
| *VEGFA*_S_OT_6_DF | ACACTCTTTCCCTACACGACGCTCTTCCGATCTTAAATGGCATCGGCATTCTG |
| *VEGFA*_S_OT_6_DR | GTGACTGGAGTTCAGACGTGTGCTCTTCCGATCTAAAACTAGCATGTGGCTGCTC |
| *VEGFA*_S_OT_7_F | TGTCTGACAGAATTCAGCTGTG |

| | |
|---|---|
| *VEGFA*_S_OT_7_R | GGGTCCTGAGGCAACATACA |
| *VEGFA*_S_OT_7_DF | ACACTCTTTCCCTACACGACGCTCTTCCGATCTTCCATCTCCTCTAGGTTTTCTAGTTT |
| *VEGFA*_S_OT_7_DR | GTGACTGGAGTTCAGACGTGTGCTCTTCCGATCTTTGCAGTAAAGACAGGCATAAGA |
| *VEGFA*_S_OT_8_F | CCTCCCTGCGAAGAGTTTAC |
| *VEGFA*_S_OT_8_R | ACATGACGGCAGGAAGAAAC |
| *VEGFA*_S_OT_8_DF | ACACTCTTTCCCTACACGACGCTCTTCCGATCTAGGAGGGACACAGATGCAAG |
| *VEGFA*_S_OT_8_DR | GTGACTGGAGTTCAGACGTGTGCTCTTCCGATCTTCGGTGCTGTGTTTGATGTT |
| *VEGFA*_S_OT_9_F | GTCTGCCTCACCCCAGACTA |
| *VEGFA*_S_OT_9_R | TCTGAGTCTGTTTCCTCCTTTG |
| *VEGFA*_S_OT_9_DF | ACACTCTTTCCCTACACGACGCTCTTCCGATCTGCCAGCTGCTTTACAAAGGT |
| *VEGFA*_S_OT_9_DR | GTGACTGGAGTTCAGACGTGTGCTCTTCCGATCTAGGCAATAATGATAGAGGCACA |
| *VEGFA*_S_OT_10_F | GCAACAGCCTCAGCAAATAA |
| *VEGFA*_S_OT_10_R | CTTGGTCATGGAGAATGCAG |
| *VEGFA*_S_OT_10_DF | ACACTCTTTCCCTACACGACGCTCTTCCGATCTTTGAAGCCCTCTCACTAGCC |
| *VEGFA*_S_OT_10_DR | GTGACTGGAGTTCAGACGTGTGCTCTTCCGATCTAGGGGTTTAAGGCATTCTGG |
| *VEGFA*_AS_OT_1_F | GCCAAAACAAAGGGGTTACA |
| *VEGFA*_AS_OT_1_R | GGTGTGTGTGTGCCTGTGA |
| *VEGFA*_AS_OT_1_DF | ACACTCTTTCCCTACACGACGCTCTTCCGATCTCATGGTCTTGGGCATCTCTC |
| *VEGFA*_AS_OT_1_DR | GTGACTGGAGTTCAGACGTGTGCTCTTCCGATCTCAAGCACTGAGGTGCTCACA |
| *VEGFA*_AS_OT_2_F | GCTCACAGCCACTCTATGAGG |
| *VEGFA*_AS_OT_2_R | TATGAAGCCAAGCACGACTC |
| *VEGFA*_AS_OT_2_DF | ACACTCTTTCCCTACACGACGCTCTTCCGATCTGGCTTCTGTCTTTGGTGCTC |
| *VEGFA*_AS_OT_2_DR | GTGACTGGAGTTCAGACGTGTGCTCTTCCGATCTGACTGGGGCAGGATCTTCTT |
| *VEGFA*_AS_OT_3_F | ATTTGCTGCTGACTCCCAAA |
| *VEGFA*_AS_OT_3_R | CCAGTCCAGATGTTGGATGA |
| *VEGFA*_AS_OT_3_DF | ACACTCTTTCCCTACACGACGCTCTTCCGATCTCACAGCTGCTATTCCTTTGC |
| *VEGFA*_AS_OT_3_DR | GTGACTGGAGTTCAGACGTGTGCTCTTCCGATCTTGGCTTTACCAGTGGACTCA |
| *VEGFA*_AS_OT_4_F | ATCTCCAGTGGGAAGTCAGG |
| *VEGFA*_AS_OT_4_R | TGAGAGAAGGCAGACATGGA |
| *VEGFA*_AS_OT_4_DF | ACACTCTTTCCCTACACGACGCTCTTCCGATCTCTCTAGGGAGGAATGACATGG |
| *VEGFA*_AS_OT_4_DR | GTGACTGGAGTTCAGACGTGTGCTCTTCCGATCTTTGCTAGTTGATTTCCTGATGA |
| *VEGFA*_AS_OT_5_F | CTCACATGCGTTCAGTTGGT |
| *VEGFA*_AS_OT_5_R | GTGAAAGAACTGCCCCTTTG |
| *VEGFA*_AS_OT_5_DF | ACACTCTTTCCCTACACGACGCTCTTCCGATCTTCTGACGTCACGATGTTCTTG |
| *VEGFA*_AS_DT_5_DR | GTGACTGGAGTTCAGACGTGTGCTCTTCCGATCTTGGCCATCATGAGTAGTGGA |
| *VEGFA*_AS_OT_6_F | CTGCAAACGGATGCTGCT |
| *VEGFA*_AS_OT_6_R | CCAAGACGGCCACATAGAAA |
| *VEGFA*_AS_OT_6_DF | ACACTCTTTCCCTACACGACGCTCTTCCGATCTTGATCTGAGGGTTGCAAAGA |
| *VEGFA*_AS_OT_6_DR | GTGACTGGAGTTCAGACGTGTGCTCTTCCGATCTCAGGTACTCACAGTGGGACTGA |
| *VEGFA*_AS_OT_7_F | TCCCTAGTAAAATCCCAGTTCTTTT |
| *VEGFA*_AS_OT_7_R | GGGCTACTTCCTGCTCCATA |
| *VEGFA*_AS_OT_7_DF | ACACTCTTTCCCTACACGACGCTCTTCCGATCTCCATTGGTCTTGGCTCACTT |
| *VEGFA*_AS_OT_7_DR | GTGACTGGAGTTCAGACGTGTGCTCTTCCGATCTGCTCCTTGTTCCTTCCCACT |
| *VEGFA*_AS_OT_8_F | CCCACAGCTTCTCTCCTCAG |
| *VEGFA*_AS_OT_8_R | CGAGCATTTTTATGACCATTATTTT |

| | |
|---|---|
| *VEGFA*_AS_OT_8_DF | ACACTCTTTCCCTACACGACGCTCTTCCGATCTCAAGTAAGATTGCAGATGGTGGT |
| *VEGFA*_AS_OT_8_DR | GTGACTGGAGTTCAGACGTGTGCTCTTCCGATCTTTCATATGTTTTCTTCACTTTTCTTGA |
| *VEGFA*_AS_OT_9_F | GGCTCAGAGACTATCCCCATT |
| *VEGFA*_AS_OT_9_R | CCACCACTCCTGGCTAATTT |
| *VEGFA*_AS_OT_9_DF | ACACTCTTTCCCTACACGACGCTCTTCCGATCTAGCCTCACTGATGCACTGGT |
| *VEGFA*_AS_OT_9_DR | GTGACTGGAGTTCAGACGTGTGCTCTTCCGATCTCTCCCAAAGTGCTGGGATTA |
| *VEGFA*_AS_OT_10_F | TTTTTCCTCACAGACGGAGT |
| *VEGFA*_AS_OT_10_R | CAGCCGGATGTTTAAAAGTTCT |
| *VEGFA*_AS_OT_10_DF | ACACTCTTTCCCTACACGACGCTCTTCCGATCTTCTTTCCTCTGTGGGCAGAT |
| *VEGFA*_AS_OT_10_DR | GTGACTGGAGTTCAGACGTGTGCTCTTCCGATCTCAGCCAGATAAAACCCCTTC |
| *PIN1*_S_2_OT_1_F | CACCAGGCCCTCAGAGATAG |
| *PIN1*_S_2_OT_1_R | GTCCTTCTCTCCTGCCTTGA |
| *PIN1*_S_2_OT_1_DF | ACACTCTTTCCCTACACGACGCTCTTCCGATCTGCGTGAGGAGGAAAGATGG |
| *PIN1*_S_2_OT_1_DR | GTGACTGGAGTTCAGACGTGTGCTCTTCCGATCTGACTGGCTTCACTAGCAGGT |
| *PIN1*_S_2_OT_2_F | TTATGTGGCCGACAGAGTTG |
| *PIN1*_S_2_OT_2_R | TTTTCCTCTCATTCTCTTTGTCC |
| *PIN1*_S_2_OT_2_DF | ACACTCTTTCCCTACACGACGCTCTTCCGATCTGGGGAACCTCAACTGTACTCA |
| *PIN1*_S_2_OT_2_DR | GTGACTGGAGTTCAGACGTGTGCTCTTCCGATCTTCTTTAGCATCCCCACTACCC |
| *PIN1*_S_2_OT_3_F | CCGTGCCCAGTCAATATAAAA |
| *PIN1*_S_2_OT_3_R | AGCTGTCTGTGGGAGCAATTA |
| *PIN1*_S_2_OT_3_DF | ACACTCTTTCCCTACACGACGCTCTTCCGATCTAAACTACAGAGAGCAAAACTCAGGA |
| *PIN1*_S_2_OT_3_DR | GTGACTGGAGTTCAGACGTGTGCTCTTCCGATCTTTGGCACTTGATTTAAGACTGG |
| *PIN1*_S_2_OT_4_F | GGTACCTGGATACTGCCACAG |
| *PIN1*_S_2_OT_4_R | GGCCCCAAAGTTGTTGTTATT |
| *PIN1*_S_2_OT_4_DF | ACACTCTTTCCCTACACGACGCTCTTCCGATCTCGAAACTAAACCATTGTGTGTCC |
| *PIN1*_S_2_OT_4_DR | GTGACTGGAGTTCAGACGTGTGCTCTTCCGATCTAAGCCATAGAGTTTACAGCATCAA |
| *PIN1*_S_2_OT_5_F | AAGCATGCTGGCTGTGGT |
| *PIN1*_S_2_OT_5_R | CTCCAAGTCCGGGTGAGATA |
| *PIN1*_S_2_OT_5_DF | ACACTCTTTCCCTACACGACGCTCTTCCGATCTCTCTCCAGGCCTCGGTTT |
| *PIN1*_S_2_DT_5_DR | GTGACTGGAGTTCAGACGTGTGCTCTTCCGATCTTAGACCACTGCTCCTTTGTGG |
| *PIN1*_S_2_OT_6_F | CTTGGAAATGGTGCCTCAAG |
| *PIN1*_S_2_OT_6_R | CCAGAGCAGAGGATGAAGGA |
| *PIN1*_S_2_OT_6_DF | ACACTCTTTCCCTACACGACGCTCTTCCGATCTCAAAGGTGACCTCCTTTTCC |
| *PIN1*_S_2_OT_6_DR | GTGACTGGAGTTCAGACGTGTGCTCTTCCGATCTATTCTGCTGTCGGCACAGTC |
| *PIN1*_S_2_OT_7_F | TGGTGGGCATCTGTACTCAC |
| *PIN1*_S_2_OT_7_R | CTGAGGATTTGCCAAGAGGT |
| *PIN1*_S_2_OT_7_DF | ACACTCTTTCCCTACACGACGCTCTTCCGATCTGCCTCTCACCTCAAACCCTA |
| *PIN1*_S_2_OT_7_DR | GTGACTGGAGTTCAGACGTGTGCTCTTCCGATCTGTTGCTGAGGACCTGCTGA |
| *PIN1*_S_2_OT_8_F | CTACCTGCTGGGGGCACT |
| *PIN1*_S_2_OT_8_R | CACACTCGTCCAGCAAGACC |
| *PIN1*_S_2_OT_8_DF | ACACTCTTTCCCTACACGACGCTCTTCCGATCTTGAGTGAAACGCAGCACAA |
| *PIN1*_S_2_OT_8_DR | GTGACTGGAGTTCAGACGTGTGCTCTTCCGATCTACAGGGCAGGAGTGAGTCAG |
| *PIN1*_S_2_OT_9_F | GGGAGGCAAGTCATGTGG |
| *PIN1*_S_2_OT_9_R | GGCAGGGCTTCATCCTAGA |

| | |
|---|---|
| *PIN1*_S_2_OT_9_DF | ACACTCTTTCCCTACACGACGCTCTTCCGATCTGGCCCCTGAATAGAATGGAT |
| *PIN1*_S_2_OT_9_DR | GTGACTGGAGTTCAGACGTGTGCTCTTCCGATCTCTCGTCTCCAGAACGCACT |
| *PIN1*_S_2_OT_10_F | GATGCCCAGGCAGAAGTTT |
| *PIN1*_S_2_OT_10_R | TTCCAAATGGGAGAAATTGG |
| *PIN1*_S_2_OT_10_DF | ACACTCTTTCCCTACACGACGCTCTTCCGATCTTCCAGACCCCAGAATGGTAG |
| *PIN1*_S_2_OT_10_DR | GTGACTGGAGTTCAGACGTGTGCTCTTCCGATCTCTGTGTCTCACATCCAGGTCA |
| *PIN1*_AS_2_OT_1_F | CAAGTTAGGGGCAGACATGG |
| *PIN1*_AS_2_OT_1_R | TGGACGTGTCCTAGATGACATT |
| *PIN1*_AS_2_OT_1_DF | ACACTCTTTCCCTACACGACGCTCTTCCGATCTCCAGAGCAGTCCTGGTTAGC |
| *PIN1*_AS_2_OT_1_DR | GTGACTGGAGTTCAGACGTGTGCTCTTCCGATCTCGACTGCCTGTCTGTCTGTCT |
| *PIN1*_AS_2_OT_2_F | AGGCCGTCCACTCTCTCTCT |
| *PIN1*_AS_2_OT_2_R | GGGTCAGACGGGTCACAT |
| *PIN1*_AS_2_OT_2_DF | ACACTCTTTCCCTACACGACGCTCTTCCGATCTCCAGGCCGTCCAGGGAG |
| *PIN1*_AS_2_OT_2_DR | GTGACTGGAGTTCAGACGTGTGCTCTTCCGATCTTCTCCTGGCCCTCGAAGTA |
| *PIN1*_AS_2_OT_3_F | TCCAACACAAAATGATGTTCAA |
| *PIN1*_AS_2_OT_3_R | AAGAAAACCAATACTGTGCACTCA |
| *PIN1*_AS_2_OT_3_DF | ACACTCTTTCCCTACACGACGCTCTTCCGATCTTACAGTAATTATGCAACCCTGGAA |
| *PIN1*_AS_2_OT_3_DR | GTGACTGGAGTTCAGACGTGTGCTCTTCCGATCTATGGAATCTGGACTTGCTATGA |
| *PIN1*_AS_2_OT_4_F | GTGGCGAGTGCTTGTAATCC |
| *PIN1*_AS_2_OT_4_R | CTTCCCTGTCCCTGAATACG |
| *PIN1*_AS_2_OT_4_DF | ACACTCTTTCCCTACACGACGCTCTTCCGATCTAAGCTACCAAAGTTGAGACAAAGA |
| *PIN1*_AS_2_OT_4_DR | GTGACTGGAGTTCAGACGTGTGCTCTTCCGATCTTCTGTCAGAGAGTAAATTGCATTC |
| *PIN1*_AS_2_OT_5_F | CACCACACCTGGCCATAGTT |
| *PIN1*_AS_2_OT_5_R | CAATCAGGTGTCTGCTTTGTTG |
| *PIN1*_AS_2_OT_5_DF | ACACTCTTTCCCTACACGACGCTCTTCCGATCTAGAAGACCCCAGGAGGAGTG |
| *PIN1*_AS_2_DT_5_DR | GTGACTGGAGTTCAGACGTGTGCTCTTCCGATCTAAGGATCCTCTATTCTACAGTTCCTG |
| *PIN1*_AS_2_OT_6_F | CCTCACCACTTCTGCACCTT |
| *PIN1*_AS_2_OT_6_R | GTGCCTCTCTGCCTCCTG |
| *PIN1*_AS_2_OT_6_DF | ACACTCTTTCCCTACACGACGCTCTTCCGATCTCCAGTGAGTCCTGCAGAGG |
| *PIN1*_AS_2_OT_6_DR | GTGACTGGAGTTCAGACGTGTGCTCTTCCGATCTCCAGGGAGGAAGAGCTCAC |
| *PIN1*_AS_2_OT_7_F | TTACAATGCCAAGGGCTCTG |
| *PIN1*_AS_2_OT_7_R | GTAGTGTGCATGGCCTCACG |
| *PIN1*_AS_2_OT_7_DF | ACACTCTTTCCCTACACGACGCTCTTCCGATCTCAGCAGGAAGGTGTGACCTA |
| *PIN1*_AS_2_OT_7_DR | GTGACTGGAGTTCAGACGTGTGCTCTTCCGATCTAATTGTGCCACCAAATCCAA |
| *PIN1*_AS_2_OT_8_F | AAACATTGTATTGTGTCTCAGTTTACC |
| *PIN1*_AS_2_OT_8_R | AACATCTCAGCAAGTAAGAAGAACA |
| *PIN1*_AS_2_OT_8_DF | ACACTCTTTCCCTACACGACGCTCTTCCGATCTTCCAAACCTGCCTGTACCTT |
| *PIN1*_AS_2_OT_8_DR | GTGACTGGAGTTCAGACGTGTGCTCTTCCGATCTCCTTGACTCACACCCCATCT |
| *PIN1*_AS_2_OT_9_F | CAGGAGGTCCCTCTGACTGA |
| *PIN1*_AS_2_OT_9_R | CATGTGGATGCGTGTTAAGG |
| *PIN1*_AS_2_OT_9_DF | ACACTCTTTCCCTACACGACGCTCTTCCGATCTGCTGCACTTCCATTTCAGGT |
| *PIN1*_AS_2_OT_9_DR | GTGACTGGAGTTCAGACGTGTGCTCTTCCGATCTAGGGAAACTGAGGCGAAGA |
| *PIN1*_AS_2_OT_10_F | CTTTCCAAAATTGCCCTGAG |
| *PIN1*_AS_2_OT_10_R | GGCACTTGCTCCCACTCTAA |

| | |
|---|---|
| *PIN1*_AS_2_OT_10_DF | ACACTCTTTCCCTACACGACGCTCTTCCGATCTTACCATAGCATTGGGAGCAG |
| *PIN1*_AS_2_OT_10_DR | GTGACTGGAGTTCAGACGTGTGCTCTTCCGATCTCCCATTTCAGCTATCTGATGATTT |
| *ATXN3*_F1 | TGCACTTTCATTAGCTTACATGC |
| *ATXN3*_R1 | TTTAAACCGAGACCTGGAAAG |
| *ATXN3*_F2 | GATCAAAGGTACTTATTTTTGGTGA |
| *ATXN3*_R2 | AAAATTAGCCAGGCGATGTG |
| *ATXN3*_TP_F | 6'-FAM-CTGCTCTTGCATTCTTTTAATACCAGTGAC |
| *ATXN3*_TP_R | GTTTCGGCGTTACGAGTGGAGTCCTGCTGCTGCTGCTG |
| *ATXN3*_TP_tail | GTTTCGGCGTTACGAGTGGA |

**Supplementary Table S3. *In silico*–predicted off-target candidate sequences corresponding to each target DNA sequence analyzed in this study.**

The sequences of the target sites and their respective off-target candidates, along with their chromosomal locations, are provided. Nucleotide mismatches between the target and each off-target candidate are indicated in red, while the PAM sequences (NGG) within the target and off-target DNA sequences are highlighted in light green.

#### *HEK3*

| Target gene | Target sequence (5' to 3') | Chromosome location |
|---|---|---|
| *HEK3*_Sense_On-target | GGCCCAGACTGAGCACGTGATGG | chr9:107422338 |
| *HEK3*_Sense_OT_1 | caCCCAGACTGAGCACGTGcTGG | chr15:79457571 |
| *HEK3*_Sense_OT_2 | aGCtCAGACTGAGCAaGTGAGGG | chr1:46540026 |
| *HEK3*_Sense_OT_3 | GagCCAGAaTGAGCACGTGAGGG | chr10:129794839 |
| *HEK3*_Sense_OT_4 | GGCCCAGACTcAGCAgGTGtGGG | chr8:142339786 |
| *HEK3*_Sense_OT_5 | GGCCCAGACTGAGCAaaaGAAGG | chr17:4740503 |
| *HEK3*_Sense_OT_6 | GGtCCAGACTGAGCAgcTGATGG | chrX:54951737 |
| *HEK3*_Sense_OT_7 | GGCtCAGACTGAaCACcTGAGGG | chr11:44587638 |
| *HEK3*_Sense_OT_8 | GGCCCAGgCTcAGCACGgGAAGG | chr11:126159755 |
| *HEK3*_Sense_OT_9 | GGCCCAGcCcGAGCAgGTGAGGG | chr19:41905245 |
| *HEK3*_Anti-Sense_On-target | TGTCCTGCGACGCCCTCTGGAGG | chr9:107422383 |
| *HEK3*_Anti-Sense_OT_1 | TGTCCTaCcACGCCCcCTGGTGG | chr17:76715007 |
| *HEK3*_Anti-Sense_OT_2 | aGTCCTGCGgCGaCCTCTGGTGG | chr3:138068757 |
| *HEK3*_Anti-Sense_OT_3 | TGcCCTGtGAaGCCCTCTGGGGG | chr3:154182379 |
| *HEK3*_Anti-Sense_OT_4 | TGTCCTGaGcCaCCCTCTGGAGG | chr5:480191 |
| *HEK3*_Anti-Sense_OT_5 | TGTCCTGCGACtCtCTCaGGGGG | chr17:57642960 |
| *HEK3*_Anti-Sense_OT_6 | TGTCCTGCatCGCCCcCTGGTGG | chr19:28879349 |
| *HEK3*_Anti-Sense_OT_7 | TGcCCTGCGACGCCCTCatGTGG | chr3:184561395 |

#### *VEGFA*

| Target gene | Target sequence (5' to 3') | Chromosome location |
|---|---|---|
| *VEGFA*_Sense_On-target | CAGGCTGCACCCATGGCAGAAGG | chr6:43774346 |
| *VEGFA*_Sense_OT_1 | CAGGCTcCACCCATGtCAGAAGG | chr6:47851289 |
| *VEGFA*_Sense_OT_2 | CtGGCTGCACCCcTGGCAGAGGG | chr11:119188020 |
| *VEGFA*_Sense_OT_3 | CgGGgTGCcCCCATGGCAGAGGG | chr20:43961110 |
| *VEGFA*_Sense_OT_4 | aAGGCTGCtgCCATGGCAGATGG | chr22:47369404 |
| *VEGFA*_Sense_OT_5 | CtGcCTGCACtCATGGCAGAAGG | chr3:7472161 |
| *VEGFA*_Sense_OT_6 | CAGGCTGCACtCtTGaCAGAAGG | chr8:84794367 |
| *VEGFA*_Sense_OT_7 | CcGGCTGaAgCCATGGCAGAAGG | chr8:106257544 |
| *VEGFA*_Sense_OT_8 | CAGGCaGCACCCAgGGCAcAGGG | chr5:16829229 |
| *VEGFA*_Sense_OT_9 | CAGGCaGCAtCaATGGCAGAGGG | chr1:66112609 |
| *VEGFA*_Sense_OT_10 | CtGGaTGCAgCCATGGCAGAGGG | chr7:49819114 |
| *VEGFA*_Anti-Sense_On-target | GGGAACCCCATCCAACAGCCAGG | chr6:43774401 |
| *VEGFA*_Anti-Sense_OT_1 | GGGAACCCCtgCCAACAGCCTGG | chr3:14703181 |
| *VEGFA*_Anti-Sense_OT_2 | GaGAcCCCCAgCCAACAGCCAGG | chr1:234648754 |
| *VEGFA*_Anti-Sense_OT_3 | tGGgACCCCATCCAgCAGCCAGG | chr2:200521394 |
| *VEGFA*_Anti-Sense_OT_4 | GGaAcgCCCATCCAACAGCCTGG | chr3:122264742 |
| *VEGFA*_Anti-Sense_OT_5 | GGGcACCCCAgCCAACAGgCTGG | chr5:173782300 |
| *VEGFA*_Anti-Sense_OT_6 | GGGAACCCCcaCCcACAGCCAGG | chr1:166234468 |
| *VEGFA*_Anti-Sense_OT_7 | GGGAAaCCCATCCAtCAtCCAGG | chr2:19547778 |
| *VEGFA*_Anti-Sense_OT_8 | aGGAACCtCATCaAACAGCCTGG | chr2:141380913 |
| *VEGFA*_Anti-Sense_OT_9 | aGGAACCCCATtCAgCAGCCTGG | chr12:111156619 |
| *VEGFA*_Anti-Sense_OT_10 | aGGAACCCaATCCAACAaCCTGG | chr11:12930431 |

### *PIN1*

| Target gene | Target sequence (5' to 3') | Chromosome location |
|---|---|---|
| *PIN1*_Sense_2_On-target | AGCCAGTGGGAGCGGCCCAGCGG | chr19:9838470 |
| *PIN1*_Sense_2_On-target (Unintended) | AGCCAGTGGGAGCGGCCCAGCGG | chr1:69919449 |
| *PIN1*_Sense_2_OT_2 | AGaCAGTGGGAGCaGCCCAGGGG | chr1:235999447 |
| *PIN1*_Sense_2_OT_3 | AGCCtGTGGGAGCGGtCCAGCGG | chr4:56856766 |
| *PIN1*_Sense_2_OT_4 | tGCCtGTGGGAGgGGCCCAGAGG | chr15:73305045 |
| *PIN1*_Sense_2_OT_5 | gcCCAGTGGGAGgGGCCCAGGGG | chr5:172721310 |
| *PIN1*_Sense_2_OT_6 | AGaCAGTGGctGCGGCCCAGAGG | chr7:72577641 |
| *PIN1*_Sense_2_OT_7 | AaCCAGTGGGAGCtGCCgAGTGG | chr8:666823 |
| *PIN1*_Sense_2_OT_8 | AGCCAGTGGaAGgGGCCCtGTGG | chr8:143562559 |
| *PIN1*_Sense_2_OT_9 | AGCCcGTGGGAGCGGCgCgGCGG | chr7:100965710 |
| *PIN1*_Sense_2_OT_10 | AGCCAGTGaaAGCaGCCCAGTGG | chr6:102784064 |
| *PIN1*_Anti-Sense_2_On-target | ACTGGCTGTGCTTCACCAGCAGG | chr19:9838553 |
| *PIN1*_Anti-Sense_2_OT_1 | ACaGGCTGTGCcTCACCAGCTGG | chr2:95383808 |
| *PIN1*_Anti-Sense_2_OT_2 | tCTGGCTGTGCTTCACCAGgAGG | chr13:110506481 |
| *PIN1*_Anti-Sense_2_OT_3 | AaTGGtTcTGCTTCACCAGCTGG | chr8:114273775 |

| | | |
|---|---|---|
| *PIN1*_Anti-Sense_2_OT_4 | AgTGaCTtTGCTTCACCAGCAGG | chr1:183419456 |
| *PIN1*_Anti-Sense_2_OT_5 | ACTcGaTGTGCTcCACCAGCCGG | chr14:23940861 |
| *PIN1*_Anti-Sense_2_OT_6 | cCTGGCTGTGtTcCACCAGCTGG | chr8:142577793 |
| *PIN1*_Anti-Sense_2_OT_7 | ACTGGgTGTctTTCACCAGCAGG | chr1:2942855 |
| *PIN1*_Anti-Sense_2_OT_8 | ACTGGgaGTGCTTCACCAaCAGG | chr12:129377303 |
| *PIN1*_Anti-Sense_2_OT_9 | ACTGGgTGTGCTTCtCCAGaGGG | chr16:57567670 |
| *PIN1*_Anti-Sense_2_OT_10 | tCTGGCTGTGCgTCAaCAGCTGG | chr9:15117898 |

#### Supplementary Figures

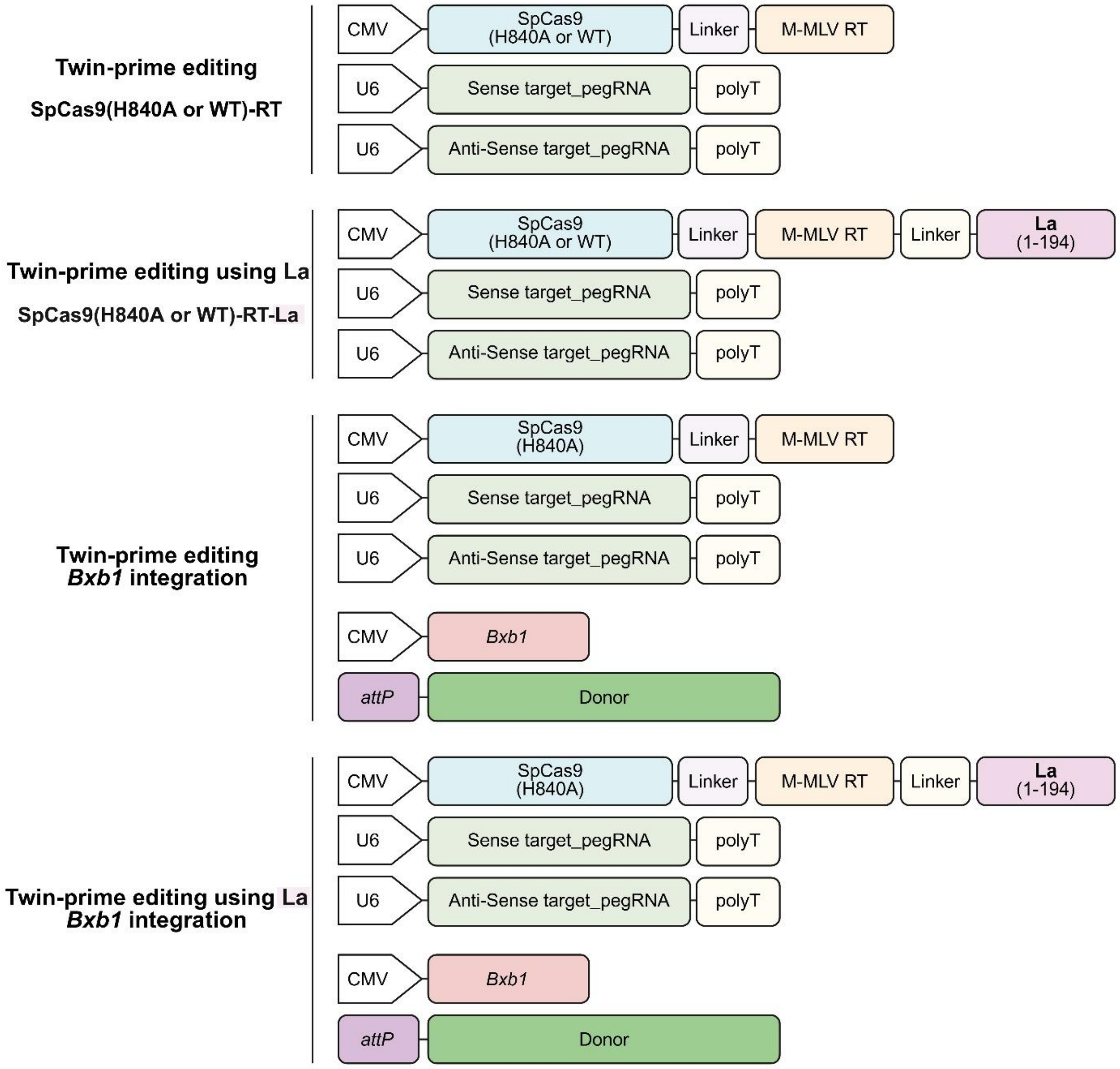


**Supplementary Figure S1. Method for inducing twin prime editing using various prime editor expression plasmid combinations.** To induce target-specific twin prime editing in human-derived cells, expression vectors based on the CMV promoter were constructed for SpCas9(H840A/WT)-RT or La-SpCas9(H840A/WT)-RT. For expression of sense and antisense pegRNAs with target DNA-specific PBS+RTT sequences, U6 promoter-based vectors were also used. For La-SpCas9(H840A/WT)-RT expression, a minimal N-terminus (1–194 aa) of the La protein was fused via a linker to the WT or nickase SpCas9 module. For experiments involving *attB* insertion and subsequent donor DNA recombination via Bxb1 recombinase in human-derived cells, vectors expressing SpCas9(H840A/WT)-RT or La-SpCas9(H840A/WT)-RT were used with the same twin prime editing strategy, along with CMV promoter-based Bxb1 expression vectors and donor DNA vectors containing the *attP* recognition sequence. MMLV-RT: Reverse transcriptase domain from Moloney Murine Leukemia Virus; pegRNA: prime editing guide RNA; La (1–194): La protein N-terminal domain (1–194 aa); Bxb1: Bxb1 recombinase; *attB*/*P*: Bxb1 recognition DNA sequences.

#### *AAVS1* target site

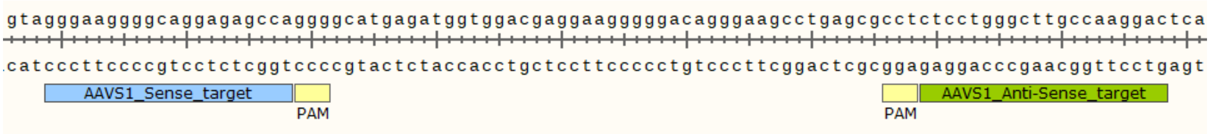


#### *AAVS1 attB* insertion

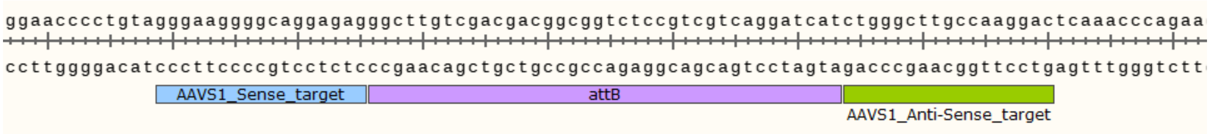


#### *AAVS1 attP* insertion

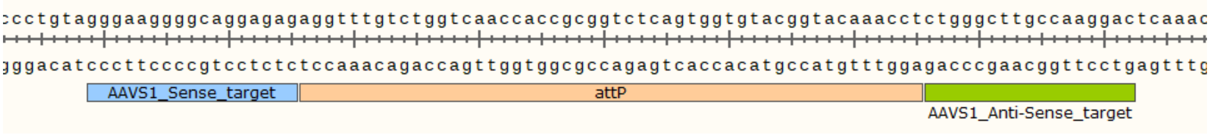


#### *HEK3* target site

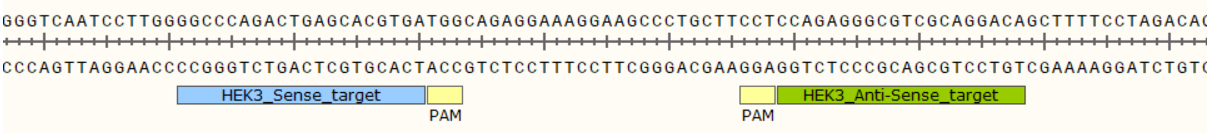


#### *HEK3 attB* insertion

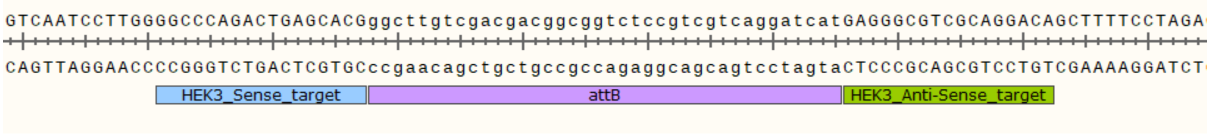


#### *FANCF* target site

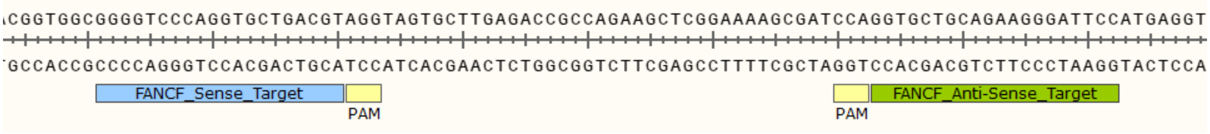


#### *FANCF attB* insertion

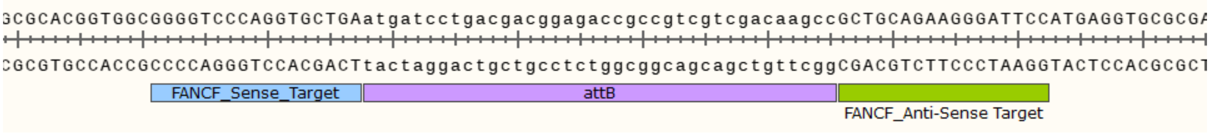


#### *CCR5* target site

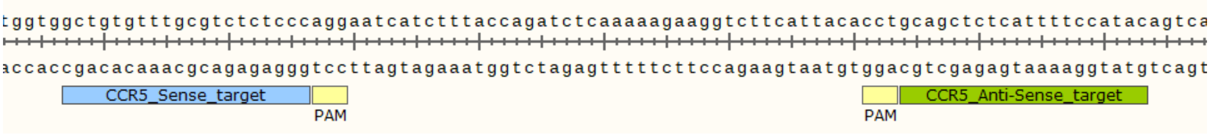


#### *CCR5 attB* insertion

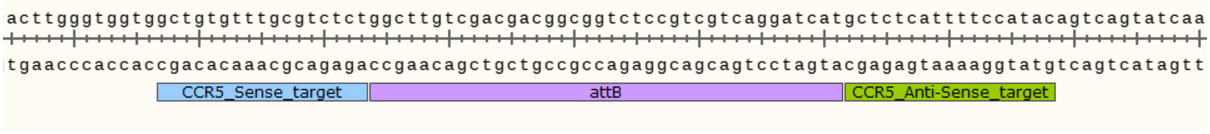


#### *EMX1_1* target stie

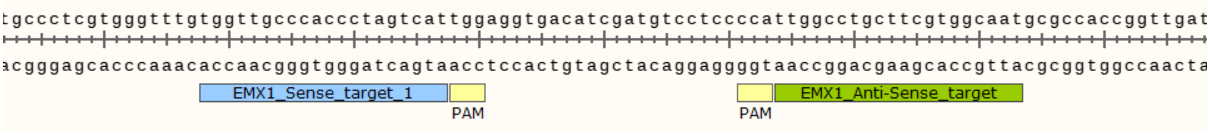


#### *EMX1_1 attB* insertion

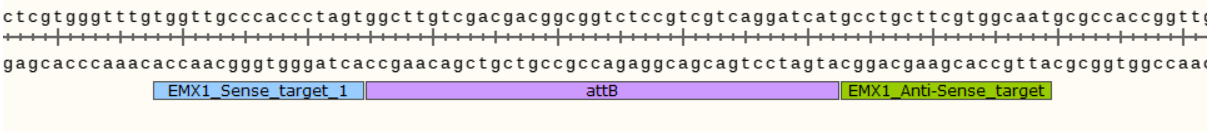


#### *EMX1_2* target site

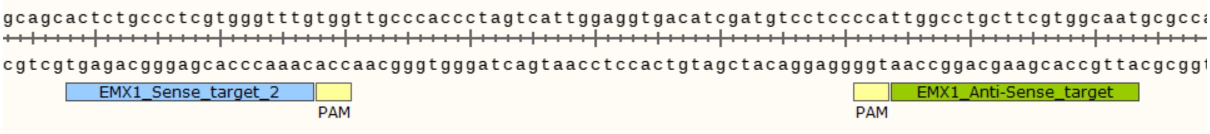


#### *EMX1_2 attB* insertion

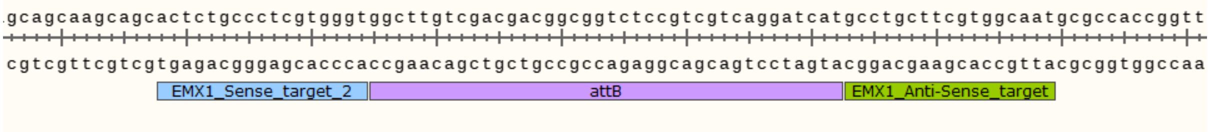


#### *VEGFA* target site

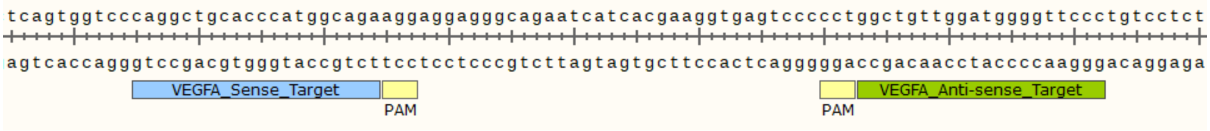


#### *VEGFA attB* insertion

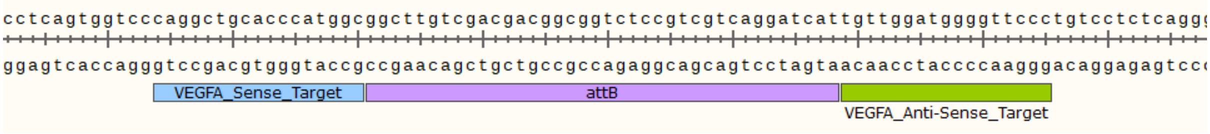


#### *DAPK1* target stie

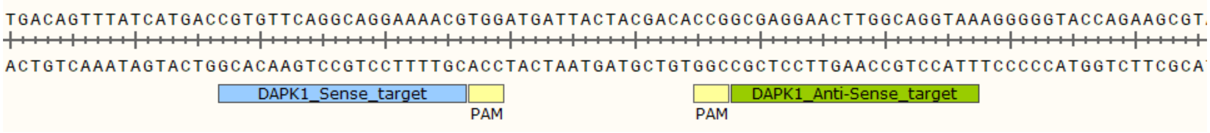


#### *DAPK1 attB* insertion

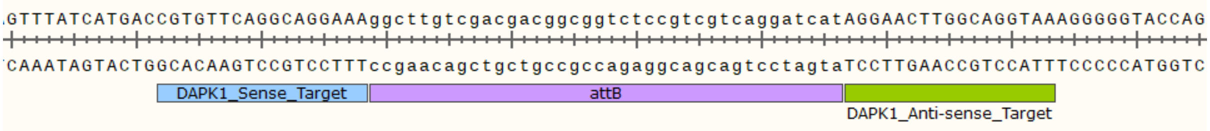


#### *PIN1_1* target site

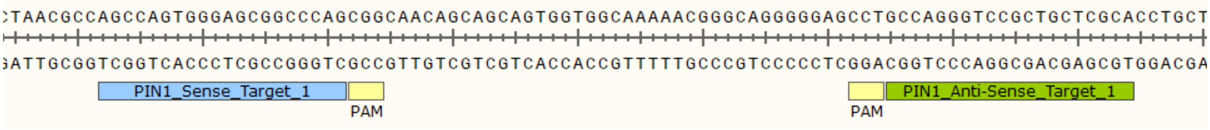


#### *PIN1_1 attB* insertion

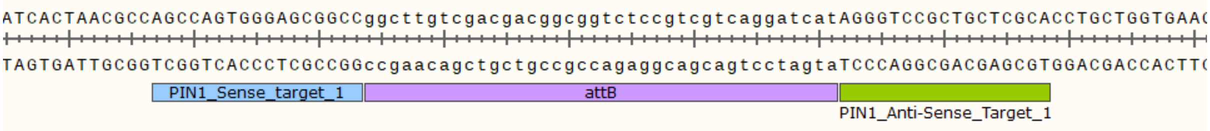


#### *PIN1_2* target site

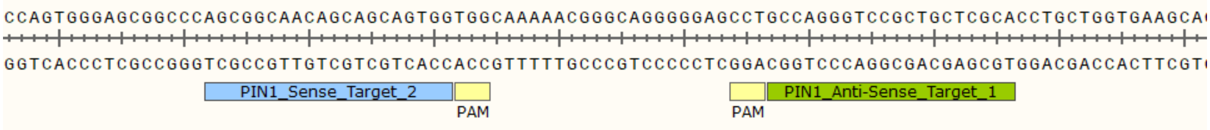


#### *PIN1_2 attB* insertion

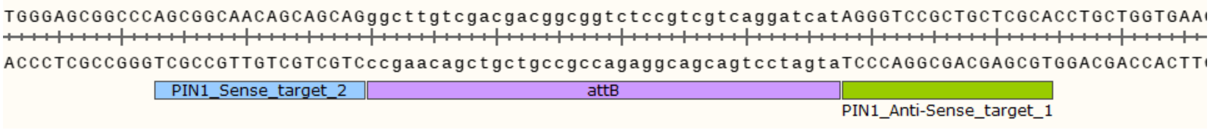


#### *PIN1_3* target site

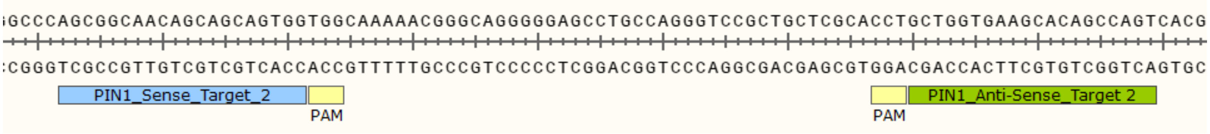


***PIN1_3 attB* insertion**

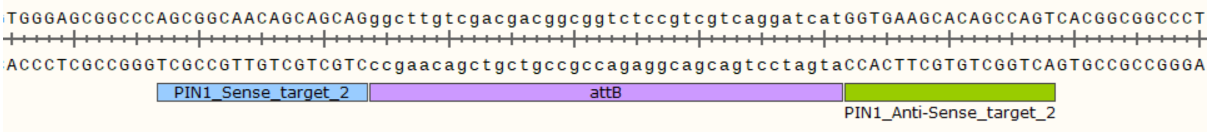


***GAPDH_1* target site**

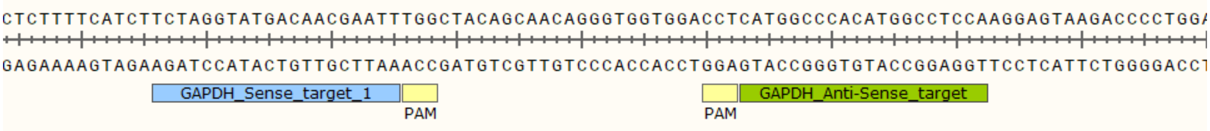


***GAPDH_1 attB* insertion**

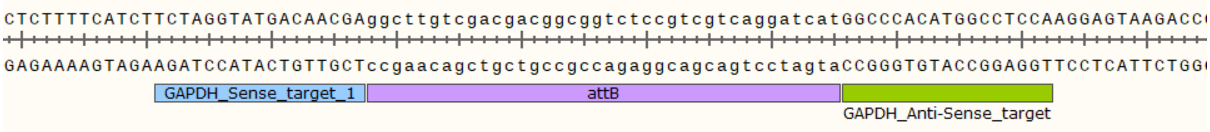


***GAPDH_2* target site**

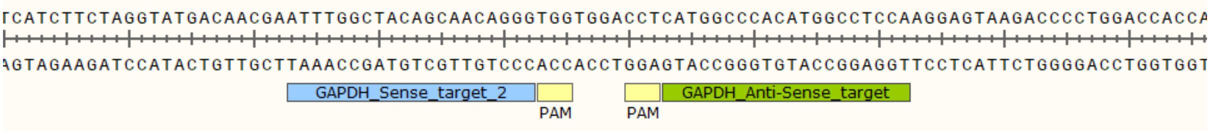


***GAPDH_2 attB* insertion**

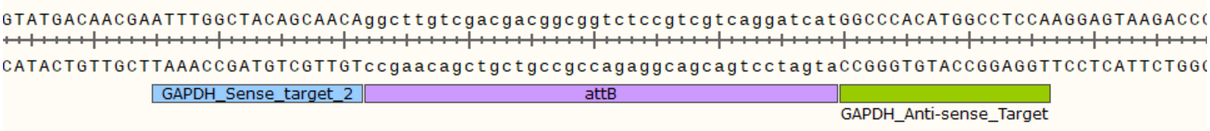


**Supplementary Figure S2. Twin prime editing strategy using paired pegRNA based on SpCas9(H840A/WT)-RT or La-SpCas9(H840A/WT)-RT.** Each DNA schematic illustrates the design of paired pegRNAs targeting specific sequences within the indicated genomic loci (*AAVS1, HEK3, FANCF, CCR5, EMX1, VEGFA, DAPK1, PIN1*, and *GAPDH*) and the precise insertion of *attB/attP* sequences achieved via SpCas9(H840A/WT)-RT or La-SpCas9(H840A/WT)-RT. Within the target DNA sequences, the protospacers and PAM (NGG) sequences corresponding to each sense and antisense pegRNA are colored in light blue, light green, and yellow, respectively.

### *AAVS1* Twin-prime editing pattern (*attB* insertion)

**AAVS1**

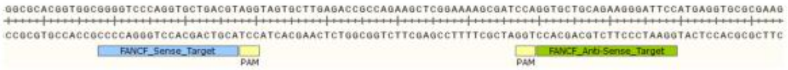


Insertion
Subtitution
Deletion : -----

**Normal control** GGGAAGGGGCAGGAGAGCCAGGGGCATGAGATGGTGGACGAGGAAGGGGGACAGGGAAGCCTGAGCGCCTCTCCTGGGCTTGCCAAGGAC

#### *AAVS1* / SpCas9(H840A)-RT / Whole overlap (*attB* insertion)

Insertion
Subtitution
Deletion : -----

| | | |
|---|---|---|
| Precise insertion | GGGAAGGGGCAGGAGAGggcttgtcgacgacggcggtctccgtcgtcaggatcatCTGGGCTTGCCAAGGAC<br>*attB* (38bp) | 19.11 % |
| Pure indel | GGGAAGGGGCAGGAGAG------------------------------------------------------------------CTGGGCTTGCCAAGGAC | 0.29 % |
|  | GGGAAGGGGCAG-------------GAGATGGTGGACGAGGAAGGGGGACAGGGAAGCCTGAGCGCCTCTCCTGGGCTTGCCAAGGAC | 0.16 % |
|  | GGGA--------------AGGGGCATGAGATGGTGGACGAGGAAGGGGGACAGGGAAGCCTGAGCGCCTCTCCTGGGCTTGCCAAGGAC | 0.02 % |
| Partial insertion | GGGAAGGGGCAGGAGAGGGCTTggcttgtcgacgacggcggtctccgtcgtcaggatcatCTGGGCTTGCCAAGGAC | 0.26 % |
|  | GGGAAGGGGCAGGAGAGggcttgtcgacgacggcggtAC-----------------------------TGGTGGACGAGGAAGGGGGACAGGGAAGCCTGAGCGCCTCTCCTGGGCTTGCCAAGGAC | 0.18 % |
| | GGGAAGGGGCAGGAGAGCgcttgtcgacgacggcggtctccgtcgtcaggatcatCTGGGCTTGCCAAGGAC | 0.12 % |
| | GGGAAGGGGCAGGAGAGCgcttgtcgacgacggcggtctccgtcgtcaggatcTCCTGGGCTTGCCAAGGAC | 0.12 % |
|  | GGGAAGGGGCAGGAGAGggcttgtcgacgacggcggtctccgtcgtcaggatc----------TGAGATGGTGGACGAGGAAGGGGGACAGGGAAGCCTGAGCGCCTCTCCTGGGCTTGCCAAGGAC | 0.12 % |

#### *AAVS1* / La-SpCas9(H840A)-RT / Whole overlap (*attB* insertion)

Insertion
Subtitution
Deletion : -----

| | | |
|---|---|---|
| Precise insertion | GGGAAGGGGCAGGAGAGggcttgtcgacgacggcggtctccgtcgtcaggatcatCTGGGCTTGCCAAGGAC<br>*attB* (38bp) | 24.10 % |
| Pure indel | None | |
| Partial insertion | GGGAAGGGGCAGGAGAGggcttgtcgacgacggc--------------------ATCTGGGCTTGCCAAGGAC | 0.55 % |
| | GGGAAGGGGCAGGAGAG-gcttgtcgacgacggcggtctccgtcgtcaggatcatCTGGGCTTGCCAAGGAC | 0.24 % |
|  | GGGAAGGGGCAGGAGAGggcttgt-----------------------------------------------------------cgtcaggatcatCTGGGCTTGCCAAGGAC<br>Sense insertion / Anti-Sense insertion | 0.18 % |
|  | GGGAAGGGGCAGGAGAGCCAGGGGCA--------------------------------------------GGAGAGggcttgtcgacgacggcggtctccgtcgtcaggatcatCTGGGCTTGCCAAGGAC | 0.18 % |
|  | GGGAAGGGGCAGGAGAGggcttgtcgacgac-----------------------GCATGAGATGGTGGACGAGGAAGGGGGACAGGGAAGCCTGAGCGCCTCTCCTGGGCTTGCCAAGGAC | 0.13 % |

#### *AAVS1* / SpCas9(WT)-RT / Whole overlap (*attB* insertion)

Insertion
Subtitution
Deletion : -----

| | | |
|---|---|---|
| Precise insertion | GGGAAGGGGCAGGAGAGggcttgtcgacgacggcggtctccgtcgtcaggatcatCTGGGCTTGCCAAGGAC<br>*attB* (38bp) | 10.35 % |
| Pure indel | GGGAAGGGGCAGGAGAG--------------------------------------------------------------CTGGGCTTGCCAAGGAC | 1.18 % |
|  | GGGAAGGGGCAGGA------------------------------------------------------------------TCATCTGGGCTTGCCAAGGAC | 1.57 % |
|  | GGGAAGGGGCAGGAGAG--------------------------------------------------------------CCTGGGCTTGCCAAGGAC | 0.52 % |
| | GGGAAGGGGCAGGAGAGCCAGGGGCATGAGATGGTGGACGAGGAAGGGGGACAGGGAAGCCTGAGCGCCTCTCCATCTGGGCTTGCCAAGGAC | 0.48 % |
| | GGGAAGGGGCAGGAGAGCCAGGGGCATGAGATGGTGGACGAGGAAGGGGGACAGGGAAGCCTGAGCGCCTCTCCCTGGGCTTGCCAAGGAC | 0.43 % |
| Partial insertion | GGGAAGGGGCAGGAGAGggcttgtcgacgacggcggtctccgtcgtcaggatcatCCAGGGGCATGAGATGGTGGACGAGGAAGGGGGACAGGGAAGCCTGAGCGCCTCTCCTGGGCTTGCCAAGGAC | 1.30 % |
|  | GGGAAGGGGCAGGAGAGggcttgtcgacgac-----------------------CCAGGGGCATGAGATGGTGGACGAGGAAGGGGGACAGGGAAGCCTGAGCGCCTCTCCTGGGCTTGCCAAGGAC | 0.73 % |
| | GGGAAGGGGCAGGAGAGCCAGGGGCATGAGATGGTGGACGAGGAAGGGGGACAGGGAAGCCTGAGCGCCTCTCCggcttgtcgacgacggcggtctccgtcgtcaggatcatCTGGGCTTGCCAAGGAC | 0.53 % |
|  | GGGAAGGGGCAGGAGAGggcttg-------------------------------TCCTGGGCTTGCCAAGGAC | 0.41 % |
| | GGGAAGGGGCAGGAGAggcttgtcgacgacggcggtctccgtcgtcaggatcatGCCAGGGGCATGAGATGGTGGACGAGGAAGGGGGACAGGGAAGCCTGAGCGCCTCTCCTGGGCTTGCCAAGGAC | 0.37 % |

#### *AAVS1* / La-SpCas9(WT)-RT / Whole overlap (*attB* insertion)

Insertion
Subtitution
Deletion : -----

| | Sequence | % |
|---|---|---|
| Precise insertion | GGGAAGGGGCAGGAGAGggcttgtcgacgacggcggtctccgtcgtcaggatcatCTGGGCTTGCCAAGGAC (*attB* (38bp)) | 28.95 % |
| Pure indel | GGGAAGGGGCAGGAGAGGGCT------------------------------------------------------------CTGGGCTTGCCAAGGAC | 0.75 % |
|  | GGGAAGGGGCAGGAGAG-------------------GCAT-----------------------------------------CTGGGCTTGCCAAGGAC | 0.51 % |
| | GGGAAGGGGCAGGAGAGCCAGGGGCATGAGATGGTGGACGAGGAAGGGGGACAGGGAAGCCTGAGCGCCTCTCCCTGGGCTTGCCAAGGAC | 0.43 % |
| | GGGAAGGGGCAGGAGAGGCCAGGGGCATGAGATGGTGGACGAGGAAGGGGGACAGGGAAGCCTGAGCGCCTCTCCTGGGCTTGCCAAGGAC | 0.40 % |
| | GGGAAGGGGCAGGAGAGCCAGGGGCATGAGATGGTGGACGAGGAAGGGGGACAGGGAAGCCTGAGCGCCTCTCCTCTGGGCTTGCCAAGGAC | 0.29 % |
| Partial insertion | GGGAAGGGGCAGGAGAGggcttgtcga----------------------------CTGGGCTTGCCAAGGAC | 0.84 % |
|  | GGGAAGGGGCAGGAGAGggcttgtcgacgacggcggtctccg--------------TGGGCTTGCCAAGGAC | 0.83 % |
| | GGGAAGGGGCAGGAGAGggcttgtcgacgacggcggtctccgtcgtcaggatcatCCAGGGGCATGAGATGGTGGACGAGGAAGGGGGACAGGGAAGCCTGAGCGCCTCTCCTGGGCTTGCCAAGGAC | 0.70 % |
|  | GGGAAGGGGCAGGAGAGggct-----------gcggtctccgtcgtcaggatcatCTGGGCTTGCCAAGGAC | 0.54 % |
|  | GGGAAGGGGCAGGAGA-------gtcgacgacggcggtctccgtcgtcaggatcatCTGGGCTTGCCAAGGAC | 0.53 % |

#### *AAVS1* / SpCas9(H840A)-RT / Homology arm (*attB* insertion)

Insertion
Subtitution
Deletion : -----

| | Sequence | % |
|---|---|---|
| Precise insertion | GGGAAGGGGCAGGAGAGggcttgtcgacgacggcggtctccgtcgtcaggatcatCTGGGCTTGCCAAGGAC (*attB* (38bp)) | 5.47 % |
| Pure indel | GGGAAGGGGCAGGAGAGCC-GGGGCATGAGATGGTGGACGAGGAAGGGGGACAGGGAAGCCTGAGCGCCTCTCCTGGGCTTGCCAAGGAC | 0.08 % |
| | GGGAAGGGGCAGGAGAG-CAGGGGCATGAGATGGTGGACGAGGAAGGGGGACAGGGAAGCCTGAGCGCCTCTCCTGGGCTTGCCAAGGAC | 0.03 % |
|  | GGGAAGGGGCAGGAGAG---------------------------------------------------------------CTGGGCTTGCCAAGGAC | 0.02 % |
| Partial insertion | GGGAAGGGGCAGGAGAGggcttgtcgacgaTggcggtctccgtcgtcaggatcatCTGGGCTTGCCAAGGAC | 1.63 % |
| | GGGAAGGGGCAGGAGAGggcCtgtcgacgacggcggtctccgtcgtcaggatcatCTGGGCTTGCCAAGGAC | 0.37 % |
| | GGGAAGGGGCAGGAGAGggcttgtcgacgacggcggtctccgtcgtcagCCtcTCCTGGGCTTGCCAAGGAC | 0.11 % |
| | GGGAAGGGGCAGGAGAGGAAGGGGCAGGAGAGggcttgtcgacgacggcggtctccgtcgtcaggatcatCTGGGCTTGCCAAGGAC | 0.11 % |
| | GGGAAGGGGCAGGAGAGggcttgtcgacgaTggcggtctccgtcgtc---atcatCTGGGCTTGCCAAGGAC | 0.07 % |

#### *AAVS1* / La-SpCas9(H840A)-RT / Homology arm (*attB* insertion)

Insertion
Subtitution
Deletion : -----

| | Sequence | % |
|---|---|---|
| Precise insertion | GGGAAGGGGCAGGAGAGggcttgtcgacgacggcggtctccgtcgtcaggatcatCTGGGCTTGCCAAGGAC (*attB* (38bp)) | 9.04 % |
| Pure indel | GGGAAGGGGCAGGAGAG---------------------------------------------------------------CTGGGCTTGCCAAGGAC | 0.21 % |
|  | GGGAAGGGGCAGGAGAG---------------------------------------------------------------GCCTCCCCTCTGGGCTTGCCAAGGAC | 0.08 % |
| Partial insertion | GGGAAGGGGCAGGAGAGggcttgtcgacgacggcggtctccgtcgtcGggatcatCTGGGCTTGCCAAGGAC | 1.10 % |
| | GGGAAGGGGCAGGAGAGggcttgtcgacgaTggcggtctccgtcgtcaggatcatCTGGGCTTGCCAAGGAC | 0.84 % |
| | GGGAAGGGGCAGGAGAGggcCtgtcgacgacggcggtctccgtcgtcaggatcatCTGGGCTTGCCAAGGAC | 0.47 % |
| | GGGAAGGGGCAGGAGAGggcttgtcgacgGcggcggtctccgtcgtcaggatcatCTGGGCTTGCCAAGGAC | 0.34 % |
| | GGGAAGGGGCAGGAGAGggcttgtcgacgGcggcggtctccgtcgtcGggatcatCTGGGCTTGCCAAGGAC | 0.15 % |

#### *AAVS1* / SpCas9(WT)-RT / Homology arm (*attB* insertion)

Insertion
Subtitution
Deletion : -----

| | Sequence | % |
|---|---|---|
| Precise insertion | GGGAAGGGGCAGGAGAGggcttgtcgacgacggcggtctccgtcgtcaggatcatCTGGGCTTGCCAAGGAC (*attB* (38bp)) | 5.18 % |
| Pure indel | GGGAAGGGGCAGGAGAG---------------------------------------------------------------CTGGGCTTGCCAAGGAC | 7.56 % |
| | GGGAAGGGGCAGGAGAGCCAGGGGCATGAGATGGTGGACGAGGAAGGGGGACAGGGAAGCCTGAGCGCCTCTCCCTGGGCTTGCCAAGGAC | 1.45 % |
| | GGGAAGGGGCAGGAGAGGCCAGGGGCATGAGATGGTGGACGAGGAAGGGGGACAGGGAAGCCTGAGCGCCTCTCCTGGGCTTGCCAAGGAC | 1.04 % |
|  | GGGAAGGGGCAGGAGA----------------------------------------------------------------CTGGGCTTGCCAAGGAC | 0.55 % |
| | GGGAAGGGGCAGGAGAGAGCCAGGGGCATGAGATGGTGGACGAGGAAGGGGGACAGGGAAGCCTGAGCGCCTCTCCTGGGCTTGCCAAGGAC | 0.52 % |
| Partial insertion | GGGAAGGGGCAGGAGAG--------------ggcggtctccgtcgtcaggatcatCTGGGCTTGCCAAGGAC | 0.47 % |
| | GGGAAGGGGCAGGAGAGGAGggcttgtcgacgacggcggtctccgtcgtcaggatcatCTGGGCTTGCCAAGGAC | 0.34 % |
|  | GGGAAGGGGCAGGAGAGggcttgtcgacgacggcggtctccgtcgtca-------CTGGGCTTGCCAAGGAC | 0.31 % |
|  | GGGAAGGGGCAGGAGAGggctt---------------------------------CTGGGCTTGCCAAGGAC | 0.30 % |
| | GGGAAGGGGCAGGAGAGggcttgtcgacgaTggcggtctccgtcgtcaggatcatCTGGGCTTGCCAAGGAC | 0.29 % |

#### *AAVS1* / La-SpCas9(WT)-RT / Homology arm (*attB* insertion)

Insertion
Subtitution
Deletion : -----

| | Sequence | % |
|---|---|---|
| **Precise insertion** | GGGAAGGGGCAGGAGAGggcttgtcgacgacggcggtctccgtcgtcaggatcatCTGGGCTTGCCAAGGAC<br>*attB* (38bp) | 4.14 % |
| **Pure indel** | GGGAAGGGGCAGGAGAG------------------------------------------------------------------CTGGGCTTGCCAAGGAC | 18.25 % |
| | GGGAAGGGGCAGGAGAGGCCAGGGGCATGAGATGGTGGACGAGGAAGGGGGACAGGGAAGCCTGAGCGCCTCTCCTGGGCTTGCCAAGGAC | 1.17 % |
| | GGGAAGGGGCAGGAGAGCCAGGGGCATGAGATGGTGGACGAGGAAGGGGGACAGGGAAGCCTGAGCGCCTCTCCCTGGGCTTGCCAAGGAC | 0.86 % |
|  | GGGAAGGGGCAGGAGAGAT-------------------------------------------------------------------CTGGGCTTGCCAAGGAC | 0.77 % |
|  | GGGAAGGGGCAGGAGAGCCAG-----------------------------------------------------------------CTGGGCTTGCCAAGGAC | 0.63 % |
| **Partial insertion** | GGGAAGGGGCAGGAGAGggcttgtcg----------------------------CTGGGCTTGCCAAGGAC | 0.82 % |
|  | GGGAAGGGGCAGGAGAGggcttgtcgacgacggcggtctccgtcgtcag------CTGGGCTTGCCAAGGAC | 0.42 % |
| | GGGAAGGGGCAGGAGAGggcttgtcgacgaTggcggtctccgtcgtcaggatcatCTGGGCTTGCCAAGGAC | 0.24 % |
| | GGGAAGGGGCAGGAGAGGAAGGGGCAGGAGAGggcttgtcgacgacggcggtctccgtcgtcaggatcatCTGGGCTTGCCAAGGAC | 0.18 % |
| | GGGAAGGGGCAGGAGAGAAGGGGCAGGAGAGggcttgtcgacgacggcggtctccgtcgtcaggatcatCTGGGCTTGCCAAGGAC | 0.17 % |

#### *AAVS1* / SpCas9(H840A)-RT / Partial overlap (*attB* insertion)

Insertion
Subtitution
Deletion : -----

| | Sequence | % |
|---|---|---|
| **Precise insertion** | GGGAAGGGGCAGGAGAGggcttgtcgacgacggcggtctccgtcgtcaggatcatCTGGGCTTGCCAAGGAC<br>*attB* (38bp) | 12.66 % |
| **Pure indel** | GGGAAGGGGCAGGAGAG------------------------------------------------------------------CTGGGCTTGCCAAGGAC | 0.08 % |
| | GGGAAGGGGCAGGAGAGGGCTCCAGGGGCATGAGATGGTGGACGAGGAAGGGGGACAGGGAAGCCTGAGCGCCTCTCCTGGGCTTGCCAAGGAC | 0.08 % |
| **Partial insertion** | GGGAAGGGGCAGGAGAGggcttgtcgacgacggcggtctc---cgtcaggatcatCTGGGCTTGCCAAGGAC | 0.24 % |
|  | GGGAAGGGGCAGGAGAGggcttgtcgacga-------------tcgtcaggatcatCTGGGCTTGCCAAGGAC | 0.22 % |
| | GGGAAGGGGCAGGAGAGggcttgt---cgacggcggtctccgtcgtcaggatcatCTGGGCTTGCCAAGGAC | 0.19 % |
|  | GGGAAGGGGCAGGAGAG--------------ggcggtctccgtcgtcaggatcatCTGGGCTTGCCAAGGAC | 0.17 % |
| | GGGAAGGGGCAGGAGAGggcttgGcgacgacggcggtctccgtcgtcaggatcatCTGGGCTTGCCAAGGAC | 0.16 % |

#### *AAVS1* / La-SpCas9(H840A)-RT / Partial overlap (*attB* insertion)

Insertion
Subtitution
Deletion : -----

| | Sequence | % |
|---|---|---|
| **Precise insertion** | GGGAAGGGGCAGGAGAGggcttgtcgacgacggcggtctccgtcgtcaggatcatCTGGGCTTGCCAAGGAC<br>*attB* (38bp) | 23.77 % |
| **Pure indel** | None | |
| **Partial insertion** | GGGAAGGGGCAGGAGAGggcttgtcgacgacggcggtctc---cgtcaggatcatCTGGGCTTGCCAAGGAC | 1.09 % |
| | GGGAAGGGGCAGGAGAGggcttgtcgacgacggcgGTCTCCgtctccgtcgtcaggatcatCTGGGCTTGCCAAGGAC | 1.02 % |
|  | GGGAAGGGGCAGGAGAGggcttgtcgacgacggcggtc--------------CCAGGGGCATGAGATGGTGGACGAGGAAGGGGGACAGGGAAGCCTGAGCGCCTCTCCTGGGCTTGCCAAGGAC | 0.50 % |
|  | GGGAAGGGGCAGGAGAGggcttgtcgacgacggcggtctccgtcgtGca-----------------------gtctccgtcgtcaggatcatCTGGGCTTGCCAAGGAC<br>Sense insertion / Anti-Sense insertion | 0.30 % |
| | GGGAAGGGGCAGGAGAGggcttgGcgacgacggcggtctccgtcgtcaggatcatCTGGGCTTGCCAAGGAC | 0.25 % |

##### *AAVS1* / SpCas9(WT)-RT / Partial overlap (*attB* insertion)

Insertion
Subtitution
Deletion : -----

**Precise insertion** GGGAAGGGGCAGGAGAGggcttgtcgacgacggcggtctccgtcgtcaggatcatCTGGGCTTGCCAAGGAC 8.21 %
*attB* (38bp)

**Pure indel**
GGGAAGGGGCAGGAGAG---------------------------------------------------------CTGGGCTTGCCAAGGAC 0.79 %
GGGAAGGGGCAGGAGAGCCAGGGGCATGAGATGGTGGACGAGGAAGGGGACAGGGAAGCCTGAGCGCCTCTCCCTGGGC TTGCCAAGGAC 0.49 %
GGGAAGGGGCAGGAGAGCCAGGGGCATGAGATGGTGGACGAGGAAGGGGACAGGGAAGCCTGAGCGCCTCTCCCATCTGGG CTTGCCAAGGAC 0.30 %
GGGAAGGGGCAGGAGAGCCAGGGGCATGAGATGGTGGACGAGGAAGGGGACAGGGAAGCCTGAGCGCCTCTCATCTGGGC TTGCCAAGGAC 0.19 %
GGGAAGGGGCAGGAGAGCCAGGGGCATGAGATGGTGGACGAGGAAGGGGACAGGGAAGCCTGAGCGCCTCTCCTCTGGGC TTGCCAAGGAC 0.17 %

**Partial insertion**
GGGAAGGGGCAGGAGAGggcttgtcgacgacggcggtctccgtcgt---------CCAGGGGCATGAGATGGTGGACGAGGAAGGGGACAGGG AAGCCTGAGCGCCTCTCCTGGGCTTGCCAAGGAC 1.22 %
GGGAAGGGGCAGGAGAG-------------------tctccgtcgtcaggatcatCTGGGCTTGCCAAGGAC 0.84 %
GGGAAGGGGCAGGAGAGggcttgtcgacgacggcggt-----------------CCAGGGGCATGAGATGGTGGACGAGGAAGGGGACAGGGAAG CCTGAGCGCCTCTCCTGGGCTTGCCAAGGAC 0.79 %
GGGAAGGGGCAGGAGAGCCAGGGGCATGAGATGGTGGACGAGGAAGGGGACAGGGAAGCCTGAGCGCCTCTC--------Cacgac ggcggtctccgtcgtcaggatcatCTGGGCTTGCCAAGGAC 0.51 %
GGGAAGGGGCAGGAGAGggcttgtcgacgacggcggtctc-------------CCAGGGGCATGAGATGGTGGACGAGGAAGGGGACAGGGAA GCCTGAGCGCCTCTCCTGGGCTTGCCAAGGAC 0.49 %

##### *AAVS1* / La-SpCas9(WT)-RT / Partial overlap (*attB* insertion)

Insertion
Subtitution
Deletion : -----

**Precise insertion** GGGAAGGGGCAGGAGAGggcttgtcgacgacggcggtctccgtcgtcaggatcatCTGGGCTTGCCAAGGAC 7.32 %
*attB* (38bp)

**Pure indel**
GGGAAGGGGCAGGAGAGGAT-------------------------------------------------CTGGGCTTGCCAAGGAC 0.98 %
GGGAAGGGGCAGGAGAG-------------------------------------------------CTGGGCTTGCCAAGGAC 0.94 %
GGGAAGGGGCAGGAGAG---------------------------------------------------TCATCTGGGCTTGCCAAGGAC 0.72 %
GGGAAGGGGCAGGAGAG---------------------GAGGA------------------------TCATCTGGGCTTGCCAAGGAC 0.69 %

**Partial insertion**
GGGAAGGGGCAGGAGAGggcttgtcgacgacggcggtctccgtcgt---------CCAGGGGCATGAGATGGTGGACGAGGAAGGGGACAGGG AAGCCTGAGCGCCTCTCCTGGGCTTGCCAAGGAC 0.86 %
GGGAAGGGGCAGGAGAGggcttgtcgacgacggcggtctccgtcg---------tGCTGCAGAAGGGATTCC 0.61 %
GGGAAGGGGCAGGAGAGggcttgtcgacgacggcggt---cgtcaggatcatGCTGCAGAAGGGATTCC 0.53 %
GGGAAGGGGCAGGAGAG-------------------------Ccgtcaggatcat GCTGCAGAAGGGATTCC 0.41 %
GGGAAGGGGCAGGAGAGggcttgt---------cggtctccgtcgtcaggatcatCTGGGCTTGCCAAGGAC 0.39 %

### *AAVS1* Twin-prime editing pattern (*attP* insertion)

**AAVS1**

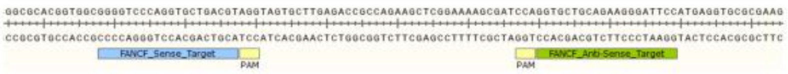


Insertion
Subtitution
Deletion : -----

**Normal control** GGGAAGGGGCAGGAGAGCCAGGGGCATGAGATGGTGGACGAGGAAGGGGACAGGGAAGCCTGAGCGCCTCTCCTGGGC TTGCCAAGGAC

##### *AAVS1* / SpCas9(H840A)-RT / Whole overlap (*attP* insertion)

Insertion
Subtitution
Deletion : -----

**Precise insertion** GGGAAGGGGCAGGAGAGaggtttgtctggtcaaccaccgcggtctcagtggtgtacggtacaaacctCTGGGCTTGCCAAGGAC 13.41 %
*attP* (50bp)

**Pure indel**
GGGAAGGGGCAGGAGAG---------------------------GAA-----------------------------------CTGGGCTTGCCAAGGAC 0.55 %
GGGA---------------AGGGGCATGAGATGGTGGACGAGGAAGGGGACAGGGAAGCCTGAGCGCCTCTCCTGGGCTTGCCAAGGAC 0.10 %

**Partial insertion**
GGGAAGGGGCAGGAGAGCggtttgtctggtcaaccaccgcggtctcagtggtgtacggtacaaacctCTGGGCTTGCCAAGGAC 0.30 %
GGGAAGGGGCAGGAGAG-------------TccaccgcggtctcagtggtgtacggtacaaacctCTGGGCTTGCCAAGGAC 0.17 %
GGGAAGGGGCAGGAGAGaggtttgtctggtcGaccaccgcggtctcagtggtgtacggtacaaacctCTGGGCTTGCCAAGGAC 0.15 %
GGGAAGGGGCAGGAGTGaggtttgtctggtcaaccaccgcggtctcagtggtgtacggtacaaacctCTGGGCTTGCCAAGGAC 0.10 %
GGGAAGGGGCAGGAGAGaggtttgtctggtcaaccCcgcggtctcagtggtgtacggtacaaacctCTGGGCTTGCCAAGGAC 0.08 %

#### *AAVS1* / La-SpCas9(H840A)-RT / Whole overlap (*attP* insertion)

Insertion
Subtitution
Deletion : -----

| | | |
|---|---|---|
| Precise insertion | GGGAAGGGGCAGGAGAGaggtttgtctggtcaaccaccgcggtctcagtggtgtacggtacaaacctCTGGGCTTGCCAAGGAC<br>*attP* (50bp) | 20.72 % |
| Pure indel | None | |
| Partial insertion | GGGAAGGGGCAGGAGTGaggtttgtctggtcaaccaccgcggtctcagtggtgtacggtacaaacctCTGGGCTTGCCAAGGAC | 0.12 % |
|  | GGGAAGGGGCAGGAGAGaggtttgtctggtcaaccaccgcggtctcagtggtgtacggtac-------------------------------------CAGGGAAGCCTGAGCGCCTCTCCTGGGCTTGCCAAGGAC | 0.12 % |
|  | GGGAAGGGGCAGGAGAGaggtttgtctggtcaaccaccgcggtctcagtggtgt------------------------------ACGAGGAAGGGGGACAGGGAAGCCTGAGCGCCTCTCCTGGGCTTGCCAAGGAC | 0.09 % |
| | GGGAAGGGGCAGGAGAGaggtttgtctggtcaacc**C**ccgcggtctcagtggtgtacggtacaaacctCTGGGCTTGCCAAGGAC | 0.07 % |
| | GGGAAGGGGCAGGAGAGaggtttgtctggtcaaccaccgcggtctcagtggtgtacggtacaa**C**cctCTGGGCTTGCCAAGGAC | 0.07 % |

#### *AAVS1* / SpCas9(WT)-RT / Whole overlap (*attP* insertion)

Insertion
Subtitution
Deletion : -----

| | | |
|---|---|---|
| Precise insertion | GGGAAGGGGCAGGAGAGaggtttgtctggtcaaccaccgcggtctcagtggtgtacggtacaaacctCTGGGCTTGCCAAGGAC<br>*attP* (50bp) | 9.55 % |
| Pure indel | GGGAAGGGGCAGGAGAGCCAGGGGCATGAGATGGTGGACGAGGAAGGGGGACAGGGAAGCCTGAGCGCCTCTC**CT**CTGGGCTTGCCAAGGAC | 2.45 % |
|  | GGGAAGGGGCAGGAGAG--------------------------------------------------------CTGGGCTTGCCAAGGAC | 1.23 % |
| | GGGAAGGGGCAGGAGAGCCAGGGGCATGAGATGGTGGACGAGGAAGGGGGACAGGGAAGCCTGAGCGCCTCTC**T**CTGGGCTTGCCAAGGAC | 0.75 % |
| | GGGAAGGGGCAGGAGAGCCAGGGGCATGAGATGGTGGACGAGGAAGGGGGACAGGGAAGCCTGAGCGCCTCTC**CTT**CTGGGCTTGCCAAGGAC | 0.41 % |
| | GGGAAGGGGCAGGAGAG**GAGG**CCAGGGGCATGAGATGGTGGACGAGGAAGGGGGACAGGGAAGCCTGAGCGCCTCTCCTGGGCTTGCCAAGGAC | 0.29 % |
| Partial insertion | GGGAAGGGGCAGGAGAGaggtttgtctggtcaaccaccgcggtctcagt------------------CTGGGCTTGCCAAGGAC | 0.76 % |
|  | GGGAAGGGGCAGGAGAGaggtttgtctggtcaacca--------------------------------CTGGGCTTGCCAAGGAC | 0.67 % |
|  | GGGAAGGGGCAGGAGAGaggtttgtctggtca------------------------------------CTGGGCTTGCCAAGGAC | 0.59 % |
|  | GGGAAGGGGCAGGAGAGaggtttgtctggt--------------------------------------CTGGGCTTGCCAAGGAC | 0.48 % |
|  | GGGAAGGGGCAGGAGAGaggtttgtctggtcaaccaccgcggtctcagtggtgtacg-----------------------------------------------gtacggtacaaacctCTGGGCTTGCCAAGGAC<br>Sense insertion / Anti-Sense insertion | 0.43 % |

#### *AAVS1* / La-SpCas9(WT)-RT / Whole overlap (*attP* insertion)

Insertion
Subtitution
Deletion : -----

| | | |
|---|---|---|
| Precise insertion | GGGAAGGGGCAGGAGAGaggtttgtctggtcaaccaccgcggtctcagtggtgtacggtacaaacctCTGGGCTTGCCAAGGAC<br>*attP* (50bp) | 5.77 % |
| Pure indel | GGGAAGGGGCAGGAGAGCCAGGGGCATGAGATGGTGGACGAGGAAGGGGGACAGGGAAGCCTGAGCGCCTCTC**CT**CTGGGCTTGCCAAGGAC | 2.38 % |
|  | GGGAAGGGGCAGGAGAG---------------------------------------------------------**CT**CTGGGCTTGCCAAGGAC | 0.93 % |
|  | GGGAAGGGGCAGGAGAG---------------------------------------------------------CTGGGCTTGCCAAGGAC | 0.45 % |
|  | GGGAAGGGGCAGGAGAG--------------------------------------------------------GCC**A**C--CTGGGCTTGCCAAGGAC | 0.29 % |
|  | GGGAAGGGGCAGGAGAG---------------------------------------------------------**C**CTGGGCTTGCCAAGGAC | 0.28 % |
| Partial insertion | GGGAAGGGGCAGGAGAGaggtttg------------------------------------------CTGGGCTTGCCAAGGAC | 0.56 % |
|  | GGGAAGGGGCAGGAGAGaggtttgtctg**T**--------------------------------------CTGGGCTTGCCAAGGAC | 0.33 % |
|  | GGGAAGGGGCAGGAGAGaggtttgtctg---------------------------------------CTGGGCTTGCCAAGGAC | 0.29 % |
|  | GGGAAGGGGCAGGAGAGaggtttgtctggtcaaccaccgcggtctcagtggtgtacggtacaa----CTGGGCTTGCCAAGGAC | 0.29 % |
| | GGGAAGGGGCAGGAGAGaggtttgtctggtcaaccaccgcggtctcagtggtgtacggtacaaacctCCAGGGGCATGAGATGGTGGACGAGGAAGGGGGACAGGGAAGCCTGAGCGCCTCTCCTGGGCTTGCCAAGGAC | 0.28 % |

#### *AAVS1* / SpCas9(H840A)-RT / Homology arm (*attP* insertion)

Insertion
Subtitution
Deletion : -----

| | | |
|---|---|---|
| Precise insertion | GGGAAGGGGCAGGAGAGaggtttgtctggtcaaccaccgcggtctcagtggtgtacggtacaaacctCTGGGCTTGCCAAGGAC<br>*attP* (50bp) | 1.93 % |
| Pure indel | GGGAAGGGGCAGGAGAG--------------------------------------------------------CTGGGCTTGCCAAGGAC | 0.04 % |
| | GGGAAGGGGCAGGAGA-CCAGGGGCATGAGATGGTGGACGAGGAAGGGGGACAGGGAAGCCTGAGCGCCTCTCCTGGGCTTGCCAAGGAC | 0.02 % |
| Partial insertion | GGGAAGGGGCAGGAGAGaggtttgtctggtcaaccaccgcggtctcag------------------------------TGGTGGACGAGGAAGGGGGACAGGGAAGCCTGAGCGCCTCTCCTGGGCTTGCCAAGGAC | 0.20 % |
|  | GGGAAGGGGCAGGAGAGaggtttgtctggtcaaccaccgcggtctcagtggtgtacggtacaaacct**CTGGGCTTGCCAAGG**--------------------------------------GCCTGAGCGCCTCTCCTGGGCTTGCCAAGGAC | 0.06 % |
|  | GGGAAGGGGCAGGAGAGaggtttgtctggtcaaccaccgcgg**C**ctcagtggtg**C**acggtacaaacct-----------------------------------GG**C**A-CC-GAGCGCCTGAGCGCCTCTCCTGGGCTTGCCAAGGAC | 0.06 % |
| | GGGAAGGGGCAGGAGAGaggtttgtc**C**ggtcaaccaccgcggtctcagtggtgtacggtacaaacctCTGGGCTTGCCAAGGAC | 0.04 % |
| | GGGAAGGGGCAGGAGAGaggtttgtctggtcaaccaccgcggtctcagtggtg**C**acggtacaaacctCTGGGCTTGCCAAGGAC | 0.04 % |

#### *AAVS1* / La-SpCas9(H840A)-RT / Homology arm (*attP* insertion)

Insertion
Subtitution
Deletion : -----

| | Sequence | Frequency |
|---|---|---|
| Precise insertion | GGGAAGGGGCAGGAGAGaggtttgtctggtcaaccaccgcggtctcagtggtgtacggtacaaacctCTGGGCTTGCCAAGGAC<br>*attP* (50bp) | 0.76 % |
| Pure indel | GGGAAGGGGCAGGAGAG-------------------------------------------------------------CTGGGCTTGCCAAGGAC | 0.26 % |
|  | GGGAAGGGGCAG--------------GAGATGGTGGACGAGGAAGGGGGACAGGGAAGCCTGAGCGCCTCTCCTGGGCTTGCCAAGGAC | 0.22 % |
|  | GGGAAGGGGCAGGA--------------------------------------GGAAGCCTGAGCGCCTCTCCTGGGCTTGCCAAGGAC | 0.11 % |
|  | GGGA--------------AGGGGCATGAGATGGTGGACGAGGAAGGGGGACAGGGAAGCCTGAGCGCCTCTCCTGGGCTTGCCAAGGAC | 0.03 % |
| | GGGAAGGGGCAGGAGAG-CAGGGGCATGAGATGGTGGACGAGGAAGGGGGACAGGGAAGCCTGAGCGCCTCTCCTGGGCTTGCCAAGGAC | 0.02 % |
| Partial insertion | GGGAAGGGGCAGGAGAGaggtttgtctggtcaaccacc---------------------------------------------------------------GCCTGAGCGCCTCTCCTGGGCTTGCCAAGGAC | 0.12 % |
| | GGGAAGGGGCAGGAGAGaggtttgtcCggtcaaccaccgcggtctcagtggtgCacggCacaaacctCTGGGCTTGCCAAGGAC | 0.10 % |
| | GGGAAGGGGCAGGAGAGaggtttgtcCggtcaaccaccgcggtctcagtggtgtacggtacaaacctCTGGGCTTGCCAAGGAC | 0.08 % |
| | GGGAAGGGGCAGGAGAGaggtttgtctggtcaaccaccgcggtctcagtggtgtacggtGcaaacctCTGGGCTTGCCAAGGAC | 0.07 % |
| | GGGAAGGGGCAGGAGAGaggCttgtctggtcaaccaccgcggtcCcagtggCgtacggCacaaacctCTGGGCTTGCCAAGGAC | 0.07 % |

#### *AAVS1* / SpCas9(WT)-RT / Homology arm (*attP* insertion)

Insertion
Subtitution
Deletion : -----

| | Sequence | Frequency |
|---|---|---|
| Precise insertion | GGGAAGGGGCAGGAGAGaggtttgtctggtcaaccaccgcggtctcagtggtgtacggtacaaacctCTGGGCTTGCCAAGGAC<br>*attP* (50bp) | 0.78 % |
| Pure indel | GGGAAGGGGCAGGAGAG-------------------------------------------------------------CTGGGCTTGCCAAGGAC | 19.14 % |
| | GGGAAGGGGCAGGAGAGCCAGGGGCATGAGATGGTGGACGAGGAAGGGGGACAGGGAAGCCTGAGCGCCTCTCCCTGGGCTTGCCAAGGAC | 2.15 % |
| | GGGAAGGGGCAGGAGAGGCCAGGGGCATGAGATGGTGGACGAGGAAGGGGGACAGGGAAGCCTGAGCGCCTCTCCTGGGCTTGCCAAGGAC | 1.10 % |
| | GGGAAGGGGCAGGAGAGCCAGGGGCATGAGATGGTGGACGAGGAAGGGGGACAGGGAAGCCTGAGCGCCTCTCCTCTGGGCTTGCCAAGGAC | 0.85 % |
|  | GGGAAGGGGCAGGAGAG-------------------------------------------------------------ACTGGGCTTGCCAAGGAC | 0.68 % |
| Partial insertion | GGGAAGGGGCAGGAGAGaggttt------------------------------------------CCAGGGGCATGAGATGGTGGACGAGGAAGGGGGACAGGGAAGCCTGAGCGCCTCTCCTGGGCTTGCCAAGGAC | 0.16 % |
|  | GGGAAGGGGCAGGAGAGaggtttgtctggtcaaccaccgc-------------------------CCAGGGGCATGAGATGGTGGACGAGGAAGGGGGACAGGAAGCCTGAGCGCCTCTCCTGGGCTTGCCAAGGAC | 0.15 % |
| | GGGAAGGGGCAGGAGAGaggtttgtctggtcaaccaccgcggtctcagtggtgtacggtacaaacctCTGGGCTTGCTGGGCTTGCCAAGGAC | 0.08 % |
| | GGGAAGGGGCAGGAGAGaggtttgtctggtcaaccaccgcggtctcagtggtgtacggtacaaacctCTGCTGGGCTTGCCAAGGAC | 0.03 % |
| | GGGAAGGGGCAGGAGAaggtttgtctggtcaaccaccgcggtctcagtggtgtacggtacaaacctCTGGGCTTGCCAGGGGCATGAGATGGTGGACGAGGAAGGGGGACAGGGAAGCCTGAGCGCCTCTCCTGGGCTTGCCAAGGAC | 0.03 % |

#### *AAVS1* / La-SpCas9(WT)-RT / Homology arm (*attP* insertion)

Insertion
Subtitution
Deletion : -----

| | Sequence | Frequency |
|---|---|---|
| Precise insertion | GGGAAGGGGCAGGAGAGaggtttgtctggtcaaccaccgcggtctcagtggtgtacggtacaaacctCTGGGCTTGCCAAGGAC<br>*attP* (50bp) | 0.61 % |
| Pure indel | GGGAAGGGGCAGGAGAG-------------------------------------------------------------CTGGGCTTGCCAAGGAC | 12.89 % |
| | GGGAAGGGGCAGGAGAGCCAGGGGCATGAGATGGTGGACGAGGAAGGGGGACAGGGAAGCCTGAGCGCCTCTCCCTGGGCTTGCCAAGGAC | 2.05 % |
| | GGGAAGGGGCAGGAGAGCCAGGGGCATGAGATGGTGGACGAGGAAGGGGGACAGGGAAGCCTGAGCGCCTCTCCTCTGGGCTTGCCAAGGAC | 0.74 % |
| | GGGAAGGGGCAGGAGAGGCCAGGGGCATGAGATGGTGGACGAGGAAGGGGGACAGGGAAGCCTGAGCGCCTCTCCTGGGCTTGCCAAGGAC | 0.68 % |
|  | GGGAAGGGGCAGGAGA-------------------------------------------------------------TCTGGGCTTGCCAAGGAC | 0.61 % |
| Partial insertion | GGGAAGGGGCAGGAGAGaggtttgtctggtcaaccaccgcggtctcagtggtgtacggtaca-----CTGGGCTTGCCAAGGAC | 0.13 % |
|  | GGGAAGGGGCAGGAGAGaggtttgtctggtcaaccaccgc--------------------------CCAGGGGCATGAGATGGTGGACGAGGAAGGGGGACAGGGAAGCCTGAGCGCCTCTCCTGGGCTTGCCAAGGAC | 0.05 % |
| | GGGAAGGGGCAGGAGAaggtttgtctggtcaaccaccgcggtctcagtggtgtacggtacaaacctCTGGGCTTGCCAAGGGCCAGGGGCATGAGATGGTGGACGAGGAAGGGGGACAGGGAAGCCTGAGCGCCTCTCCTGGGCTTGCCAAGGAC | 0.04 % |
| | GGGAAGGGGCAGGAGAGaggtttgtctggtcaaccaccgcggtctcagtggtgtacggtacaaacctCTCTGGGCTTGCCAAGGAC | 0.04 % |
|  | GGGAAGGGGCAGGAGAGaggtttgtctggtcaaccaccgcggtctc--------------------CTGGGCTTGCCAAGGAC | 0.04 % |

**Insertion**
**Subtitution**
**Deletion : -----**

#### *AAVS1* / SpCas9(H840A)-RT / Partial overlap (*attP* insertion)

**Precise insertion** GGGAAGGGGCAGGAGAGaggtttgtctggtcaaccaccgcggtctcagtggtgtacggtacaaacctCTGGGCTTGCCAAGGAC 6.39 %
*attP* (50bp)

**Pure indel**
GGGA---------------AGGGGCATGAGATGGTGGACGAGGAAGGGGGACAGGGAAGCCTGAGCGCCTCTCCTGGGCTTGCCAAGGAC 0.08 %
GGGAAGGGGCAGGAGAG-CAGGGGCATGAGATGGTGGACGAGGAAGGGGGACAGGGAAGCCTGAGCGCCTCTCCTGGGCTTGCCAAGGAC 0.02 %

**Partial insertion**
GGGAAGGGGCAGGAGAG------------AtcaaccaccgcggtctcagtggtgtacggtacaaacctCTGGGCTTGCCAAGGAC 0.10 %
GGGAAGGGGCAGGAGAGaggtttgtctggt-------cgcggtctcagtggtgtacggtacaaacctCTGGGCTTGCCAAGGAC 0.09 %
GGGAAGGGGCAGGAGAGaggtttgtctggtcaaccaccgc------------------------------------gtcaaccaccgcggtctcagtggtgtacggtacaaacctCTGGGCTTGCCAAGGAC 0.06 %
Sense insertion Anti-Sense insertion
GGGAAGGGGCAGGAGAGCCAGGGGCATGAGATGGTGGACGAGGAAGGGGGACAGGGAAGC-------------------gtcaaccaccgcggtctcagtggtgtacggtacaaacctCTGGGCTTGCCAAGGAC 0.05 %
GGGAAGGGGCAGGAGAGaggtttgtctggtcaaccaccgcggtctcagtggtgtacggtacaaacctCTGGGCT---------------------------------------GAAGCCTGAGCGCCTCTCCTGGGCTTGCCAAGGAC 0.04 %

**Insertion**
**Subtitution**
**Deletion : -----**

#### *AAVS1* / La-SpCas9(H840A)-RT / Partial overlap (*attP* insertion)

**Precise insertion** GGGAAGGGGCAGGAGAGaggtttgtctggtcaaccaccgcggtctcagtggtgtacggtacaaacctCTGGGCTTGCCAAGGAC 11.79 %
*attP* (50bp)

**Pure indel** None

**Partial insertion**
GGGAAGGGGCAGGAGAG--------ctgCgtcaaccaccgcggtctcagtggtgtacggtacaaacctCTGGGCTTGCCAAGGAC 0.18 %
GGGAAGGGGCAGGAGAGaggtttgtctggtcaaccaccgc---------ggtgtacggtacaaacctCTGGGCTTGCCAAGGAC 0.11 %
GGGAAGGGGCAGGAGAGaggtttgtctCgtcaaccaccgcggtctcagtggtgtacggtacaaacctCTGGGCTTGCCAAGGAC 0.08 %
GGGAAGGGGCAGGAGAGaggtttgtctggtcaaccaccgcggtctcagCggt-----------------------------------gcggtctcagtggtgtacggtacaaacctCTGGGCTTGCCAAGGAC 0.07 %
Sense insertion Anti-Sense insertion
GGGAAGGGGCAGGAGAGaggtttgtctggtcaaccaccgcggtctcagtggt----------------------------------------------gtggtgtacggtacaaacctCTGGGCTTGCCAAGGAC 0.07 %
Sense insertion Anti-Sense insertion

**Insertion**
**Subtitution**
**Deletion : -----**

#### *AAVS1* / SpCas9(WT)-RT / Partial overlap (*attP* insertion)

**Precise insertion** GGGAAGGGGCAGGAGAGaggtttgtctggtcaaccaccgcggtctcagtggtgtacggtacaaacctCTGGGCTTGCCAAGGAC 3.74 %
*attP* (50bp)

**Pure indel**
GGGAAGGGGCAGGAGAG------------------------------------------------------------------CTGGGCTTGCCAAGGAC 2.90 %
GGGAAGGGGCAGGAGAGGCCAGGGGCATGAGATGGTGGACGAGGAAGGGGGACAGGGAAGCCTGAGCGCCTCTCCTGGGCTTGCCAAGGAC 1.71 %
GGGAAGGGGCAGGAGAGCCAGGGGCATGAGATGGTGGACGAGGAAGGGGGACAGGGAAGCCTGAGCGCCTCTCCCTGGGCTTGCCAAGGAC 0.35 %
GGGAAGGGGCAG------------GAGATGGTGGACGAGGAAGGGGGACAGGGAAGCCTGAGCGCCTCTCCTGGGCTTGCCAAGGAC 0.27 %
GGGAAGGGGCAGGAGAGCCAGGGGCATGAGATGGTGGACGAGGAAGGGGGACAGGGAAGCCTGAGCGCCTCTCCTCTGGGCTTGCCAAGGAC 0.21 %

**Partial insertion**
GGGAAGGGGCAGGAGAG----------CgtcaaccaccgcggtctcagtggtgtacggtacaaacctCTGGGCTTGCCAAGGAC 0.35 %
GGGAAGGGGCAGGAGAGaggtttgtctggtcaaccaccgcggtctcagtggtgtacg----------CTGGGCTTGCCAAGGAC 0.33 %
GGGAAGGGGCAGGAGAGaggtttgtctggtcaaccaccgcggtctcagtg-----------------CTGGGCTTGCCAAGGAC 0.30 %
GGGAAGGGGCAGGAGAGaggtttgtctggtcaaccaccgcg-------------------------CTGGGCTTGCCAAGGAC 0.30 %
GGGAAGGGGCAGGAGAG--------c--gtcaaccaccgcggtctcagtggtgtacggtacaaacctCTGGGCTTGCCAAGGAC 0.20 %

##### *AAVS1* / La-SpCas9(WT)-RT / Partial overlap (*attP* insertion)

Insertion
Subtitution
Deletion : -----

**Precise insertion** GGGAAGGGGCAGGAGAGaggtttgtctggtcaaccaccgcggtctcagtggtgtacggtacaaacctCTGGGCTTGCCAAGGAC — 5.12 %
*attP* (50bp)

**Pure indel**
GGGAAGGGGCAGGAGAG--------------------------------------------------------CTGGGCTTGCCAAGGAC — 1.51 %
GGGAAGGGGCAGGAGAGAG-------------------------------------------------------CTGGGCTTGCCAAGGAC — 0.74 %
GGGAAGGGGCAGGAGAGCCAGGGGCATGAGATGGTGGACGAGGAAGGGGGACAGGGAAGCCTGAGCGCCTCTCTCTGGGCTTGCCAAGGAC — 0.23 %
GGGAAGGGGCAGGAGAGCCAGGGGCATGAGATGGTGGACGAGGAAGGGGGACAGGGAAGCCTGAGCGCCTCTCCTCTGGGCTTGCCAAGGAC — 0.13 %
GGGAAGGGGCAGGAGAGAGGTCCAGGGGCATGAGATGGTGGACGAGGAAGGGGGACAGGGAAGCCTGAGCGCCTCTCCTGGGCTTGCCAAGGAC — 0.03 %

**Partial insertion**
GGGAAGGGGCAGGAGAGaggtttgtctgg--------------------------------------CTGGGCTTGCCAAGGAC — 0.27 %
GGGAAGGGGCAGGAGAGaggtttgtctggtcaa---------------------------------CCTGGGCTTGCCAAGGAC — 0.26 %
GGGAAGGGGCAGGAGAGaggtttgtctggtcaa---------------------------------CTGGGCTTGCCAAGGAC — 0.20 %
GGGAAGGGGCAGGAGAGaggtttgtctggtcaaccaccgc------------------------------CCAGGGGCATGAGATGGTGGACGAGGAAGGGGGACAGGGAAGCCTGAGCGCCTCTCCTGGGCTTGCCAAGGAC — 0.20 %
GGGAAGGGGCAGGAGAGaggtttgtctggtcaaccaccgcggtctcagtggtgtac-----------CCAGGGGCATGAGATGGTGGACGAGGAAGGGGGACAGGGAAGCCTGAGCGCCTCTCCTGGGCTTGCCAAGGAC — 0.17 %

#### *HEK3* Twin-prime editing pattern (*attB* insertion)

***HEK3***

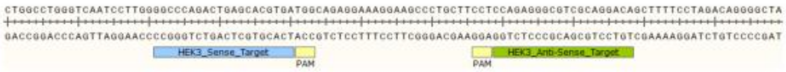

Insertion
Subtitution
Deletion : -----

**Normal control** GGCCCAGACTGAGCACGTGATGGCAGAGGAAAGGAAGCCCTGCTTCCTCCAGAGGGCGTCGCAGGACA

##### *HEK3* / SpCas9(H840A)-RT / Whole overlap (*attB* insertion)

Insertion
Subtitution
Deletion : -----

**Precise insertion** GGCCCAGACTGAGCACGggcttgtcgacgacggcggtctccgtcgtcaggatcatGAGGGCGTCGCAGGACA — 24.24 %
*attB* (38bp)

**Pure indel** None

**Partial insertion**
GGCCCAGACTGAGCACGggcttgtcgacgacggcggtctccgtcgtcaggatcatGCCAGAGGGCGTCGCAGGACA — 0.43 %
GGCCCAGACTGAGCACGTgcttgtcgacgacggcggtctccgtcgtcaggatcatGAGGGCGTCGCAGGACA — 0.43 %
GGCCCAGACTGAGCACG-----------------------------caggatcatGAGGGCGTCGCAGGACA — 0.31 %
GGCCCAGACTGAGCACGTGATGGCAAAGGATggcttgtcgacgacggcggtctccgtcgtcaggatcatGAGGGCGTCGCAGGACA — 0.31 %
GGCCCAGACTGAGCACGTGATGGCAAAGGAAAGGggcttgtcgacgacggggtctccgtcgtcaggatcatGAGGGCGTCGCAGGACA — 0.25 %

##### *HEK3* / La-SpCas9(H840A)-RT / Whole overlap (*attB* insertion)

Insertion
Subtitution
Deletion : -----

**Precise insertion** GGCCCAGACTGAGCACGggcttgtcgacgacggcggtctccgtcgtcaggatcatGAGGGCGTCGCAGGACA — 49.59 %
*attB* (38bp)

**Pure indel** None

**Partial insertion**
GGCCCAGACTGAGCACGggctt---------------------------------GAGGGCGTCGCAGGACA — 0.43 %
GGCCCAGACTGAGCACGTgcttgtcgacgacggcggtctccgtcgtcaggatcatGAGGGCGTCGCAGGACA — 0.39 %
GGCCCAGACTGAGCACGggcttgtcgacgacggcggtctccgtcgtcaggatcat----GCGTCGCAGGACA — 0.33 %
GGCCCAGACTGAGCACGggcttgtcgacgacggcggtctccgtcgtcaggatcatGCAGGGCGTCGCAGGACA — 0.33 %
GGCCCAGACTGAGCACGggcttgtcgacgacggcggtctccgtcgtcagga-----------GGGCGTCGCAGGACA — 0.24 %

Insertion
Subtitution
Deletion : -----

#### *HEK3* / SpCas9(WT)-RT / Whole overlap (*attB* insertion)

| | Sequence | % |
|---|---|---|
| Precise insertion | GGCCCAGACTGAGCACGggcttgtcgacgacggcggtctccgtcgtcaggatcatGAGGGCGTCGCAGGACA<br>*attB* (38bp) | 18.21 % |
| Pure indel | GGCCCAGACTGAGCACG--------------------------------------GAGGGCGTCGCAGGACA | 12.29 % |
|  | GGCCCAGACTGAGCAC---------------------------------------GAGGGCGTCGCAGGACA | 0.80 % |
|  | GGCCCAGACTGAGCACG-GAT-----------------------------------catGAGGGCGTCGCAGGACA | 0.73 % |
| | GGCCCAGACTGAGCACGGTGATGGCAGAGGAAAGGAAGCCCTGCTTCCTCCAGAGGGCGTCGCAGGACA | 0.50 % |
| | GGCCCAGACTGAGCACGAGTGATGGCAGAGGAAAGGAAGCCCTGCTTCCTCCAGAGGGCGTCGCAGGACA | 0.28 % |
| Partial insertion | GGCCCAGACTGAGCAC-ggcttgtcgacgacggcggtctccgtcgtcaggatcatGAGGGCGTCGCAGGACA | 1.68 % |
| | GGCCCAGACTGAGCACGTGATGGCAGAGGAAAGGAAGCCCTGCTTCCTCCAggcttgtcgacgacggcggtctccgtcgtcaggatcatGAGG<br>GCGTCGCAGGACA | 0.83 % |
|  | GGCCCAGACTGAGCACG--------gacgacggcggtctccgtcgtcaggatcatGAGGGCGTCGCAGGACA | 0.66 % |
|  | GGCCCAGACTGAGCACGggcttgtcg-----------------------------GAGGGCGTCGCAGGACA | 0.58 % |
|  | GGCCCAGACTGAGCACGggcttgtcgacgacggcggtctccg------------tGAGGGCGTCGCAGGACA | 0.49 % |

Insertion
Subtitution
Deletion : -----

#### *HEK3* / La-SpCas9(WT)-RT / Whole overlap (*attB* insertion)

| | Sequence | % |
|---|---|---|
| Precise insertion | GGCCCAGACTGAGCACGggcttgtcgacgacggcggtctccgtcgtcaggatcatGAGGGCGTCGCAGGACA<br>*attB* (38bp) | 25.00 % |
| Pure indel | GGCCCAGACTGAGCACG--------------------------------------GAGGGCGTCGCAGGACA | 6.26 % |
|  | GGCCCAGACTGAGCAC---------------------------------------GAGGGCGTCGCAGGACA | 1.58 % |
|  | GGCCCAGACTGAGCA---------AGGAAAGGAAGCCCTGCTTCCTCCAGAGGGCGTCGCAGGACA | 0.36 % |
| | GGCCCAGACTGAGCACGGTGATGGCAGAGGAAAGGAAGCCCTGCTTCCTCCAGAGGGCGTCGCAGGACA | 0.21 % |
| | GGCCCAGACTGAGCACGCTGATGGCAGAGGAAAGGAAGCCCTGCTTCCTCCAGAGGGCGTCGCAGGACA | 0.21 % |
| Partial insertion | GGCCCAGACTGAGCACGggcttgtcgacgacggcggtctccgtcgtcaggatc--GAGGGCGTCGCAGGACA | 1.24 % |
| | GGCCCAGACTGAGCACGggcttgtcgacgacggcggtctccgtcgtcaggatcatTGATGGCAGAGGAAAGGAAGCCCTGCTTCCTCCAGAGG<br>GCGTCGCAGGACA | 1.16 % |
|  | GGCCCAGACTGAGCACGggcttgtcgacgacggcggtctc---------------GAGGGCGTCGCAGGACA | 0.99 % |
|  | GGCCCAGACTGAGCACGggcttgtcgacgacggcgg-------------------GAGGGCGTCGCAGGACA | 0.79 % |
|  | GGCCCAGACTGAGCACGggcttgtcgacgacggcggtctccg-T-----------GAGGGCGTCGCAGGACA | 0.79 % |

Insertion
Subtitution
Deletion : -----

#### *HEK3* / SpCas9(H840A)-RT / Homology arm (*attB* insertion)

| | Sequence | % |
|---|---|---|
| Precise insertion | GGCCCAGACTGAGCACGggcttgtcgacgacggcggtctccgtcgtcaggatcatGAGGGCGTCGCAGGACA<br>*attB* (38bp) | 10.24 % |
| Pure indel | None | |
| Partial insertion | GGCCCAGACTGAGCACGCTGAGCACGggcttgtcgacgacggcggtctccgtcgtcaggatcatGAGGGCGTCGCAGGACA | 0.47 % |
| | GGCCCAGACTGAGCACGCggcttgtcgacgaGCACGggcttgtcgacgacggcggtctccgtcgtcaggatcatGAGGGCGTCGCAGGACA<br>Sense insertion / Anti-Sense insertion | 0.29 % |
|  | GGCCCAGACTGAGCACGCgcttgtcgacgacggcggtctccgtcgtcaggatcat-----------GAGGGC-GTCTCCGTCGTCAGGATCATGAGGGC<br>GTCGCAGGACA<br>Sense insertion / Anti-Sense insertion | 0.20 % |
| | GGCCCAGACTGAGCACGCggcttgtcgacgacggcggtctccgtcgtcaggatcatGAAGGCGTCGCAGGACA | 0.03 % |
| | GGCCCAGACTGAGCACGCggcttgtcgacgacggcgCtctccgtcgtcaggatcatGAGGGCGTCGCAGGACA | 0.02 % |

Insertion
Subtitution
Deletion : -----

#### *HEK3* / La-SpCas9(H840A)-RT / Homology arm (*attB* insertion)

| | Sequence | % |
|---|---|---|
| Precise insertion | GGCCCAGACTGAGCACGggcttgtcgacgacggcggtctccgtcgtcaggatcatGAGGGCGTCGCAGGACA<br>*attB* (38bp) | 7.31 % |
| Pure indel | None | |
| Partial insertion | GGCCCAGACTGAGCACGCggcttgtcgacgacggcggtctccgtcgtcaggatcatGAGGGCGTGAGGGCGTCGCAGGACA | 0.32 % |
| | GGCCCAGACTGAGCACGCggcttgtcgacgacggcggtctccgtcgtcaggatcatGAGGGCGTCGCAGGGAGGGCGTCGCAGGACA | 0.31 % |
| | GGCCCAGACTGAGCACGCggcttgtcgacgacggcggtctccgtcgtcaggatcCtGAGGGCGTCGCAGGACA | 0.05 % |
| | GGCCCAGACTGAGCACGCggcttgtcgacgacggTggtctccgtcgtcaggatcatGAGGGCGTCGCAGGACA | 0.01 % |
| | GGCCCAGACTGAGCACGCggcttgtcgacgacggcggCctccgtcgtcaggatcatGAGGGCGTGAGGGCGTCGCAGGACA | 0.01 % |

#### *HEK3* / SpCas9(WT)-RT / Homology arm (*attB* insertion)

Insertion
Subtitution
Deletion : -----

| | | |
|---|---|---|
| Precise insertion | GGCCCAGACTGAGCACGggcttgtcgacgacggcggtctccgtcgtcaggatcatGAGGGCGTCGCAGGACA<br>*attB* (38bp) | 3.52 % |
| Pure indel | GGCCCAGACTGAGCACG--------------------------------------GAGGGCGTCGCAGGACA | 34.10 % |
|  | GGCCCAGACTGAGCAC---------------------------------------GAGGGCGTCGCAGGACA | 0.57 % |
|  | GGCCCAGACTGA----------GCAGAGGAAAGGAAGCCCTGCTTCCTCCAGAGGGCGTCGCAGGACA | 0.55 % |
|  | GGCCCAGAC-----------TGGCAGAGGAAAGGAAGCCCTGCTTCCTCCAGAGGGCGTCGCAGGACA | 0.38 % |
|  | GGCCCAGACTGAGC-----ATGGCAGAGGAAAGGAAGCCCTGCTTCCTCCAGAGGGCGTCGCAGGACA | 0.34 % |
| Partial insertion | GGCCCAGACTGAGCACGGCACGggcttgtcgacgacggcggtctccgtcgtcaggatcatGAGGGCGTCGCAGGACA | 0.39 % |
|  | GGCCCAGACTGAGCACG-----------gacggcggtctccgtcgtcaggatcatGAGGGCGTCGCAGGACA | 0.30 % |
|  | GGCCCAGACTGAGCACG--------------------ctccgtcgtcaggatcatGAGGGCGTCGCAGGACA | 0.26 % |
|  | GGCCCAGACTGAGCACG---------acgacggcggtctccgtcgtcaggatcatGAGGGCGTCGCAGGACA | 0.19 % |
| | GGCCCAGACTGAGCACGCCCAGACTGAGCACGggcttgtcgacgacggcggtctccgtcgtcaggatcatGAGGGCGTCGCAGGACA | 0.17 % |

#### *HEK3* / La-SpCas9(WT)-RT / Homology arm (*attB* insertion)

Insertion
Subtitution
Deletion : -----

| | | |
|---|---|---|
| Precise insertion | GGCCCAGACTGAGCACGggcttgtcgacgacggcggtctccgtcgtcaggatcatGAGGGCGTCGCAGGACA<br>*attB* (38bp) | 3.57 % |
| Pure indel | GGCCCAGACTGAGCACG--------------------------------------GAGGGCGTCGCAGGACA | 36.12 % |
|  | AGCCCTGGCCT--------------------------------------------------------------------GGACA | 4.02 % |
| | GGCCCAGACTGAGCAGCGTGATGGCAGAGGAAAGGAAGCCCTGCTTCCTCCAGAGGGCGTCGCAGGACA | 0.45 % |
| | GGCCCAGACTGAGCAC-TGATGGCAGAGGAAAGGAAGCCCTGCTTCCTCCAGAGGGCGTCGCAGGACA | 0.31 % |
| | GGCCCAGACTGAGCACGTGATGGCAGAGGAAAGGAAGCCCTGCTTCCTCCAGGGAGGGCGTCGCAGGACA | 0.29 % |
| Partial insertion | GGCCCAGACTGAGCACGggcttgtcgacgacggcggtctccgtcgtcaggatcatGAGGGCGTCGCAGGAGCGAGGGCGTCGCAGGACA | 1.36 % |
| | GGCCCAGACTGAGCACGGCACggcttgtcgacgacggcggtctccgtcgtcaggatcatGAGGGCGTCGCAGGACA | 0.42 % |
| | GGCCCAGACTGAGACTGAGCACGggcttgtcgacgacggcggtctccgtcgtcaggatcatGAGGGCGTCGCAGGACA | 0.34 % |
| | GGCCCAGACTGAGCACGggcttgtTGATGGCAGAGGAAAGGAAGCCCTGCTTCCTCCAGAGGGCGTCGCAGGACA | 0.24 % |
|  | GGCCCAGACTGAGCACGggcttgtcgacgacggcggtctccgtcg----------TGATGGCAGAGGAAAGGAAGCCCTGCTTCCTCCAGAGGGC<br>GTCGCAGGACA | 0.22 % |

#### *HEK3* / SpCas9(H840A)-RT / Partial insertion (*attB* insertion)

Insertion
Subtitution
Deletion : -----

| | | |
|---|---|---|
| Precise insertion | GGCCCAGACTGAGCACGggcttgtcgacgacggcggtctccgtcgtcaggatcatGAGGGCGTCGCAGGACA<br>*attB* (38bp) | 15.95 % |
| Pure indel | None | |
| Partial insertion | GGCCCAGACTGAGCACGggcttgtcgacgacggcggtctcc--cCtc-----ca-GAGGGCGTCGCAGGACA | 0.21 % |
| | GGCCCAGACTGAGCACGggcttgtCGAcgacgacggcggtctccgtcgtcaggatcatGAGGGCGTCGCAGGACA | 0.19 % |
|  | GGCCCAGACTGAGCACGggcttgtcgacgacggcg---------gtcgtcaggatcatGAGGGCGTCGCAGGACA | 0.19 % |
| | GGCCCAGACTGAGCACGggcttgtcgacgacggcggtctcCGTCGTGcgtcgtcaggatcatGAGGGCGTCGCAGGACA | 0.18 % |
|  | GGCCCAGACTGAGCACG--------------ggcggtctccgtcgtcaggatcatGAGGGCGTCGCAGGACA | 0.17 % |

#### *HEK3* / La-SpCas9(H840A)-RT / Partial insertion (*attB* insertion)

Insertion
Subtitution
Deletion : -----

| | | |
|---|---|---|
| Precise insertion | GGCCCAGACTGAGCACGggcttgtcgacgacggcggtctccgtcgtcaggatcatGAGGGCGTCGCAGGACA<br>*attB* (38bp) | 47.51 % |
| Pure indel | None | |
| Partial insertion | GGCCCAGACTGAGCACGggcttgtcgacgacggcggtctccgtcgt--g---ca-GAGGGCGTCGCAGGACA | 1.00 % |
|  | GGCCCAGACTGAGCACGggcttgtcgacgac-----------------------GAGGGCGTCGCAGGACA | 0.96 % |
|  | GGCCCAGACTGAGCACGggcttgtcgacgac------------------------------------GGACA | 0.81 % |
|  | GGCCCAGACTGAGCACGggcttgtcgacgacggcggtctc---------------------------cggcggtctccgtcgtcaggatcatGAGGGCGTCGCAGGACA<br>Sense insertion / Anti-Sense insertion | 0.69 % |
|  | GGCCCAGAC----------------------------------gtcgtcaggatcatGAGGGCGTCGCAGGACA | 0.51 % |

##### *HEK3* / SpCas9(WT)-RT / Partial insertion (*attB* insertion)

Insertion
Subtitution
Deletion : -----

**Precise insertion** GGCCCAGACTGAGCACGggcttgtcgacgacggcggtctccgtcgtcaggatcatGAGGGCGTCGCAGGACA — 13.78 %
*attB* (38bp)

**Pure indel**
GGCCCAGACTGAGCACG--------------------------------CATGAGGGCGTCGCAGGACA — 2.59 %
GGCCCAGAC--------TGATGGCAGAGGAAAGGAAGCCCTGCTTCCTCCAGAGGGCGTCGCAGGACA — 1.20 %
GGCCCAGACTGAGCACGTCAT-----------------------------------GAGGGCGTCGCAGGACA — 0.94 %
GGCCCAGACTGAGCACG---TGGCAGAGGAAAGGAAGCCCTGCTTCCTCCAGAGGGCGTCGCAGGACA — 0.75 %
GGCCCAGACTGAGCACGCGTGATGGCAGAGGAAAGGAAGCCCTGCTTCCTCCAGAGGGCGTCGCAGGACA — 0.31 %

**Partial insertion**
GGCCCAGACTGAGCACGggcttgtcgac-----------------------GAGGGCGTCGCAGGACA — 2.71 %
GGCCCAGACTGAGCACGggcttgtcgacgacggcgCt-----------------GAGGGCGTCGCAGGACA — 2.39 %
GGCCCAGACTGAGCACGggcttgtcgacgacggcggtctccgtcgt---------GAGGGCGTCGCAGGACA — 1.98 %
GGCCCAGACTGAGCACGggcttgtcgacgacg----------------------GAGGGCGTCGCAGGACA — 1.61 %
GGCCCAGACTGAGCACG--------------------tccgtcgtcaggatcatGAGGGCGTCGCAGGACA — 1.60 %

##### *HEK3* / La-SpCas9(WT)-RT / Partial insertion (*attB* insertion)

Insertion
Subtitution
Deletion : -----

**Precise insertion** GGCCCAGACTGAGCACGggcttgtcgacgacggcggtctccgtcgtcaggatcatGAGGGCGTCGCAGGACA — 14.82 %
*attB* (38bp)

**Pure indel**
GGCCCAGACTGAGCACG-----------------------------------GAGGGCGTCGCAGGACA — 2.49 %
GGCCCAGACTGAGCACGTGATGGCAGAGGAAAGGAAGCCCTGCTTCCTCCAAGAGGGCGTCGCAGGACA — 0.54 %
GGCCCAGACTGAGCACGTGATGGCAGAGGAAAGGAAGCCCTGCTTCCTCCA----GCGTCGCAGGACA — 0.32 %
GGCCCAGACTGAGCACGTGATGGCAGAGGAAAGGAAGCCCTGCTTCCTCCATGAGGGCGTCGCAGGACA — 0.26 %
GGCCCAGACTGAGCACGTGATGGCAGAGGAAAGGAAGCCCTGCTTCCTCCATCATGAGGGCGTCGCAGGACA — 0.18 %

**Partial insertion**
GGCCCAGACTGAGCA-----------cgacggcggtctccgtcgtcaggatcatGAGGGCGTCGCAGGACA — 4.13 %
GGCCCAGACTGAGCACGggcttgtcgacgacggcggtct----------------GAGGGCGTCGCAGGACA — 2.52 %
GGCCCAGACTGAGCACGggcttgtcgacgacggcggtctccgtcgt---------GAGGGCGTCGCAGGACA — 2.37 %
GGCCCAGACTGAGCACGggcttgtcgacgacggcggtctccgtcgt---------TGATGGCAGAGGAAAGGAAGCCCTGCTTCCTCCAGAGGGC
GTCGCAGGACA — 2.21 %
GGCCCAGACTGAGCACGggcttgtcgacgacggcggtct---------------GAGGGCGTCGCAGGACA — 1.78 %

### *FANCF* Twin-prime editing pattern (*attB* insertion)

***FANCF***

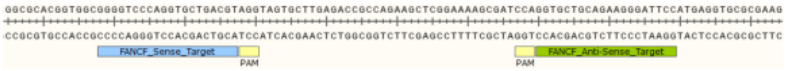


Insertion
Subtitution
Deletion : -----

**Normal control** GGGGTCCCAGGTGCTGACGTAGGTAGTGCTTGAGACCGCCAGAAGCTCGGAAAAGCGATCCAGGTGCTGCAGAAGGGATTCC

##### *FANCF* / SpCas9(H840A)-RT / Whole overlap (*attB* insertion)

Insertion
Subtitution
Deletion : -----

**Precise insertion** GGGGTCCCAGGTGCTGAatgatcctgacgacggagaccgccgtcgtcgacaagccGCTGCAGAAGGGATTCC — 23.31 %
*attB* (38bp)

**Pure indel**
GGGGTCCCAGGTGCTGACGT---------------------AGCTCGGAAAAGCGATCCAGGTGCTGCAGAAGGGATTCC — 0.56 %

**Partial insertion**
GGGGTCCCAGGTGCTGACGTAGGTAGTGCTTGAGACCGCC----------------------------------------------gccgtcgtcgacaagccGCTGCAG
AAGGGATTCC — 1.88 %
GGGGTCCCAGGTGCTGAatgatcctgacgacg----------------------------------GAGACCGCCAGAAGCTCGGAAAAGCGATCCAGGTGCT
GCAGAAGGGATTCC — 0.48 %
GGGGTCCCAGGTGCTGA------------------------------CcaagccGCTGCAGAAGGGATTCC — 0.44 %
GGGGTCCCAGGTGCTGA---------------gagaccgccgtcgtcgacaagccGCTGCAGAAGGGATTCC — 0.22 %
GGGGTCCCAGGTG----CatgatcctgacgacggagaccgccgtcgtcgacaagccGCTGCAGAAGGGATTCC — 0.22 %

#### *FANCF* / La-SpCas9(H840A)-RT / Whole overlap (*attB* insertion)

Insertion
Subtitution
Deletion : -----

| | | |
|---|---|---|
| Precise insertion | GGGGTCCCAGGTGCTGAatgatcctgacgacggagaccgccgtcgtcgacaagccGCTGCAGAAGGGATTCC<br>*attB* (38bp) | 36.23 % |
| Pure indel | None | 0.56 % |
| Partial insertion | GGGGTCCCAGGTGCTGACGTAGGTAGTGCTTGAGACCGCC---------------------------------------------------gtcgtcgacaagccGCTGCAGAAGGGATTCC | 1.25 % |
|  | GGGGTCCCAGGTGCTGAatgatcctgacgacggagaccgc---------------atgatcctgacgacggagaccgccgtcgtcgacaagccGCTGCAGAAGGGATTCC | 0.27 % |
|  | GGGGTCCCAGGTGCTGAatgatcctgacgacggagaccgccgtcgtc--------------ctgacgacggagaccgccgtcgtcgacaagccGCTGCAGAAGGGATTCC<br>Sense insertion / Anti-Sense insertion | 0.11 % |
|  | GGGGTCCCAGGTGCTGAatgatcctgacgacg-------------------------GAGACCGCCAGAAGCTCGGAAAAGCGATCCAGGTGCTGCAGAAGGGATTCC | 0.11 % |
| | GGGGTCCCAGGTGCTGAatgatcctgacgacggagaccgccgtcgtcgacaagGTGCTGCAGAAGGGATTCC | 0.08 % |

#### *FANCF* / SpCas9(WT)-RT / Whole overlap (*attB* insertion)

Insertion
Subtitution
Deletion : -----

| | | |
|---|---|---|
| Precise insertion | GGGGTCCCAGGTGCTGAatgatcctgacgacggagaccgccgtcgtcgacaagccGCTGCAGAAGGGATTCC<br>*attB* (38bp) | 16.16 % |
| Pure indel | GGGGTCCCAGGTGCTGA------------------------------------------------GCTGCAGAAGGGATTCC | 7.01 % |
|  | GGGGTCCCAGGTGCTGA------------------------------------------------GCCGCTGCAGAAGGGATTCC | 1.15 % |
|  | GGGGTC----------ACGTAGGTAGTGCTTGAGACCGCCAGAAGCTCGGAAAAGCGATCCAGGTGCTGCAGAAGGGATTCC | 0.63 % |
| | GGGGTCCCAGGTGCTGAATCGTAGGTAGTGCTTGAGACCGCCAGAAGCTCGGAAAAGCGATCCAGGTGCTGCAGAAGGGATTCC | 0.37 % |
|  | GGGGTCCC--------ACGTAGGTAGTGCTTGAGACCGCCAGAAGCTCGGAAAAGCGATCCAGGTGCTGCAGAAGGGATTCC | 0.35 % |
| Partial insertion | GGGGTCCCAGGTGCTGAatgatcctgacgacggagaccgccgtcgtcgacaagccCGTAGGTAGTGCTTGAGACCGCCAGAAGCTCGGAAAAGCGATCCAGGTGCTGCAGAAGGGATTCC | 0.88 % |
|  | GGGGTCCCAGGTGCTGACGTAGGTAGTGCTTGAGACCGCC-------------------------------------------------gtcgtcgacaagccGCTGCAGAAGGGATTCC | 0.77 % |
| | GGGGTCCCAGGTGCTG-atgatcctgacgacggagaccgccgtcgtcgacaagccGCTGCAGAAGGGATTCC | 0.61 % |
|  | GGGGTCCCAGGTGCTGAatgatcctgacg-------------------------CGTAGGTAGTGCTTGAGACCGCCAGAAGCTCGGAAAAGCGATCCAGGTGCTGCAGAAGGGATTCCGCTGCAGAAGGGATTCC | 0.54 % |
| | GGGGTCCCAGGTGCTGAatgatcctgacgacggagaccgccgtcgtcgacaagc--CTGCAGAAGGGATTCC | 0.46 % |

#### *FANCF* / La-SpCas9(WT)-RT / Whole overlap (*attB* insertion)

Insertion
Subtitution
Deletion : -----

| | | |
|---|---|---|
| Precise insertion | GGGGTCCCAGGTGCTGAatgatcctgacgacggagaccgccgtcgtcgacaagccGCTGCAGAAGGGATTCC<br>*attB* (38bp) | 26.09 % |
| Pure indel | GGGGTCCCAGGTGCTGA------------------------------------------------GCTGCAGAAGGGATTCC | 5.06 % |
|  | GGGGTCCCAGGTGCTGA------------------------------------------------AGCTGCAGAAGGGATTCC | 1.14 % |
| | GGGGTCCCAGGTGCTGACGTAGGTAGTGCTTGAGACCGCCAGAAGCTCGGAAAAGCGATCCAGGTGGCTGCAGAAGGGATTCC | 0.14 % |
| Partial insertion | GGGGTCCCAGGTGCTGACGTAGGTAGTGCTTGAGACCGCC-------------------------------------------------gtcgtcgacaagccGCTGCAGAAGGGATTCC | 0.91 % |
| | GGGGTCCCAGGTGCTGACGTAGGTAGTGCTTGAGACCGCCAGAAGCTCGGAAAAGCGATCCAGGTatgatcctgacgacggagaccgccgtcgtcgacaagccGCTGCAGAAGGGATTCC | 0.67 % |
| | GGGGTCCCAGGTGCTGAatgatcctgacgacggagaccgccgtcgtcgacaa---GCTGCAGAAGGGATTCC | 0.62 % |
|  | GGGGTCCCAGGTGCTGAatgatcctg------------------------------GCTGCAGAAGGGATTCC | 0.55 % |
| | GGGGTCCCAGGTGCTGAatgatcctgacgacggagaccgccgtcgtcgacaagccAGCTGCAGAAGGGATTCC | 0.49 % |

#### *FANCF* / SpCas9(H840A)-RT / Homology arm (*attB* insertion)

Insertion
Subtitution
Deletion : -----

| | | |
|---|---|---|
| Precise insertion | GGGGTCCCAGGTGCTGAatgatcctgacgacggagaccgccgtcgtcgacaagccGCTGCAGAAGGGATTCC<br>*attB* (38bp) | 7.61 % |
| Pure indel | GGGGTCCCAGGTGCTGAAGTA-GTAGTGCTTGAGACCGCCAGAAGCTCGGAAAAGCGATCCAGGTGCTGCAGAAGGGATTCC | 0.04 % |
|  | GGGGTCCCAGGTGCTGACGTAGGTAGTGCTTGAGACCGCCAGAAGCTCGG----GCGATCCCAGGTGCTGCAGAAGGGATTCC | 0.03 % |
| Partial insertion | GGGGTCCCAGGTGCTGACGTAGGTAGTGCTTGAGACCGCC-------------------------------------------------gtcgtcgacaagccGCTGCAGAAGGGATTCC | 0.77 % |
| | GGGGTCCCAGGTGCTGAGtgatcctgacgacggagaccgccgtcgtcgacaagccGCTGCAGAAGGGATTCC | 0.42 % |
| | GGGGTCCCAGGTGCTGAatgatcctgGcgacggagaccgccgtcgtcgacaagccGCTGCAGAAGGGATTCC | 0.40 % |
|  | GGGGTCCCAGGTGCTGACGTAGGTAGTGCTTGAGACCGC----------------------------------------gagaccgccgtcgtcgacaagccGCTGCAGAAGGGATTCC | 0.33 % |
| | GGGGTCCCAGGTGCTGAatgGtcctgacgacggagaccgccgtcgtcgacaagccGCTGCAGAAGGGATTCC | 0.23 % |

Insertion
Subtitution
Deletion : -----

#### *FANCF* / La-SpCas9(H840A)-RT / Homology arm (*attB* insertion)

| | Sequence | % |
|---|---|---|
| Precise insertion | GGGGTCCCAGGTGCTGAatgatcctgacgacggagaccgccgtcgtcgacaagccGCTGCAGAAGGGATTCC<br>*attB* (38bp) | 9.87 % |
| Pure indel | GGGGTCCCAGGTGCTGACGTAGGTAGTGC------------------------GCGATCCCAGGTGCTGCAGAAGGGATTCC | 0.04 % |
|  | GGGGTCCCAGGTGCTGA-----------------------------------------GGCTCCAGGTGCTGCAGAAGGGATTCC | 0.01 % |
| Partial insertion | GGGGTCCCAGGTGCTGAatgatcctgacgacggagaccgccgtcgtcgacaGgccGCTGCAGAAGGGATTCC | 0.99 % |
| | GGGGTCCCAGGTGCTGAatgatccCgacgacggagaccgccgtcgtcgacaagccGCTGCAGAAGGGATTCC | 0.75 % |
| | GGGGTCCCAGGTGCTGAGtgatcctgacgacggagaccgccgtcgtcgacaagccGCTGCAGAAGGGATTCC | 0.34 % |
| | GGGGTCCCAGGTGCTGAatgGtcctgacgacggagaccgccgtcgtcgacaagccGCTGCAGAAGGGATTCC | 0.31 % |
| | GGGGTCCCAGGTGCTGAatgatcctgGcgacggagaccgccgtcgtcgacaagccGCTGCAGAAGGGATTCC | 0.28 % |

Insertion
Subtitution
Deletion : -----

#### *FANCF* / SpCas9(WT)-RT / Homology arm (*attB* insertion)

| | Sequence | % |
|---|---|---|
| Precise insertion | GGGGTCCCAGGTGCTGAatgatcctgacgacggagaccgccgtcgtcgacaagccGCTGCAGAAGGGATTCC | 2.72 % |
| Pure indel | GGGGTCCCAGGTGCTGAC--------------------------------------------GCTGCAGAAGGGATTCC | 63.87 % |
|  | GGGGTCCCAGGTGCTGA---------------------------------------------GCTGCAGAAGGGATTCC | 1.23 % |
|  | G----------------------------GACCGCCAGAAGCTCGGAAAAGCGATCCAGGTGCTGCAGAAGGGATTCC | 0.70 % |
| | GGGGTCCCAGGTGCTGAATGCGTAGGTAGTGCTTGAGACCGCCAGAAGCTCGGAAAAGCGATCCAGGTGCTGCAGAAGGGATTCC | 0.27 % |
| | GGGGTCCCAGGTGCTGAATCGTAGGTAGTGCTTGAGACCGCCAGAAGCTCGGAAAAGCGATCCAGGTGCTGCAGAAGGGATTCC | 0.25 % |
| Partial insertion | GGGGTCCCAGGTGCTGAatgatcctgac--------------------------GCTGCAGAAGGGATTCC | 0.33 % |
| | GGGGTCCCAGGTGCTGAatgatcctgacgacggagaccgccgtcgtcgacaagccCGCTGGCTGCAGAAGGGATTCC | 0.23 % |
|  | GGGGTCCCAGGTGCTGAatgatcc------------------------------GCTGCAGAAGGGATTCC | 0.23 % |
|  | GGGGTCCCAGGTGCTGACGTAGGTAGTGCTTGAGACCGCC---------------------------------------------gtcgtcgacaagccGCTGCAGAAGGGATTCC | 0.16 % |
| | GGGGTCCCAGGTGCTGAatgatcctgacgacggagaccgccgtcgtcgacaagccGCTGCAGAAGGGATTCGTAGGTAGTGCTTGAGACCGCCAGAAGCTCGGAAAAGCGATCCAGGTGCTGCAGAAGGGATTCC | 0.16 % |

Insertion
Subtitution
Deletion : -----

#### *FANCF* / La-SpCas9(WT)-RT / Homology arm (*attB* insertion)

| | Sequence | % |
|---|---|---|
| Precise insertion | GGGGTCCCAGGTGCTGAatgatcctgacgacggagaccgccgtcgtcgacaagccGCTGCAGAAGGGATTCC | 2.21 % |
| Pure indel | GGGGTCCCAGGTGCTGA---------------------------------------------GCTGCAGAAGGGATTCC | 46.04 % |
|  | GGGGTCCCAGGTGCTGAC--------------------------------------------GCTGCAGAAGGGATTCC | 0.55 % |
|  | GGGGTCCCAGGTGCTGAGTGAG---------------------------------------------GCTGCAGAAGGGATTCC | 0.52 % |
|  | GGGGTCCCAGGTGCTGAGCC---------------------------------------------GCTGCAGAAGGGATTCC | 0.49 % |
|  | GGGGTCCCAGGTGCTGAA---------------------------------------------GCTGCAGAAGGGATTCC | 0.42 % |
| Partial insertion | GGGGTCCCAGGTGCTGAatgatcctgacgacggagaccgccgtcgtcgacaagccGCTGCAGAAGGGATTGCTGCAGAAGGGATTCC | 0.45 % |
| | GGGGTCCCAGGTGCTGAatgatcctgacgacggagaccgccgtcgtcgacaagccGCTGCAGAAGCTGCAGAAGGGATTCC | 0.24 % |
|  | GGGGTCCCAGGTGCTGAatgatcctgacgac-----------------------GCTGCAGAAGGGATTCC | 0.22 % |
| | GGGGTCCCAGGTGCTGAatgatcctgacgacggagaccgccgtcgtcgacaagccGCTGCAGCTGCAGAAGGGATTCC | 0.15 % |
|  | GGGGTCCCAGGTGCTGACGTAGGTAGTGCTTGAGACCGCC---------------------------------------------gtcgtcgacaagccGCTGCAGAAGGGATTCC | 0.14 % |

Insertion
Subtitution
Deletion : -----

#### *FANCF* / SpCas9(H840A)-RT / Partial insertion (*attB* insertion)

| | Sequence | % |
|---|---|---|
| Precise insertion | GGGGTCCCAGGTGCTGAatgatcctgacgacggagaccgccgtcgtcgacaagccGCTGCAGAAGGGATTCC | 18.33 % |
| Pure indel | None | |
| Partial insertion | GGGGTCCCAGGTGCTGACGTAGGTAGTGCTTGAGACCGCC---------------------------------------------gtcgtcgacaagccGCTGCAGAAGGGATTCC | 1.31 % |
|  | GGGGTCCCAGGTGCTGAatgatcctgacgacggagac----------------------------cgacggagaccgccgtcgtcgacaagccGCTGCAGAAGGGATTCC<br>Sense insertion / Anti-Sense insertion | 0.51 % |
| | GGGGTCCCAGGTGCTGAatgatcctgacgacggagaccgcCGTcgtcgtcgacaagccGCTGCAGAAGGGATTCC | 0.44 % |
| | GGGGTCCCAGGTGCTGAatgatcct---gacggagaccgccgtcgtcgacaagccGCTGCAGAAGGGATTCC | 0.40 % |
|  | GGGGTCCCAGGTGCTGAatgatcctgacgacggagaccgccgtG------------------------cggagaccgccgtcgtcgacaagccGCTGCAGAAGGGATTCC<br>Sense insertion / Anti-Sense insertion | 0.35 % |

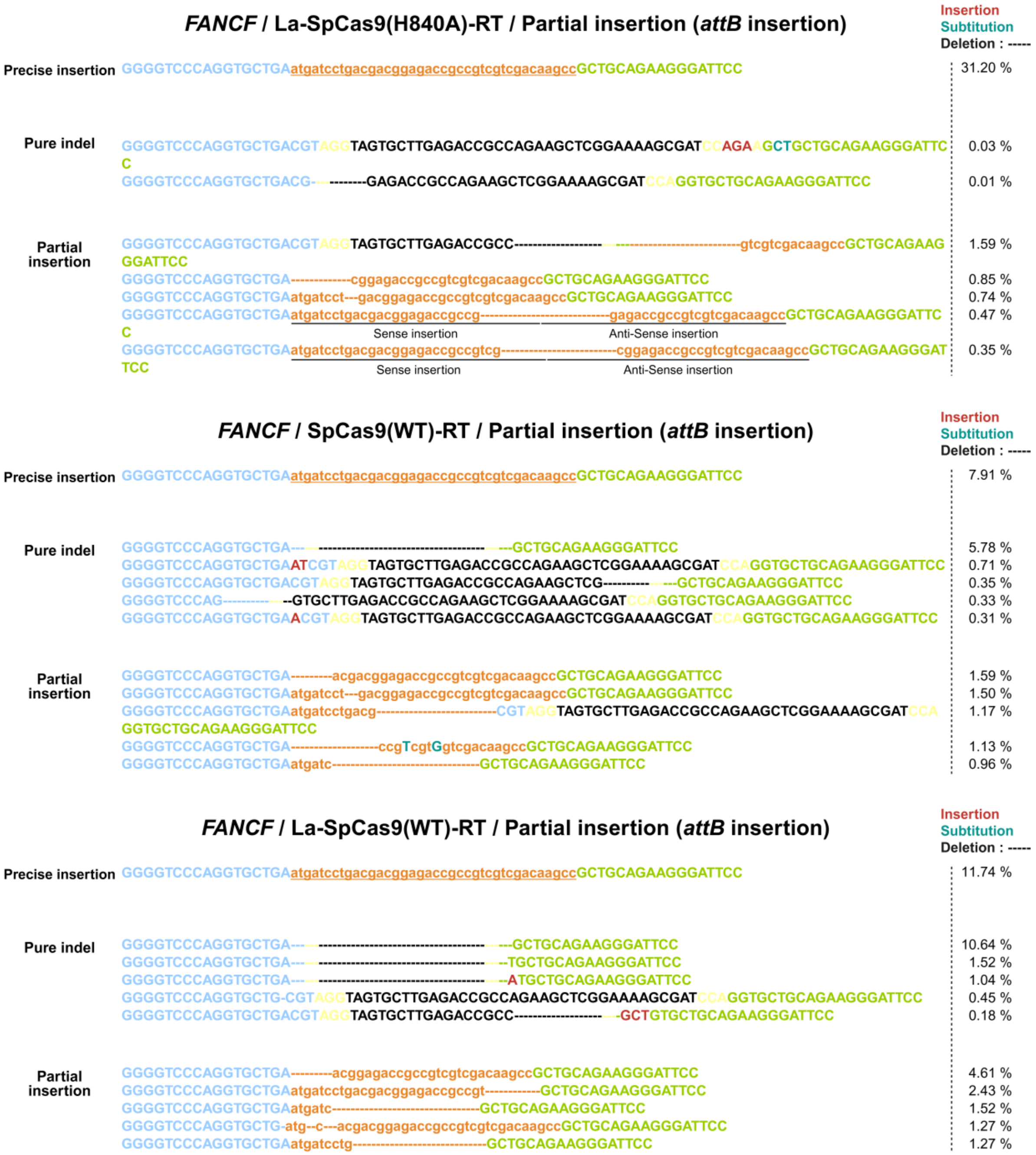


**Supplementary Figure S3. Commonly edited target sequences and corresponding NGS analysis results mediated by SpCas9(H840A/WT)-RT or La-SpCas9(H840A/WT)-RT.** For each gene, the target sequences of the pegRNA pairs used with SpCas9(H840A/WT)-RT or La-SpCas9(H840A/WT)-RT, along with their respective PAM (NGG) sequences, are shown. Within the target DNA sequences, the protospacers and PAM (NGG) sequences corresponding to each sense and antisense pegRNA are colored in light blue, light green, and yellow, respectively. In the edited products, the frequencies (%) of precise insertions, indels (insertions/deletions), and substitutions, as determined by NGS analysis, are indicated to the right of each sequence. In the sequencing results, red nucleotides represent insertions, dashed lines indicate deletions, and blue nucleotides denote substitutions. Orange highlights correspond to precisely or partially inserted *attB*/*attP* sequences.

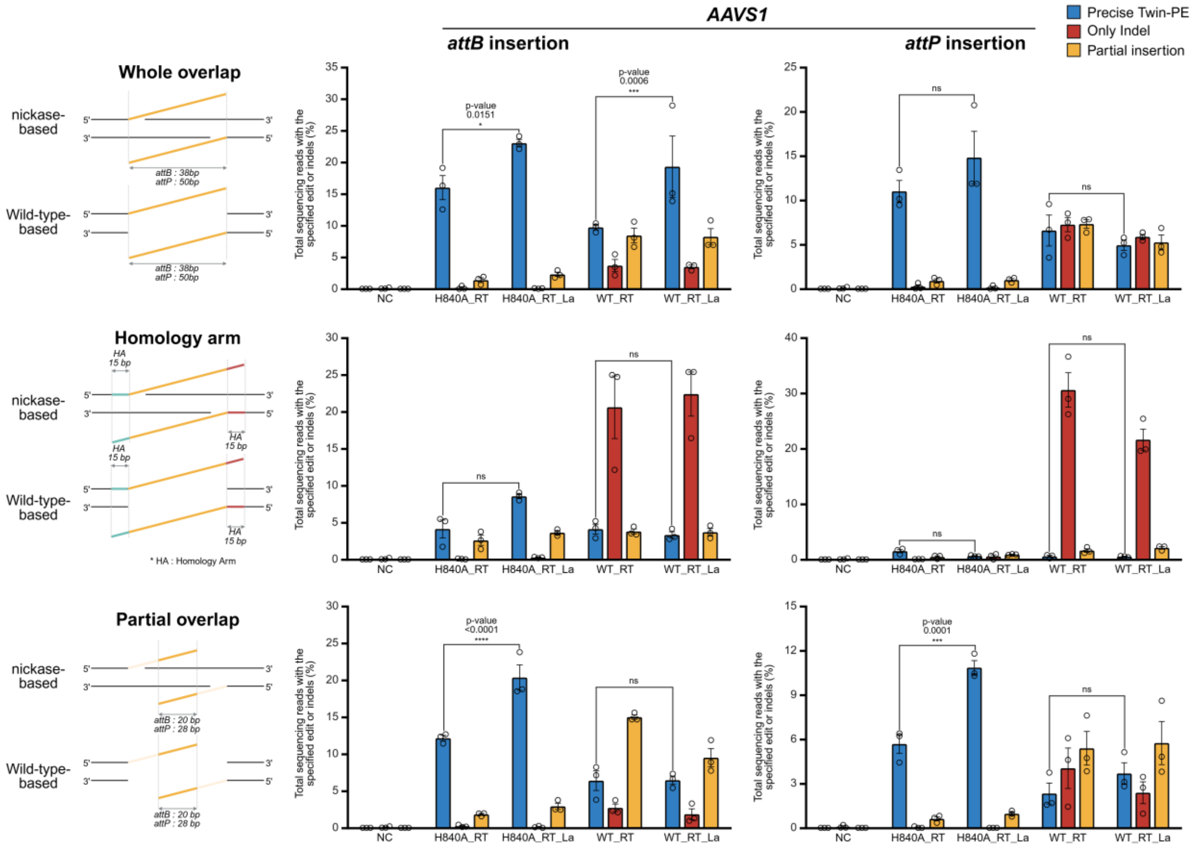


**Supplementary Figure S4. Optimization of twin prime editing components and insertion of various sequences (*attB*/*attP*) at the *AAVS1* locus in human-derived cells.** Comparison of twin prime editing efficiency (%) for insertion of a 38 bp *attB* or a 50 bp *attP* Bxb1 recognition sequence at the *AAVS1* locus in human-derived cells (HEK293FT) using various pegRNA overlap strategies. A comparative analysis was conducted using prime editors based on either nickase or wild-type SpCas9 (SpCas9(H840A/WT)-RT), and those fused with the La domain (La-SpCas9(H840A/WT)-RT). The pegRNA designs were categorized into three types: (i) Whole overlap pairs, in which the entire insertion sequence is fully complementary in both pegRNAs; (ii) Homology arm pairs, where each pegRNA includes the intended insertion sequence along with flanking homology arms; and (iii) Partial overlap pairs, which share only a 20 bp segment of the insertion sequence between the two pegRNAs. Editing outcomes were categorized as precise Twin-PE (%), only indel (%), and partial insertion (%), **(Supplementary Figure S3)**. Each histogram represents mean ± SEM from three independent experiments. P-values were calculated by two-way ANOVA and Dunnett's test (ns: not significant, *P = 0.0332, **P = 0.0021, ***P = 0.0002, ****P < 0.0001). RT: reverse transcriptase, La: La domain, NC: negative control, H840A_RT: SpCas9(H840A)-RT, H840A_RT_La: La-SpCas9(H840A)-RT, WT_RT: SpCas9(WT)-RT, WT_RT_La: La-SpCas9(WT)-RT.

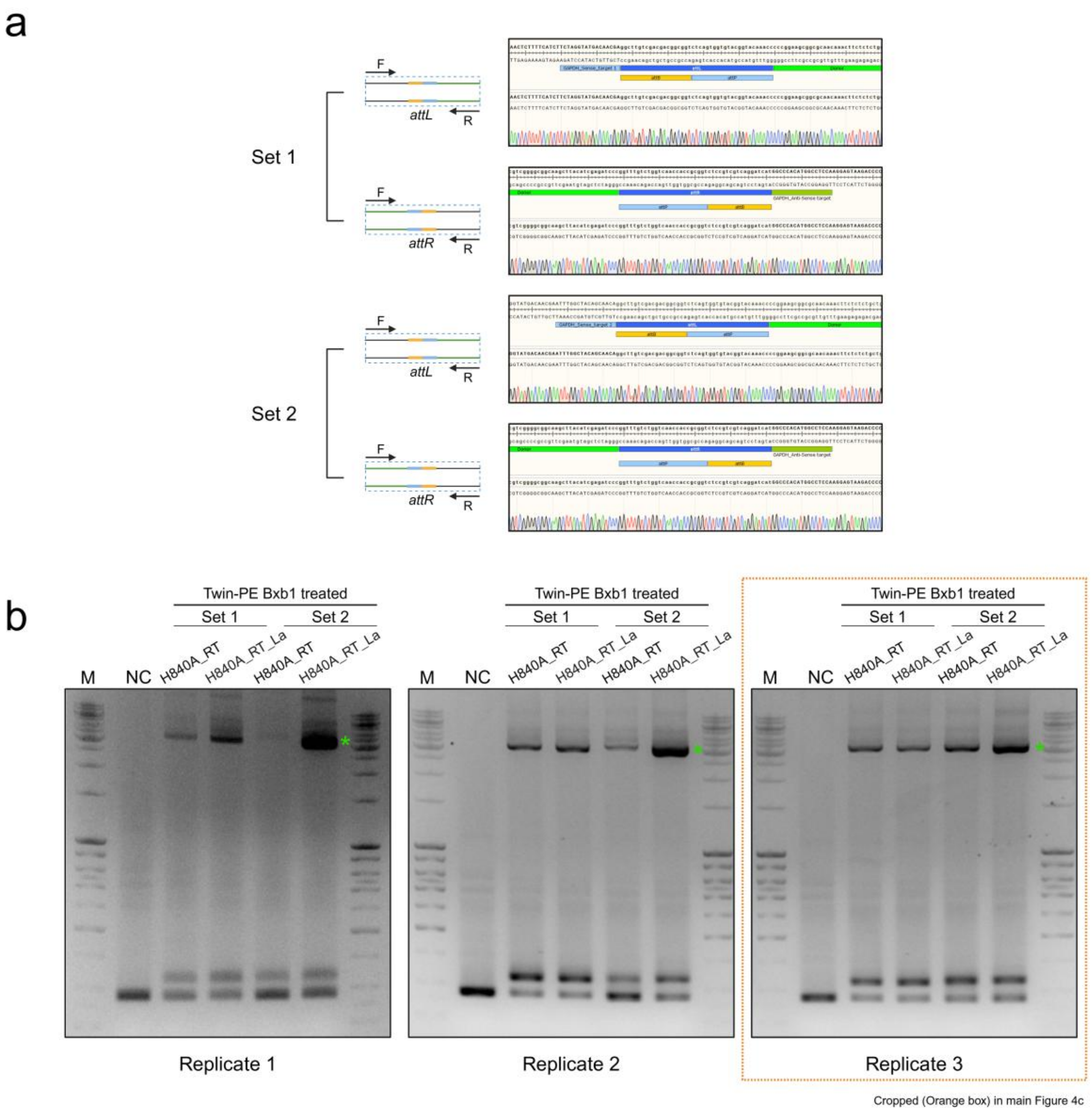


**Supplementary Figure S5. Validation of large gene insertion using optimized twin prime editing in human-derived cells.** (a) Sanger sequencing results from the top-shifted bands in Figure 4, confirming precise donor DNA insertion. The recombined *attL* and *attR* sequences generated from *attB-attP* recombination at the *GAPDH* locus were identified and categorized according to pegRNA pairs (Set1, Set2). (b) Comparison of donor DNA insertion efficiency mediated by twin prime editing using SpCas9(H840A)-RT or La-SpCas9(H840A)-RT, followed by sequential action of the Bxb1 recombinase. Shown are 1.5% agarose gel images displaying PCR-amplified products from the *GAPDH* locus in triplicate experiments, in which cells were treated with each prime editor and the Bxb1 enzyme. Top band: Fragment containing inserted donor DNA. Middle band: Fragment with *attB* insertion only via twin prime editing.

Bottom band: Wild-type DNA fragment without editing. The data highlighted in orange box is presented in Figure 4.

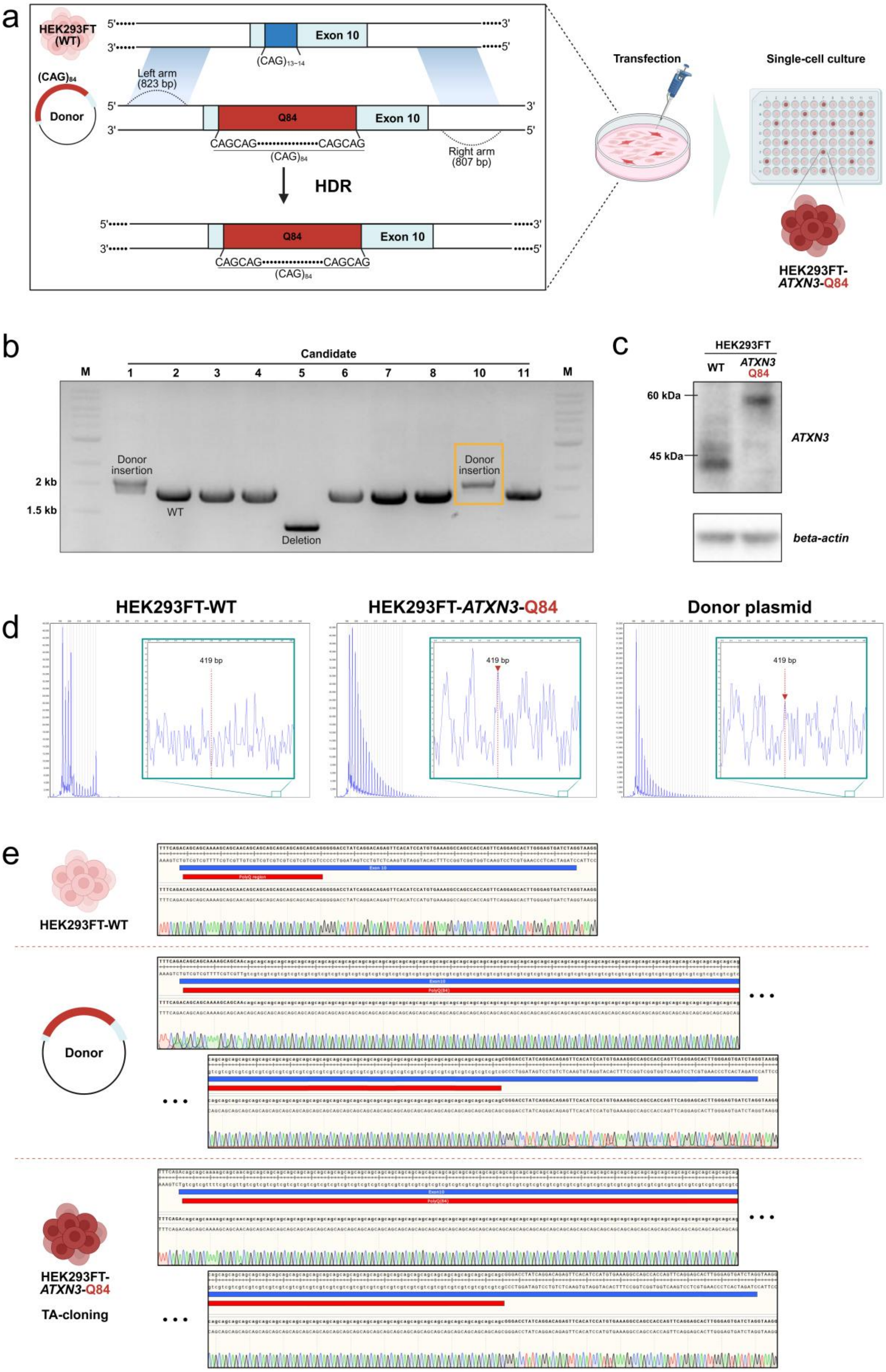
a
HEK293FT (WT)
Exon 10
(CAG)13~14
(CAG)84
Donor
Left arm (823 bp)
Q84
Exon 10
CAGCAG••••••••••••••••CAGCAG
(CAG)84
Right arm (807 bp)
HDR
Transfection
Single-cell culture
HEK293FT-ATXN3-Q84
b
Candidate
M 1 2 3 4 5 6 7 8 10 11 M
Donor insertion
2 kb
1.5 kb
WT
Deletion
c
HEK293FT
WT
ATXN3 Q84
60 kDa
45 kDa
ATXN3
beta-actin
d
HEK293FT-WT
HEK293FT-ATXN3-Q84
Donor plasmid
419 bp
e
HEK293FT-WT
Donor
HEK293FT-ATXN3-Q84
TA-cloning

**Supplementary Figure S6. Construction and validation of a human-derived mutant HEK293FT-*ATXN3*-Q84 cell line.** (a) Schematic of HDR strategy for inserting 84 CAG repeats into exon 10 of the *ATXN3* gene in HEK293FT cells to generate an SCA3 patient-mimicking mutant cell line. Donor DNA containing the 84 CAG repeats and left/right homology arms was co-delivered with a SpCas9 or sgRNA expression vector. (b) Gel image showing *ATXN3* PCR amplicons from single-cell clones. Only clones with precise 84 CAG repeat insertion at exon 10 were selected for subculture. Clone in lane 10 (orange box) was selected as the mutant HEK293FT-*ATXN3*-Q84 cell line. (c) Western blot data comparing *ATXN3* protein expression between wild-type and mutant HEK293FT-*ATXN3*-Q84 cell lines using anti-*ATXN3* and anti-*beta-actin* antibodies. (d) TP-PCR analysis comparing CAG repeat expansion in wild-type and mutant HEK293FT cells. PCR results were compared with those of the donor plasmid. (e) Sequencing of the polyQ region in wild-type HEK293FT, donor plasmid, and mutant HEK293FT-*ATXN3*-Q84 cells confirmed the precise 84 CAG repeat insertion in exon 10 of *ATXN3*.

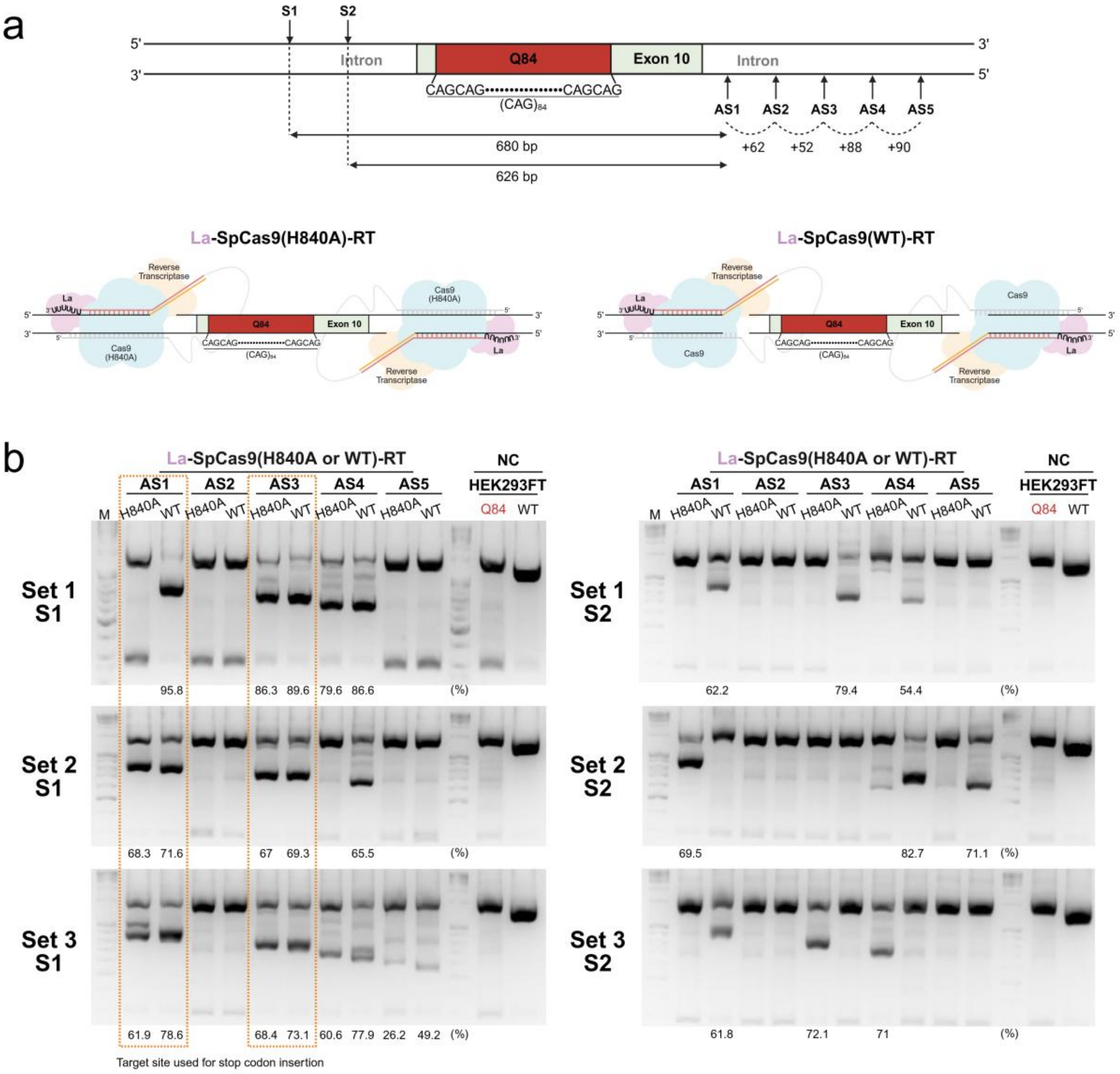


**Supplementary Figure S7. Selection of targeting sites for efficient twin prime editing in human-derived mutant HEK293FT-*ATXN3*-Q84 cells.** (a) Schematic of targeting designs for pegRNA pairs (S1+AS1~AS5, S2+AS1~AS5) in upstream and downstream regions of *ATXN3* exon 10 in the mutant HEK293FT-*ATXN3*-Q84 cell line. Candidate pegRNA combinations using La-SpCas9(H840A)-RT or La-SpCas9(WT)-RT were tested to induce high-efficiency twin prime editing around the polyQ repeats. (b) Gel image comparing PCR amplicons from the *ATXN3* exon 10 region after applying various pegRNA combinations. Lower bands indicate *attB* sequence insertion or simple deletion of exon 10. Orange boxes indicate pegRNA pairs (S1-AS1, S1-AS3) selected for high-efficiency twin prime editing.

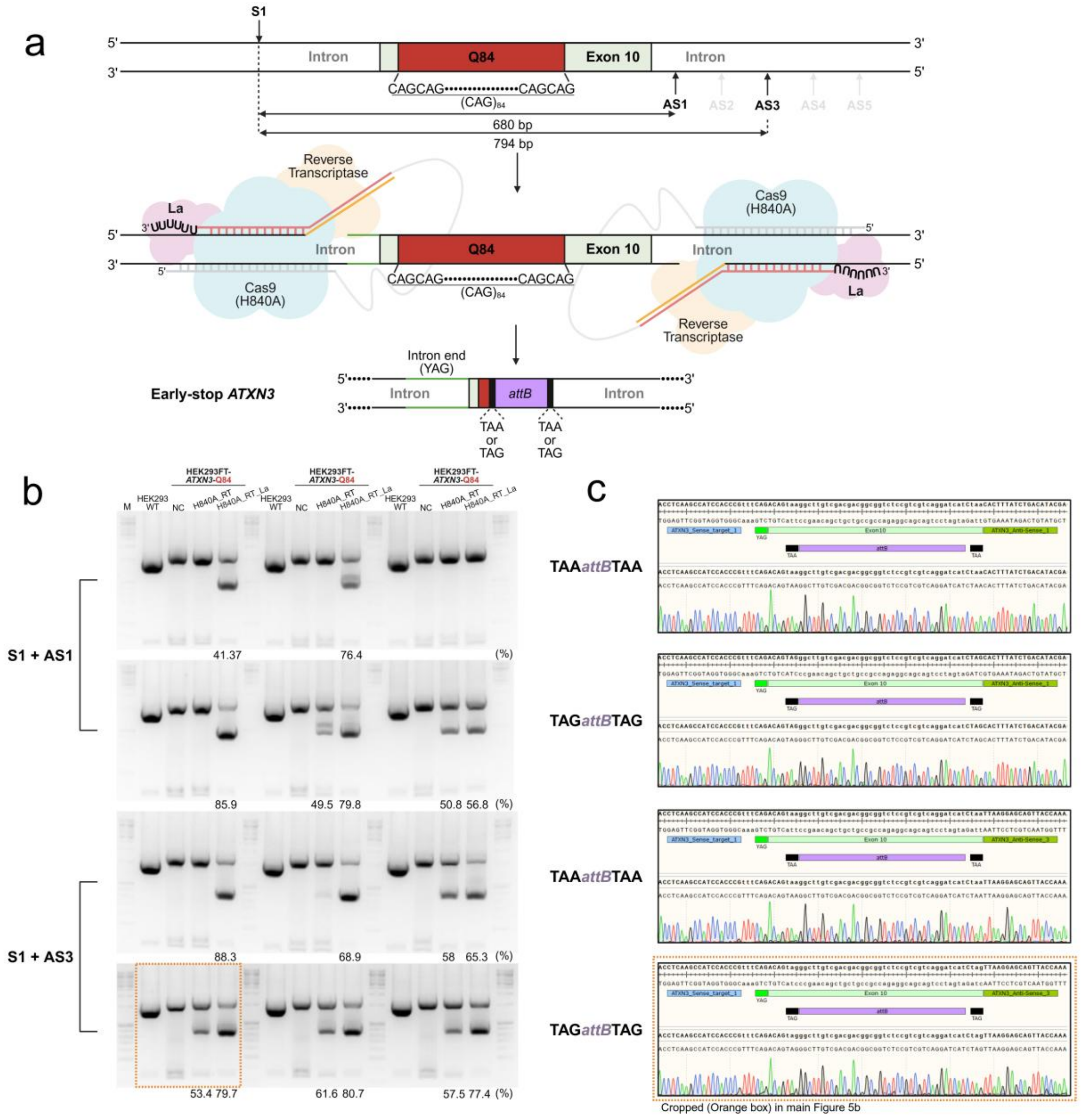

a
S1
Intron
Q84
Exon 10
CAGCAG••••••••••••••••CAGCAG
(CAG)84
AS1 AS2 AS3 AS4 AS5
680 bp
794 bp
Reverse Transcriptase
La
Cas9 (H840A)
Intron end (YAG)
Early-stop ATXN3
attB
TAA or TAG
b
HEK293FT-ATXN3-Q84
M
HEK293 WT
NC
H840A_RT
H840A_RT_La
S1 + AS1
S1 + AS3
41.37
76.4
85.9
49.5 79.8
50.8 56.8
88.3
68.9
58 65.3
53.4 79.7
61.6 80.7
57.5 77.4
(%)
c
TAAattBTAA
TAGattBTAG
Cropped (Orange box) in main Figure 5b

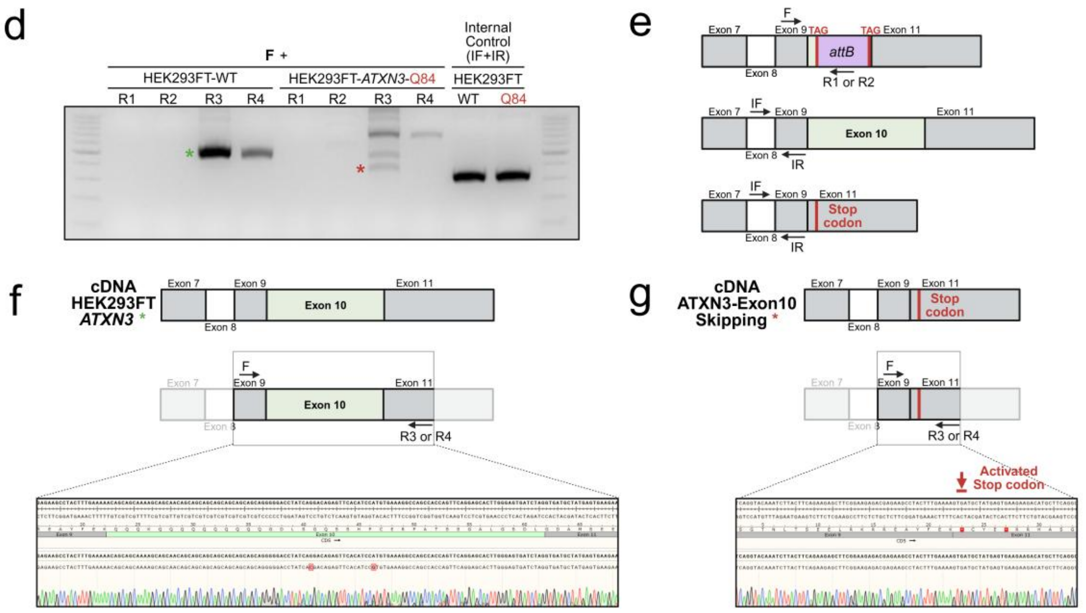


**Supplementary Figure S8. Verification of polyQ removal and early stop codon insertion using optimized twin prime editing in human-derived mutant HEK293FT-*ATXN3*-Q84 cells.** (a) Schematic diagram illustrating the strategy for insertion of an early stop codon (TAA or TAG) and removal of polyQ repeats in the *ATXN3* gene of a human-derived SCA3 disease model cell line using twin prime editing. (b) Efficiency analysis of polyQ removal and early stop codon insertion by twin prime editing targeting *ATXN3* exon 10 using SpCas9(H840A)-RT or La-SpCas9(H840A)-RT. Lower gel bands represent deletion of exon 10 and stop codon insertion. The best pegRNA combinations (S1+AS1 (top), S2+AS3 (bottom)) from Figure S7 were used. Stop codon insertion efficiency (%) was calculated as size-normalized bottom band intensity / total band intensity. The data highlighted in orange box is presented in Figure 5. (c) Sanger sequencing results confirming correct stop codon insertion from the shifted bands in (b). (d) Representative gel image illustrating the comparative analysis of cDNA synthesized from transcriptomes derived from wild-type HEK293FT cells and La-SpCas9(H840A)-RT treated mutant HEK293FT-*ATXN3*-Q84 cells. The green and red asterisks denote the cDNA amplicons derived from wild-type HEK293FT cells and La-SpCas9(H840A)-RT-treated mutant HEK293FT-*ATXN3*-Q84 cells, respectively. F and R (R1–R4) indicate the combinations of PCR primers used for cDNA sequencing, where F represents the forward primer and R indicates the reverse primers. IF and IR denote the forward and reverse primers, respectively, used for internal control amplification. (e) Each schematic illustrates the binding positions of

DNA primers targeting specific regions of the expected cDNA species shown in panel (d). (f–g) PolyQ repeat sequencing results of *ATXN3* cDNA (green and red asterisk) synthesized from transcriptomes derived from wild-type HEK293FT cells (f) and mutant HEK293FT-*ATXN3*-Q84 cells treated with La-SpCas9(H840A)-RT (g). The DNA sequencing results show the junction regions of Exon 9–Exon 10, Exon 10–Exon 11 and Exon 9–Exon 11 within the cDNA, respectively.
